# Stably expressed genes in single-cell RNA-sequencing

**DOI:** 10.1101/475426

**Authors:** Julie M. Deeke, Johann A. Gagnon-Bartsch

**Affiliations:** Department of Statistics, University of Michigan, Ann Arbor

## Abstract

**Motivation:** In single-cell RNA-sequencing (scRNA-seq) experiments, RNA transcripts are extracted and measured from isolated cells to understand gene expression at the cellular level. Measurements from this technology are affected by many technical artifacts, including batch effects. In analogous bulk gene expression experiments, external references, e.g., synthetic gene spike-ins often from the External RNA Controls Consortium (ERCC), may be incorporated to the experimental protocol for use in adjusting measurements for technical artifacts. In scRNA-seq experiments, the use of external spike-ins is controversial due to dissimilarities with endogenous genes and uncertainty about sufficient precision of their introduction. Instead, endogenous genes with highly stable expression could be used as references within scRNA-seq to help normalize the data. First, however, a specific notion of stable expression at the single cell level needs to be formulated; genes could be stable in absolute expression, in proportion to cell volume, or in proportion to total gene expression. Different types of stable genes will be useful for different normalizations and will need different methods for discovery.

**Results:** We compile gene sets whose products are associated with cellular structures and record these gene sets for future reuse and analysis. We find that genes whose final product are associated with the cytosolic ribosome have expressions that are highly stable with respect to the total RNA content. Notably, these genes appear to be stable in bulk measurements as well.

**Supplementary information:** The Supplement is available on bioRxiv, and the gene set database is available through GitHub.

## 1 Introduction

Single-cell RNA-sequencing (scRNA-seq) experiments measure gene expression at the cellular level, capturing details at a resolution previously not possible. However, challenges arise due to unwanted variation that scRNA-seq experiences. Some sources of unwanted variation include read depth, capture efficiency, amplification biases, batch effects, and cell cycle [Hicks *et al.*, 2018, Phipson *et al.*, 2017, Lun and Marioni, 2017, Dabney and Meyer, 2012, Kolodziejczyk *et al.*, 2015]. Methods have been developed to remove some sources of unwanted variation, often using certain sets of reference genes to aid in removing either specific or general effects. For example, Chen and Zhou [2017] use Bayesian methods to identify control genes that are unassociated with a factor of interest and adjust the target genes based on the control genes. Buettner *et al.* [2015] uses genes that have been annotated as associated with the cell cycle to remove cell cycle effects from the data. Both Brennecke *et al.* [2013] and Grün *et al.* [2014] propose using external spike-in references to remove some of the technical noise present in the data. Finally, Lin *et al.* [2018] propose using stably expressed genes as a form of negative controls to remove unwanted variation using a procedure called scMerge.

Commercially generated, synthetic, external spike-in references (often External RNA Controls Consortium (ERCC) spike-ins) can be used in bulk gene expression experiments [Baker *et al.*, 2005, Jiang *et al.*, 2011, Risso *et al.*, 2014, Pine *et al.*, 2016]. The ability to incorporate external spike-in references in scRNA-seq varies based on the cell isolation protocol. Spike-ins are not typically included in droplet-based isolation protocols but can be in plate- and well-based isolation methods including the Fluidigm C1 system [Macosko *et al.*, 2015, Bacher and Kendziorski, 2016, Lun *et al.*, 2016, Tung *et al.*, 2017].

Limitations to the use of spike-ins are not limited to scRNA-seq contexts. A critique in both single cell and bulk experiments is that the spike-ins possess qualities that are dissimilar to endogenous genes [Grün and vanOudenaarden, 2015, Tung *et al.*, 2017]. Spike-ins are designed to exhibit artificially wide ranging characteristics, like length and proportion of guanine and cytosine bases in the nucleic acid sequence, in order to understand how these characteristics might affect downstream results. Specific to scRNA-seq, the quantity of solution added for each cell is much smaller, so minor pipetting errors (with the Fluidigm C1 system) or other volume errors affect results much more than with a larger quantity of solution [Tung *et al.*, 2017]. The technical challenge of accurately introducing and measuring a smaller volume of spike-ins reduces their effectiveness as negative controls in scRNA-seq [Robinson and Oshlack, 2010].

Endogenous genes that are reasonably stably expressed have been proposed for use in normalization of microarray data [Eisenberg and Levanon, 2003, 2013, Gagnon-Bartsch and Speed, 2012]. However, it is unclear that the single cell expression of these same genes exhibit the same stability as in bulk experiments. For example, a gene may be expressed with a bursting mechanism, increasing its variability [Jiang *et al.*, 2017, Suter *et al.*, 2011, Fukaya *et al.*, 2016]. Bulk expression data might identify the gene as stably expressed, but that same classification at the single cell level would be inappropriate.

There is a need to discover single cell-specific stably expressed genes. Lin *et al.* [2017] propose a method of creating an index at the single cell level for generating a set of stably expressed genes across all cell conditions. Desired characteristics of stably expressed genes from Lin *et al.* [2017] include a distribution with a small proportion of measurements with low values and a small variance among the measurements with high values as estimated from parameters of a Gamma-Gaussian model.

The goals of this paper are: (1) to clarify the notion of “stable expression” at the single cell level, and in particular to define multiple such notions, (2) to propose a method in which to identify a set of genes that exhibit stable expression, (3) to organize sets of genes based on the cellular component with which the final gene product is associated, and (4) to suggest the set of cytosolic ribosomal genes as stably expressed with respect to total RNA content.

## 2 Approach

### 2.1 Notions of Stability

We first consider explicitly what it means for a gene to be stable. The idea of stable expression has previously been addressed either implicitly or without much elaboration. However, the notion of stability at the single cell level is inherently ambiguous and requires a precise definition. We consider multiple notions of stability.

One notion of a stably expressed gene would be that the gene is expressed at a constant absolute level. In other words, the number of RNA molecules present within each cell should be approximately constant, e.g., each cell has about 10 RNA molecules of that gene. We refer to this notion of stability as “absolute stability.” Genes that are absolutely stable could replace the external spike-ins, as they are expected to be present at a fixed absolute amount in each cell. Like spike-ins, these genes could be used to pick up certain technical effects, such as reaction efficiency.

A second notion of stable expression would be that genes are expressed at a constant proportion with respect to cell volume; that is, they are stable in terms of concentration. We refer to this as “stable in concentration.” In addition to picking up technical effects, genes that are stable in concentration could also pick up and adjust for effects that are associated with cell size.

Yet another notion of a stably expressed gene would be that the gene is expressed at a constant proportion with respect to the total RNA content of the cell from all genes. We refer to this as “stable in proportion to total RNA content,” or, when it is clear from context, simply as “proportionally stable.” In practice, we expect that sets of genes that are stable in concentration will be similar to sets of genes that are proportionally stable, provided that total RNA scales with cell size; in that case, both are likely to pick up cell size effects.

The notions of stability described above are biological in nature; they make no reference to *measurements* of gene expression, such as those provided by scRNA-seq data. In a hypothetical, extremely high quality dataset, these biological notions of stability would map clearly to features of the data. An absolutely stable gene would have a small standard deviation in terms of raw counts; a proportionally stable gene would have a small standard deviation after dividing the raw counts by total cell count.

Real data, however, is subject to technical factors that strongly affect the observed counts. In particular, some factors, like reaction efficiency, have a strong global effect on all counts for a given cell, effectively introducing a random scaling factor for each cell. Thus in real data, genes that are absolutely stable will not necessarily appear particularly stable in terms of their raw counts.

Normalizing the raw counts by the total cell count can adjust for the random scaling effect, and such normalizations are common (e.g., rpkm is a variant of this). After normalizing by total cell count, however, the notion of stability that is most relevant is proportional stability. That is, genes that appear stable in the normalized data would be those genes that are proportionally stable, and genes that are in truth absolutely stable would not necessarily appear stable in the data. Indeed, for this reason – that normalization by total cell count is effectively necessary to adjust for global technical effects, but that normalization by cell total also obscures absolutely stable expression – absolutely stable genes are especially difficult to identify. Note also that similar comments apply to efforts to discover stably expressed genes in bulk tissue.

Importantly, note that a stably expressed gene can be viewed as the opposite of a differentially expressed gene or a highly variable gene. That is, the notion of stability, whether implicit or explicit, also implies the notion of instability or variability. In practical terms, the normalization that is applied to the data may not simply “clean” the data, but also alter the biological interpretation of the data, and determine which biological questions can (and cannot) be answered by the data. For example, normalizing the data against a set of absolutely stable genes would allow one to identify “absolutely differentially expressed genes,” while normalizing the data against a set of proportionally stable genes would allow one to identify “proportionally differentially expressed genes.” These two sets of differentially expressed genes could be quite different. Thus, finding sets of genes that exhibit different notions of stability would allow for different types of normalization, which provide different biological insights.

### 2.2 Localization of Stable Genes

The final product of a gene (protein, ribosomal RNA, etc.) is often localized to specific structure(s) within the cell, e.g., nucleus, cell membrane, etc. We may therefore associate a gene with the location(s) where that gene’s final product(s) are active.

We hypothesize that certain structures may be enriched with genes that are absolutely stable, while other structures may be enriched with genes that are stable in concentration. For example, because each cell has one nucleus, there may be a set of nuclear genes that are constant in absolute expression. In contrast, there may be a set of genes enriched in the cytosol that are reasonably constant in concentration.

We perform our analysis under the hypothesis that structures are important for identifying stably expressed genes. We therefore create gene sets for each cell structure and assess the sets as a whole.

## 3 Methods

### 3.1 Mapping Genes to Cell Structures

The Gene Ontology Consortium maintains a database that specifies the cellular component(s) with which each gene’s final product is associated [Xin *et al.*, 2016, Wu *et al.*, 2013]. Many of the annotations provided by the Gene Ontology Consortium are highly detailed (e.g., “mitochondrial respiratory chain complex I”); we coarsen these annotations into ten categories corresponding to major cellular structures; see the Supplement for details. The ten categories are: nucleus, endoplasmic reticulum, Golgi body, cytoplasm, membrane, ribosome, mitochondria, mitochondrial ribosome, ribonucleoprotein complex, and cytosolic ribosome. We allow a single gene to be associated with more than one cellular structure, and some genes are not associated with any category. We will refer to these sets of genes as “nuclear genes”, “cytoplasm genes,” etc.

### 3.2 Expression Data

We downloaded six datasets containing human cells processed by three different lab groups and from four different tissue types from Gene Expression Omnibus; see the Supplement for more details [Edgar *et al.*, 2002, Arguel *et al.*, 2017, Das *et al.*, 2017, Tung *et al.*, 2017]. We selected these datasets because they were conducted with the Fluidigm C1 platform and included ERCC spike-ins. Note also that all but one of these datasets make use of unique molecular identifiers (UMIs) [Kivioja *et al.*, 2012]. We filter to the genes that are expressed at least once in all datasets to ensure that genes are expressed in a wide variety of tissue types and to compare the datasets more directly.

### 3.3 Absolute Stability

Genes that are absolutely stable would ideally appear stable in the data in terms of raw counts; however, as noted in Section 2, due to technical factors that result in strong variation in library size from well to well, simply looking for genes that are stable in terms of raw counts is not a feasible way to discover absolutely stable genes. Instead, we leverage the stable absolute “expression” of the ERCC spike-ins, and find absolutely stable genes by looking for those genes that have a high correlation with the ERCCs.

More specifically, for each of the six datasets we perform the following analysis. We begin with raw counts of UMIs. We transform the raw counts by log + 1. In addition, for each cell we also compute the sum of the raw UMIs of all ERCCs; we refer to this as the “ERCC total.” Finally, for each gene, we compute Pearson’s correlation (across all cells) between that gene’s (log + 1) expression and the log of the ERCC total.

We then summarize the correlations by finding, for each gene, the mean of the correlations across the six datasets. Thus, for each gene, we now have an average correlation of that gene’s expression with the ERCC total, and we regard this as a measure of that gene’s absolute stability. To see if any cellular structures are enriched for stably expressed genes, we plot histograms of the correlations subsetted by structure, and inspect these histograms to see if any structures have especially high correlations.

### 3.4 Proportional Stability

We also attempt to find genes that are proportionally stable. Our method is similar to the one for absolute stability but with an adjusted cell total replacing the ERCC total. For a given structure *S*, we sum a cell’s overall measured expression after removing the set of all genes associated with structure *S*; we refer to this as an “adjusted cell total”. For each gene, we compute Pearson’s correlation (across all cells) between that gene’s (log + 1) expression and the log of the adjusted cell total.

Again, we summarize correlations by finding, for each gene, the mean of the correlations across the six datasets. For each structure, we plot a histogram of that structure’s correlations with that structure’s adjusted cell total.

## 4 Results

### 4.1 Absolute Stability

Figure 1 shows a histogram of the correlations with the ERCC total; histograms of correlations separated by gene sets can be found in Supplementary Figures 2 and 3. The most notable observation is that the all correlations were smaller in absolute value than 0.3. The weak correlations indicate that no gene captures the same artifacts as the spike-ins, and vice versa. One possible explanation for this is that the spike-ins are not exposed to some technical effect(s) that endogenous genes are affected by. As cells need to be lysed and RNA needs to be extracted from the cells, we believe that technical factors affect the measurements for endogenous genes while the spike-ins do not experience the same variability.

**Figure 1:**
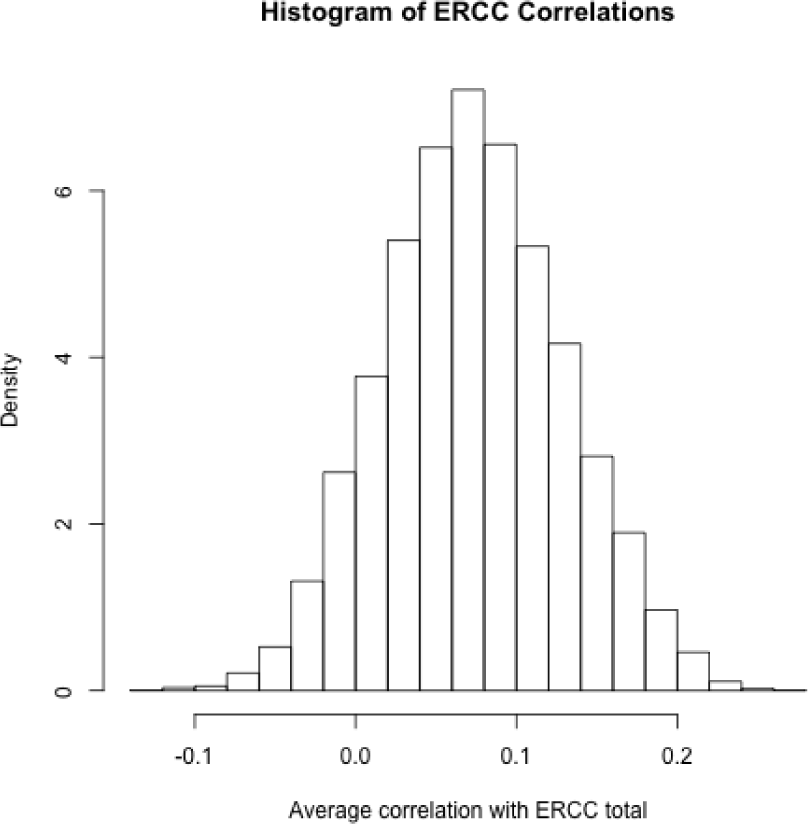
Histogram of the correlations of all genes with the ERCC totals. The supplement contains detailed histograms with separation based on the structure.

The spike-in measurements exhibit lower variability than the biological measurements within our datasets. We compared the biological cell total to the ERCC cell total on a log scale for each of the datasets (Figure 2). We see that the variability for the ERCC measurements is considerably smaller than the variability for the biological total of a cell. Figures examining the mean and standard deviation of each of the genes and ERCC spike-ins separates demonstrate that, again, the ERCC spike-ins have much lower variability when compared to biological genes with similar overall expression (Supplementary Figure 1).

**Figure 2:**
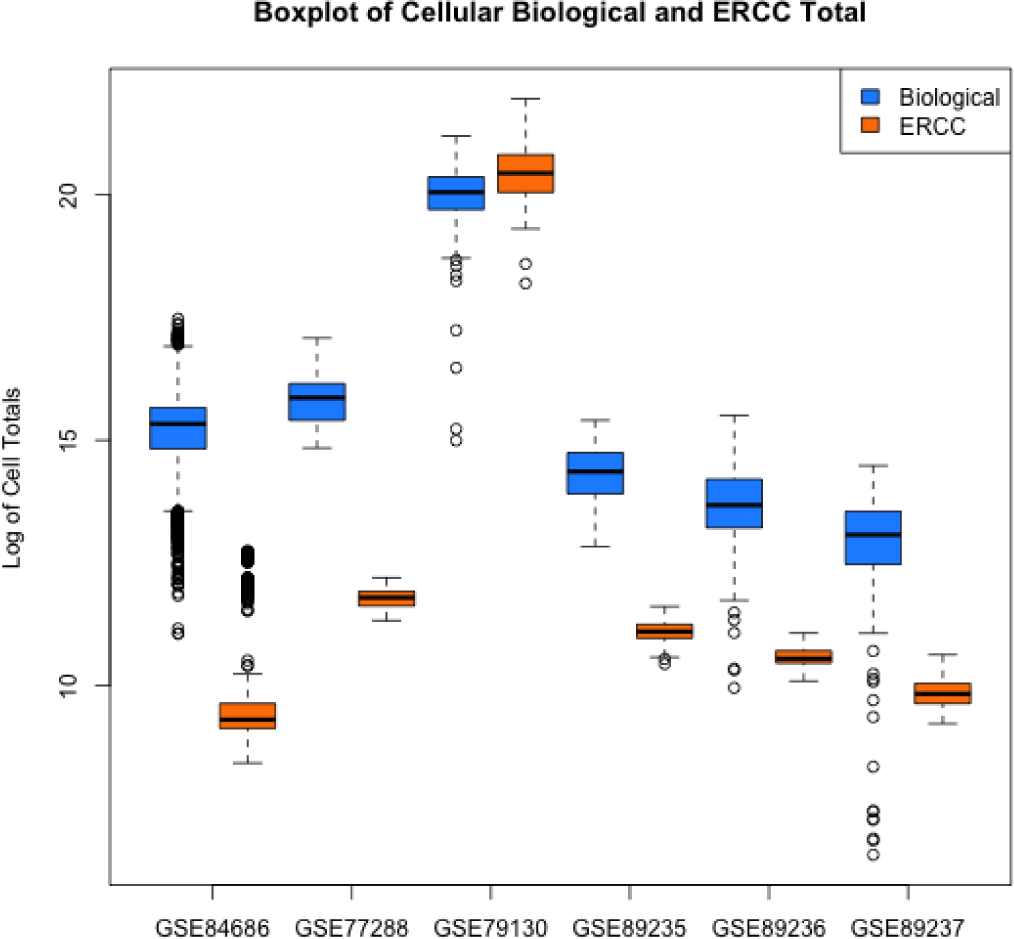
Boxplots of the biological expression of a cell and the ERCC measurements from each cell. The blue boxplots represent the log-transformed sum of the biological expression for each cell, while the orange boxplots are the log-transformed sum of the ERCC measurements for each cell. Separate boxplots are plotted for each dataset.

Overall, the spike-ins appear to have measurements that are more absolutely stable than the endogenous genes. Since each dataset captures one type of cell and Figure 2 displays cell totals, technical effects likely contribute most of the variation. The technical factors affecting the biological cell totals appear larger than technical factors affecting the spike-ins, indicating that they do not appear to be capturing the same technical effects that are affecting the biological measurements. Using spike-ins to identify genes that are absolutely stably expressed is inappropriate.

### 4.2 Proportional Stability

We examine the measures of proportional stability in Figure 3 and Supplementary Figure 4. Unlike the correlations with the ERCC measurements, we see large correlations, with values ranging from −0.03 to 0.95. The cytosolic ribosomal genes exhibit the highest measures of proportional stability based on their correlations.

**Figure 3:**
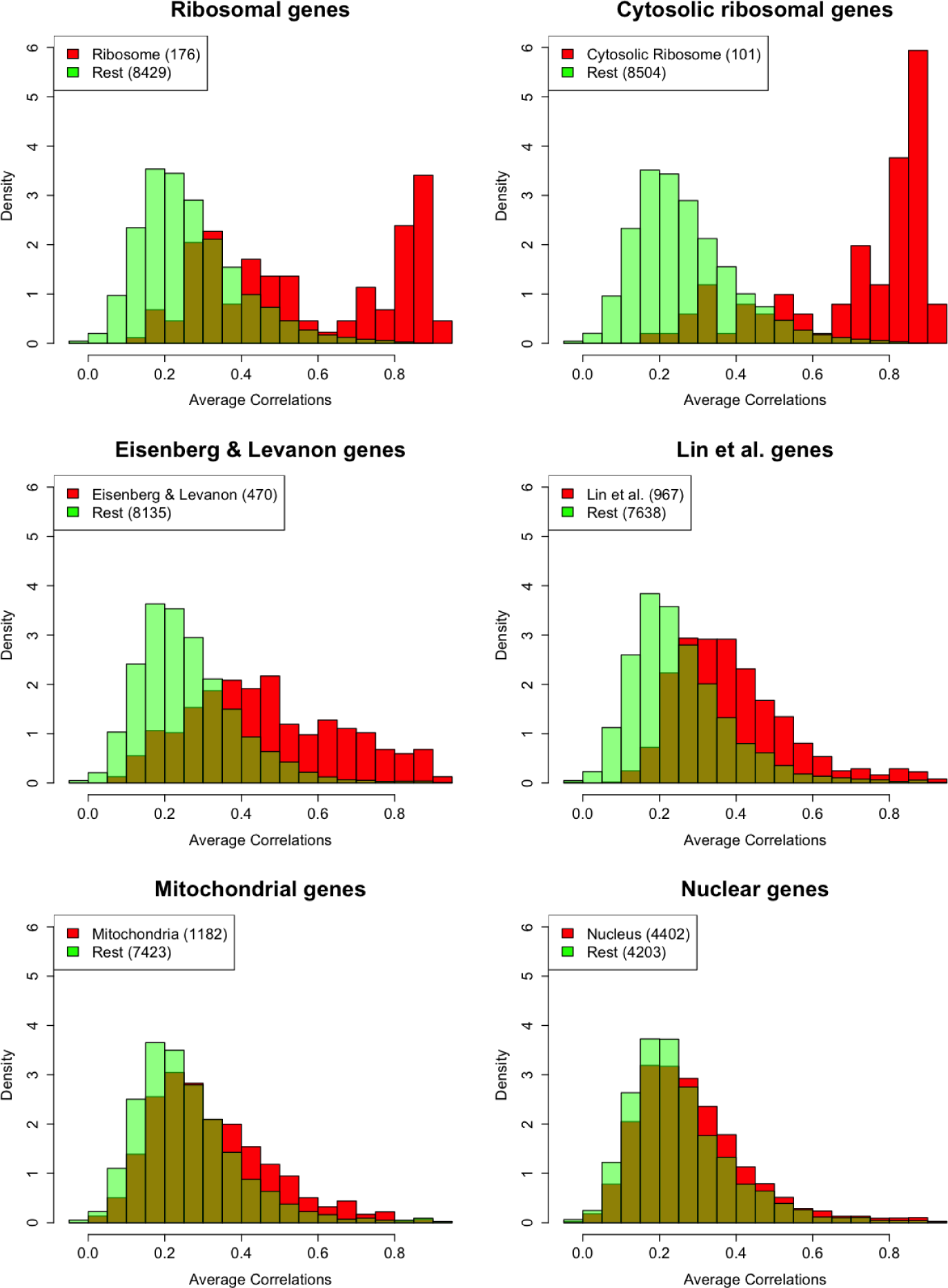
Histograms of the average correlations of each gene over the six datasets considered, with comparisons between the set of genes of interest and the remaining genes. Note that, unlike Figure 1, the x-axis ranges from −0.05 to 0.95. Figures for the additional six strctures can be found in Supplementary Figure 4.

Table 1 displays the distribution of cell structures among the genes in our dataset. We also show the distribution of different subsets of genes, including the genes of Eisenberg and Levanon [2003] and of Lin *et al.* [2017]. Of the genes with the highest average correlations, 44 of the top 45 are cytosolic ribosomal genes. 61 of 101 cytosolic ribosomal genes are in the top 100 genes by correlation; for comparison, 69 of 4,402 nuclear genes, 51 of 470 housekeeping genes from Eisenberg and Levanon [2003], and 37 of 967 single cell housekeeping genes from Lin *et al.* [2017] are in the 100 genes with the highest correlations.

**Table 1:** Distribution of Cell Structures Amongst Different Sets of Genes

| Structure<br>Number of Genes | All Genes<br>8,601 | E & L<br>470 | Lin et al.<br>967 | Top 100<br>100 | Cyto. Ribo.<br>101 |
| --- | --- | --- | --- | --- | --- |
| Nucleus | 51% | 60% | 71% | 69% | 67% |
| Endoplasmic Reticulum | 11% | 14% | 10% | 12% | 8% |
| Golgi Bodies | 10% | 11% | 9% | 1% | 1% |
| Cytoplasm | 30% | 37% | 32% | 37% | 34% |
| Ribosome | 2% | 8% | 8% | 63% | 100% |
| Mitochondria | 14% | 22% | 14% | 20% | 12% |
| Mitochondrial Ribosome | 1% | 1% | 1% | 0% | 4% |
| Ribonucleoprotein | 2% | 5% | 6% | 18% | 18% |
| Membrane | 37% | 51% | 36% | 63% | 63% |
| Cytosolic Ribosome | 1% | 7% | 4% | 61% | 100% |

Ribosomal genes, from which the cytosolic ribosomal genes are a subset, have been previously identified as a plausible option for reference genes for microarray experiments [Thorrez *et al.*, 2008]. Thorrez *et al.* [2008] note that the ribosomal genes do exhibit tissue-dependent variation but are the most stable set of genes that they had encountered.

The expressions of the cytosolic ribosomal genes indicate that they could be effective as reference genes, as most cells express cytosolic ribosomal genes and at a fairly high level (Supplementary Figure 7).

Finally, we compare the correlations of cytosolic ribosomal genes to those of GAPDH and Beta-actin, two genes that have previously been identified as stable genes. Both GAPDH and Beta-actin appear to have similar or smaller correlations as the cytosolic ribosomal genes (on average), indicating that the proportional stability is similar to two commonly used stable genes; the correlation for GAPDH is 0.82 and for Beta-actin is 0.75. For a sense of how these correlations compare to those for the cytosolic ribosomal genes, see Supplementary Figure 6 or examine Panel 2 of Figure 3.

### 4.3 Stability of Cytosolic Ribosomal Genes in Bulk Tissues

For assessment of the stability of cytosolic ribosomal genes in bulk samples from many tissue types, we analyzed data collected with bulk sequencing from the Genotype-Tissue Expression (GTEx) project [Carithers *et al.*, 2015]. The GTEx project systematically collected multiple tissue types from many individuals for genetic profiling and analysis. The tissue types are often subcategorized into additional specific subtissue types.

We examine the stability of different sets of genes by seeing how strongly their expressions vary across tissue type. More specifically, for a given set of genes, we compute the singular-value decomposition (SVD) and plot the first two singular vectors to see how strongly the samples cluster by tissue type. The more pronounced the clustering, the less stable the expression. Prior to calculating the SVD, the GTEx samples are RPKM-normalized, transformed with the log + 1, and centered by gene. In these plots, clustering by tissue type indicates that the genes examined have some differential expression between tissue types, whereas lack of clustering provides evidence of stable expression across tissue types. We examine the SVD plots to assess the stability of the cytosolic ribosomal genes compared to all genes, the Eisenberg and Levanon [2003] genes, and the Lin *et al.* [2017] genes.

We perform the SVDs for both single tissues and combinations of two tissue types. We selected the seven tissue types with the most samples (brain, skin, esophagus, blood vessel, adipose tissue, heart, and muscle); of these, six had subtissue types. We first plotted the SVD for these six tissue types separately with coloring by subtissue type to examine how stably expressed genes are within a tissue type; see Figure 4 and Supplementary Figures 33 to 38. We also plotted SVDs for each combination of two of the seven tissue types to examine how stably expressed genes are across those two tissue types; see Supplementary Figures 39 to 80. Figure 5 shows two sets of two tissue types across which the cytosolic ribosomal genes are especially stable (left panel) and especially not stable (right panel). In general, the cytosolic ribosomal genes exhibit the most stable expression between subtissue types and across different tissue types. However, as also noted by Thorrez *et al.* [2008], the cytosolic ribosomal genes are not highly stable across all tissue types, and thus caution is required when using them for adjustment.

**Figure 4:**
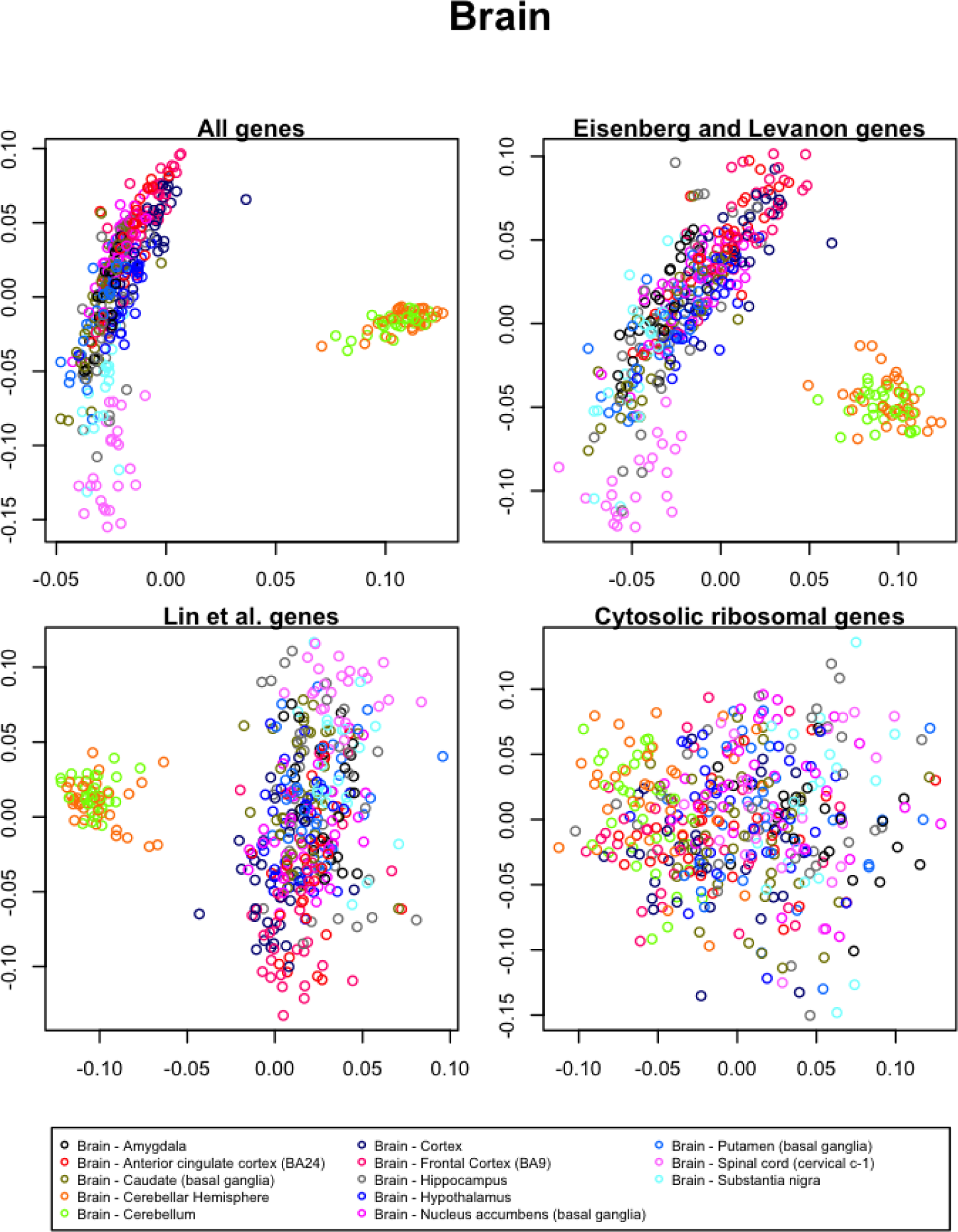
Singular Value Decomposition of GTEx data from the brain using different sets of genes. The overlap of the clusters denoted by the coloring indicate how stably expressed the genes are; the overlap is the strongest for the cytosolic ribosomal genes, indicating that they are the most stably expressed of the three options between the brain subtissue types. Note that the number of genes in each of the four SVD plots differ. More specifically, 89 cytosolic ribosomal genes are present in the available GTEx data, while more are present in each of the other three gene sets. To ensure that differences in the number of features are not accounting for differences in apparent differential expression, we repeat the analysis with random samples of 89 genes from each of the gene sets; Supplementary Figures 81 to 86 show these figures.

**Figure 5:**
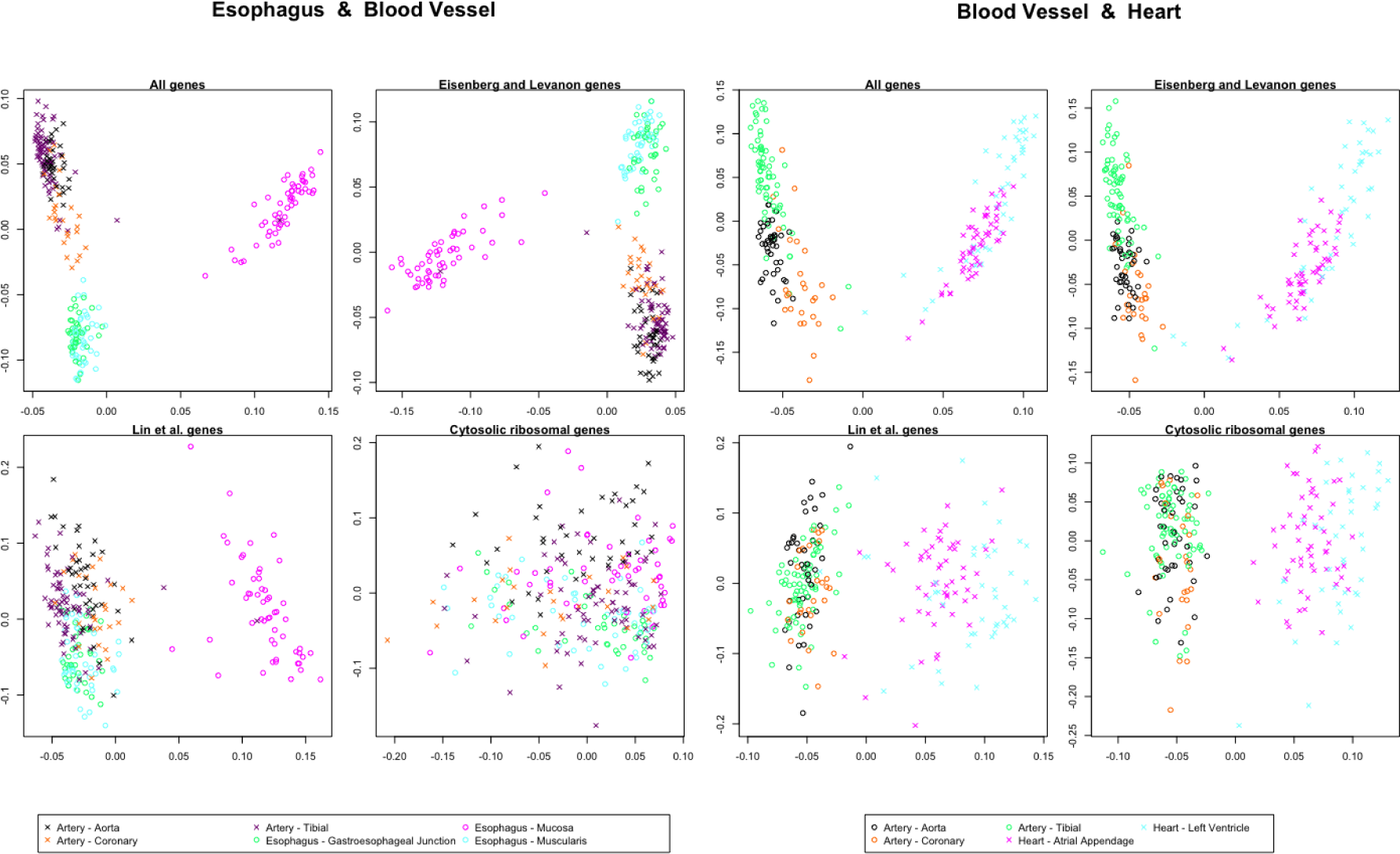
Singular Value Decompositions of GTEx data from the esophagus and blood vessels and blood vessels and heart, using different sets of genes as features. For each set of tissues, the first plot on the top left uses all genes as features; the plot on the top right uses genes proposed by Eisenberg and Levanon [2003] as features; the plot on the bottom left uses genes proposed by Lin *et al.* [2017] as features; and the plot on the bottom right uses cytosolic ribosomal genes as features. The plots display the first two singular vectors. The two tissue types are denoted by either the circle or cross symbol, and the tissue subtypes are denoted by the coloring. The esophagus and blood vessel overlap substantially for the cytosolic ribosomal genes, indicating that their expression is reasonably stable across these tissues and subtissues. Overlap is much smaller for the other three sets of genes, indicating less stable expression of these gene sets. The separation for the blood vessel and heart is clear, indicating that the four sets of genes are not stably expressed across the blood vessel and heart; this separation is one of the strongest that we observed in Supplementary Figures 39 to 80, in which we repeat the analysis for different combinations of tissue types. Again, note that different numbers of genes are used for each of these SVDs. Supplementary Figures 87 to 106 show SVDs for random samples of 89 genes from each gene set.

## 5 Table of Gene Information

We have generated a database that contains summary information about the genes in our datasets. This database contains our structural annotations; means, standard deviations, and correlations from the single cell datasets; and means, standard deviations, and F-statistics for the tissue types from the GTEx data. This information can help customize a set of reference genes specific to task and needs.

Detailed information and code can be found in the supplementary materials for more information and replication of these findings.

## 6 Discussion and Conclusions

Based on our analysis, the cytosolic ribosomal genes appear to be stably expressed proportional to the total RNA content of a cell. The set of cytosolic ribosomal genes have been identified based on high correlations with an adjusted cell total, and thus appear to have good measures of proportional stability. Expression patterns of cytosolic ribosomal genes observed in bulk GTEx experiments further support the conclusion that cytosolic ribosomal genes are proportionally stably expressed.

Proportional stability of the cytosolic ribosomal genes is biologically plausible given their function. The primary function of the ribosome is to translate mRNA transcripts into proteins. We anticipate that the cytosolic ribosome is active at a rate that scales with gene expression.

In addition to identifying a set of genes that appear to be proportionally stably expressed, we have introduced other notions of stable expression, in particular absolute stability. Importantly, normalization steps can affect the types of stability that are discoverable in the data. Technical effects captured by each of notion of stability vary. Thus, these different notions of stability could be used separately to identify specific technical effects, or in conjunction to remove multiple types of technical effects.

While previous sets of stably expressed genes have not been defined by considering a notion of stability, these gene sets can be considered with respect to the notions that we have defined. By nature of being from microarray data, Eisenberg and Levanon [2003] are only able to capture averages across a bulk sample of cells. Since there are no indicators of the number of cells, these genes are likely to be proportionally stable. The characteristics that Eisenberg and Levanon [2003] search for also support that their genes are likely to be the most similar to proportional stability. Lin *et al.* [2017] discover their stably expressed genes from scRNA-seq. Prior to analysis, an RPKM normalization is performed, leading to proportional stability being most prevalent in the data. They then look for genes with high RPKM-normalized expression and low variability. While Lin *et al.* [2017] search for features that are not directly related to proportional stability, the notion of stability is closest to proportional.

## Supporting information

## Acknowledgements

We would like to thank the members of the Michigan Center for Single-Cell Genomic Data Analytics, especially Justin Colacino, Anna Gilbert, Jun Li, and Xiang Zhou, and Lana Garmire and Greg Hunt.

