## Supplementary material for "Stably expressed genes in single-cell RNA-sequencing"

### 1 Supplementary Table Description

We created a table that contains information for human genes, including structural annotations and numerical summaries from our single-cell analysis and GTEx validation. The first ten columns contain indicators for whether the gene was classified as a member of each of our structures; these names are denoted as the full name of the structure followed by "_indicator". The following two columns contain indicators of membership of the gene sets published by Eisenberg and Levanon [2003] and by Lin *et al.* [2017] and denoted with the author names.

Summary statistics, including the mean, standard deviation, and correlation, for each single cell dataset are calculated for the genes expressed in all six scRNA-seq datasets. The mean and standard deviation of the log + 1 transformed expressions for each of the genes is calculated for each of the datasets; these variables are denoted with the five digit GEO accession number (GSE#####) followed by either a "mean" or "sd" suffix to denote which summary. The correlation with the unadjusted cell total is also included for each dataset, as well as the mean unadjusted cell total across the six datasets. Our analysis in the main text primarily looks at correlations with adjusted cell totals; we provide the mean correlation across the six datasets for each gene with each of the adjusted cell totals. These variables have the name "sc_cormean_less" followed by the name of the structure removed during the cell total calculation.

Prior to analysis, the GTEx data is normalized according to an adjusted RPKM transformation. The total number of reads for a given sample is calculated and divided by 1,000,000. Each individual entry is then transformed with log (entry + 1/ total samples per million). Then, the mean and standard deviation of each gene are calculated for each of the tissue types contained within the data. F-statistics and their respective degrees of freedom are calculated for each of the tissue types that contain subtissue types; these F-statistics are a measure of how differentially expressed the gene is between the subtissue types. Note that genes that are stably expressed will have low F-statistics. These variables are denoted with GTEx_tissue type_statistic, where the "tissue type" and "statistic" are replaced with the appropriate designation; the options for the statistic are mean, standard deviation, F statistic, or degrees of freedom. Note that we have removed the "Transformed fibroblasts" from the Skin tissue type, as these were prepared differently than the other samples.

### 2 Data Sources

#### 2.1 Experimental Data

Data were retrieved from the Gene Expression Omnibus [Edgar *et al.*, 2002]. Selection criteria for data included being conducted with the Fluidigm C1 platform for human cells and including ERCC spike-ins. The accession numbers are: GSE77288 [Tung *et al.*, 2017], GSE84686 [Das *et al.*, 2017], GSE79130,

Table 1: Data Source Information

| GEO sample | Tissue type | Number cells | Number genes |
|---|---|---|---|
| 77288 | Stem cells | 2,592 | 20,425 |
| 84686 | T cells | 96 | 25,462 |
| 79130 | Kidney | 47 | 18,730 |
| 89235 | Kidney | 76 | 17,594 |
| 89236 | Nasal epithelium | 96 | 16,160 |
| 89237 | Nasal epithelium | 96 | 15,670 |

GSE89235, GSE89236, and GSE89237 [Arguel *et al.*, 2017]. The last four data sources compose data from one experiment collected by the same lab. All except for GSE84686 also used unique molecular identifiers (UMIs) [Kivioja *et al.*, 2012]. General information on these data sources can be found in Table 1.

Genes in GSE77288 were reported by Ensembl identifiers, while genes in the other five datasets were reported by gene symbol. We transformed the Ensembl identifiers in GSE77288 to gene symbols using getGenes from `mygene` [Wu *et al.*, 2013, Xin *et al.*, 2016]. The Ensembl identifiers for which a gene symbol could not be identified were removed from the dataset, and multiple Ensembl identifiers that matched with the same gene symbol were summed.

One cell was removed from each of GSE89236 and GSE89237. These cells had two and zero UMIs measured, respectively.

#### 2.2 Structural Annotation Dictionary

Our structural annotation dictionary combines a list of human genes from Ensembl's Biomart with the Gene Ontology Consortium's (GO) information on genes gathered with `mygene` in R [Wu *et al.*, 2013, Xin *et al.*, 2016]. Gene names and not Ensembl identifiers were used.

The query through `mygene` returns the top matches from GO for a given gene symbol. We retained only the top match. Any genes not identified from GO were removed. The biological process, molecular function, and cellular component were returned from GO. Each of these fields could have no values or multiple values.

We considered the 1,629 unique cellular components for mapping the genes into ten broad cell structure (or location) categories. The cellular categories that were mapped to the cell structures were identified with a combination of text matching and review. First terms associated with the structure of interest were identified with the text term in the second column of Table 2. The cellular components that matched that text string were reviewed for appropriate fit. Cellular component categories that did not seem to fit were removed with the text string(s) specified in the third column of Table 2. The cell structures we considered are: nucleus, endoplasmic reticulum, Golgi apparatus, cytoplasm, membrane, mitochondria, ribosome, mitochondrial ribosome, ribonucleoprotein

Table 2: Structural Annotation Dictionary Criteria

| Structure | Matching Term | Removal Term(s) |
|---|---|---|
| Nucleus | nucl | mitochond, nucleotide, ribonucleoprotein, male, endonuclease, ribonuclease |
| Endoplasmic reticulum | reticu | sarcoplasmic, cortical |
| Golgi bodies | Golgi | vesicle |
| Cytoplasm | cytop | side of, axon, nuclear pore |
| Membrane | membrane, cytoplasmic vesicle membrane, azurophil granule membrane | mitochond, Golgi, nucl, endoplasmic, organ, secretory, lysosom, endosome, ruffle, photoreceptor, vacuol, platelet, synaptic, peroxisomal, sperm, vesic, acrosomal, granule, basement, melanosome, coat, neuron, phago, sarcoplasmic, muscle, omegasome, tubular, spine, attack, junctional, transport, ER, chain, bounded, succinate, lamellar |
| Ribosome | ribosom | mitochond |
| Mitochondria | mitochond | ribosom |
| Mitochondria ribosome | mitochond, ribosom | |
| Ribonucleoprotein complex | nucl, ribonucleoprotein | |
| Cytosolic ribosome | ribosom, cytosolic | |

complex, and cytosolic ribosome.

The mapping was not one-to-one. In fact, each cellular component could be mapped to no structure, one structure, or multiple structures. There was nesting within the ribosomal structures with the cytosolic ribosome being a subset of the ribosome. In addition, each gene could have zero to 41 cellular components as annotated by GO, with 75% of genes having four or fewer cellular components.

#### 2.3 Cytosolic Ribosomal Genes

The 101 cytosolic ribosome genes from the scRNA-seq datasets are:

COA1, DDX3X, EIF2A, EIF2AK4, EIF2D, EIF4G1, FXR2, GEMIN5, LARP 4, MRPL1, MRPL4, MRPS18A, MRPS18C, MARPS5, NAA10, NHP2, NUFIP1, RPL10, RPL10A, RPL11, RPL12, RPL13 RPL13A, RPL14, RPL23A, RPL24, RPL26, RPL26L1, RPL27, RPL27A, RPL28, RPL29, RPL3, RPL30,

Table 3: Overlap of Gene Sets

| Structure | All Genes | E & L | Lin et al. | Top 100 |
|---|---|---|---|---|
| Number of Genes | 8,601 | 470 | 967 | 100 |
| Eisenberg & Levanon (2003) | 470 | 470 | 94 | 51 |
| Lin et al. (2017) | 967 | 94 | 967 | 37 |
| Top 100 Correlations | 100 | 51 | 37 | 100 |

RPL31, RPL32, RPL34, RPL35, RPL35A, RPL36, RPL36A, RPL36AL, RPL37, RPL37A, RPL38, RPL39, RPL39L, RPL4, RPL41, RPL5, RPL6, RPL7, RPL7A, RPL7L1, RPL8, RPL9, RPLP0, RPLP1, RPLP2, RPS10, RPS11, RPS12, RPS14, RPS15, RPS15A, RPS16, RPS18, RPS19, RPS2, RPS20, RPS21, RPS23, RPS24, RPS25, RPS26, RPS27, RPS27A, RPS27L, RPS28, RPS29, RPS3, RPS3A, RPS4X, RPS5, RPS6, RPS7, RPS9, RPSA, RSL1D1, RSL24D1, SURF6, ZNF622

The 89 cytosolic ribosome genes from the GTEx dataset are:

APOD, DDX3X, EIF2A, EIF2AK4, FXR2, GEMIN5, HBA1, HBA2, MCTS1, MRPL4, MRPS18A, MRPS18C, MRPS5, NAA11, NHP2, NUFIP1, PPARGC1A, RPL10A, RPL10L, RPL11, RPL12, RPL13, RPL13AP3, RPL14, RPL17, RPL18, RPL18A, RPL19, RPL21, RPL22, RPL23, RPL23A, RPL24, RPL26, RPL26L1, RPL27, RPL27A, RPL28, RPL29, RPL30, RPL31, RPL32, RPL36, RPL36A, RPL36AL, RPL37, RPL37A, RPL38, RPL39, RPL39L, RPL39P5, RPL3L, RPL41, RPL7, RPLP1, RPLP2, RPS10, RPS10P5, RPS11, RPS12, RPS13, RPS14, RPS15, RPS16, RPS17, RPS18, RPS19, RPS20, RPS23, RPS25, RPS27, RPS27A, RPS27L, RPS28, RPS29, RPS3A, RPS4X, RPS5, RPS6, RPS7, RPS8, RPS9, RPSA, RSL1D1, RSL24D1, SNU13, SURF6, UBA52, ZNF622

### 3 Stability Measures

Let $i$ represent the gene, $j$ a dataset, $k$ a cell, $\mathcal{G}$ some set of genes, and $c_{i,j,k}$ the expression recorded for gene $i$ in cell $k$ in dataset $j$. Let $s_{\mathcal{G},j}$ be a vector representing the sum of the set of genes $\mathcal{G}$; in other words, $s_{\mathcal{G},j,k} = \sum_{i \in \mathcal{G}} c_{i,j,k}$ is the $k$th entry in the vector $s_{\mathcal{G},j}$. Conversely, let $s_{-\mathcal{G},j}$ be a vector representing the sum of the genes not contained in the set of genes $\mathcal{G}$, or $s_{-\mathcal{G},j,k} = \sum_{i \notin \mathcal{G}} c_{i,j,k}$ is the $k$th entry of the vector $s_{-\mathcal{G},j}$.

The correlations that serve as a measure of absolute stability for a given gene $i$ and a given dataset $j$ are calculated by $r_{ERCC,i,j} = cor(c_{i,j}, s_{\mathcal{E},j})$ where $\mathcal{E}$ is the set of ERCC spike-ins. The overall measure of absolute stability for gene $i$ is $r_{ERCC,i} = \frac{1}{6}\sum_{j=1}^{6} r_{ERCC,i,j}$.

The correlations that serve as a measure of proportional stability for a given gene $i$, a given dataset $j$, and a given cell structure with gene set $\mathcal{S}$ are defined as $r_{\mathcal{S},i,j} = cor(c_{i,j}, s_{-\mathcal{S},j})$. The overall measure of stability for gene $i$ is $r_{\mathcal{S},i} = \frac{1}{6}\sum_{j=1}^{6} r_{\mathcal{S},i,j}$.

We calculated an additional measure of proportional stability that was comparable across all genes. This uses an unadjusted cell total, unlike the measures of proportional stability defined above that uses a cell total adjusting for a given structure. $s_j$ is a vector where $s_{j,k} = \sum_{i=1}^{I} c_{i,j,k}$; that is, we sum over all of the genes for a given cell. The measure for a given dataset is $r_{i,j} = cor(c_{i,j}, s_j)$ and is summarized over the 6 datasets by $r_i = \frac{1}{6}\sum_{j=1}^{6} r_{i,j}$.

### 4 Additional Figures

The following figures provide extensions or additional information to those given in the paper. Figure 2 to Figure 8 in particular provide additional information that relate to many figures directly contained within the paper.

#### 4.1 Correlation Histograms for Each Dataset

Supplementary Figures 9 to 20 show the histograms of ERCC correlations found in Supplementary Figures 2 and 3 for each of the distinct datasets.

Supplementary Figures 21 to 32 show the histograms of correlations found in Figure 3 and Supplementary Figure 4 for each of the distinct datasets.

#### 4.2 Singular Value Decomposition of Single GTEx Tissue Types

Supplementary Figures 33 to 38 show the first two left singular values from a singular value decomposition from a single tissue. These are colored by detailed tissue subtypes. In each of these figures, the top left panel shows the singular value decomposition using all genes as features. The top right panel shows the singular value decomposition using the bulk stably expressed genes from Eisenberg and Levanon [2003] as features. The bottom left panel shows the singular value decomposition from the single cell stably expressed genes from Lin *et al.* [2017] as features. The bottom right panel shows the singular value decomposition using the 89 cytosolic ribosomal genes found in the GTEx data as features. Note that we removed the Transformed fibroblasts subtype from the skin samples, due to technical differences in their preparation prior to experimental procedures.

#### 4.3 Singular Value Decomposition of Two GTEx Tissue Types

Supplementary Figures 39 to 80 show the singular value decomposition using the first two left singular values. These figures are colored by the two tissue types. The top left panel shows the singular value decomposition using all genes as features. The top right panel shows the singular value decomposition using the bulk genes presented in Eisenberg and Levanon [2003] as features. The bottom left uses the single cell stably expressed genes from Lin *et al.* [2017]

as features. The bottom right uses the cytosolic ribosomal genes as features. The first figure for each of these tissue types are colored by the general tissue type. The second figure for each of these tissue types are colored by the detailed sub-tissue type with shapes representing the two different tissue types. Genes that are more stably expressed across tissue types result in clusters that are less easily distinguishable. In these figures, the singular value decomposition from the cytosolic ribosomal genes result in the most overlap between clusters, indicating that the cytosolic ribosomal genes are more stably expressed across both general tissue types and across the more detailed tissue subtypes.

#### 4.4 Singular Value Decomposition With Equal Number of Genes from GTEx

The four different singular value decompositions shown in Supplementary Figures 33 to 80 have different numbers of genes used as features; we also provide the analogous figures that have been randomly subsetted to 89 genes. Supplementary Figures 81 to 106 show that the number of features does not explain the differences in separation of tissue clusters.

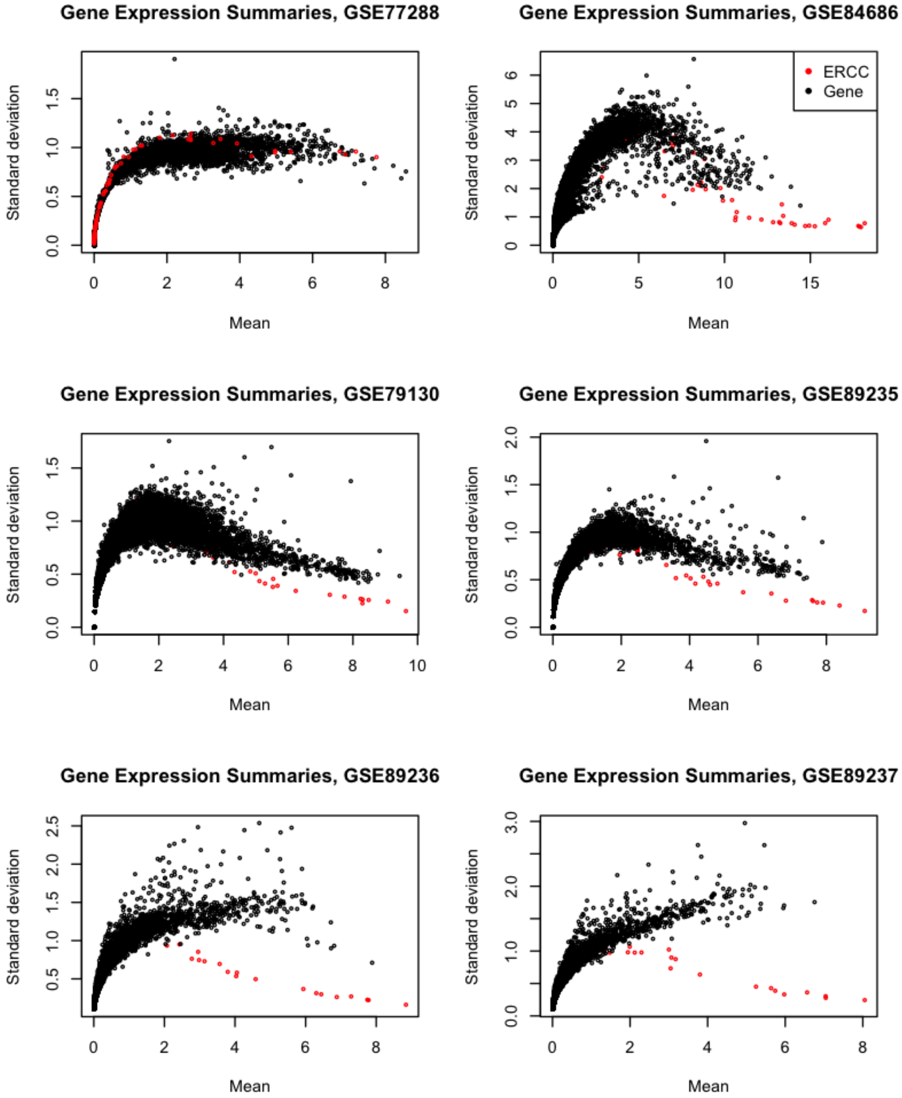


Figure 1: Scatterplots of the mean and standard deviation of ERCC spike-ins and the endogenous genes on the log scale for each of the six datasets. From these graphs, it appears that the variance exhibited by the spike-ins is smaller than those for endogenous genes, even after accounting for the mean measured expression.

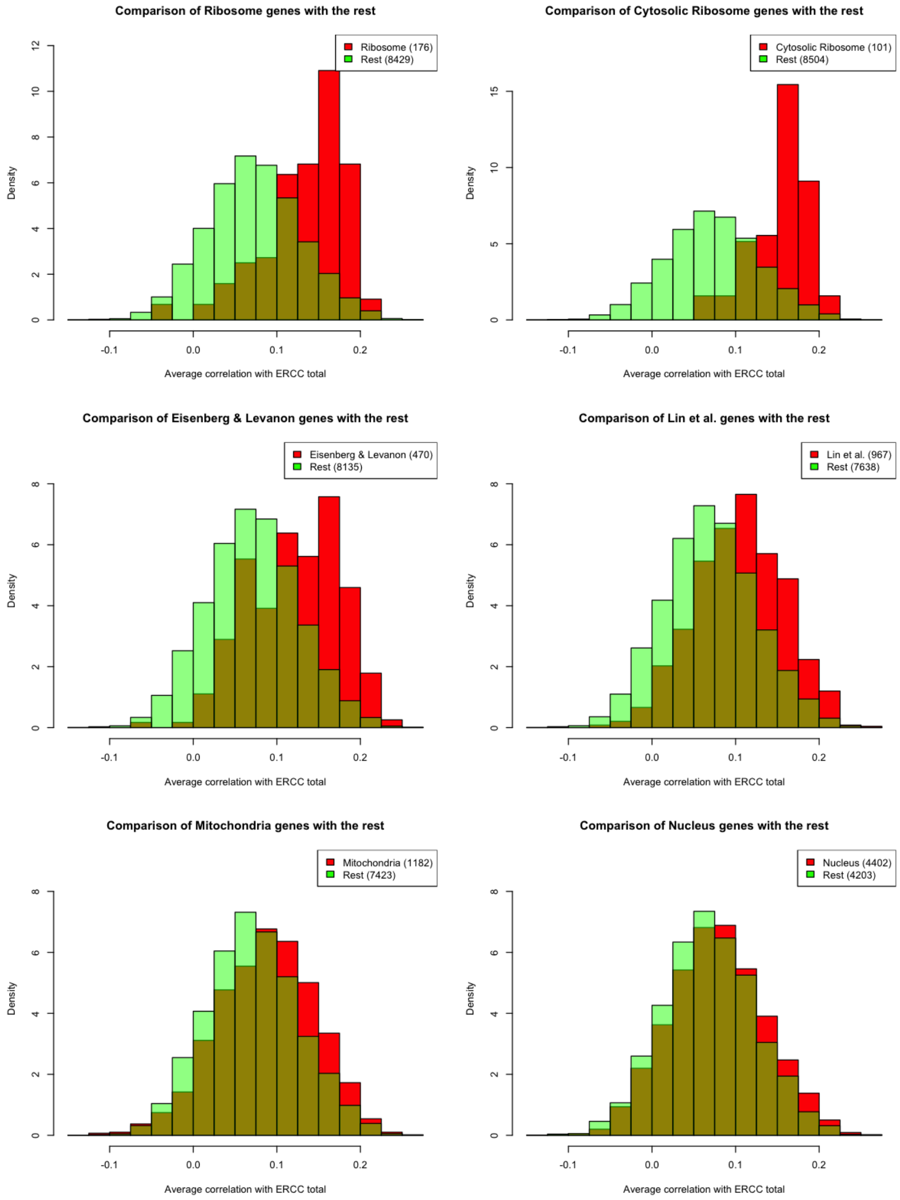


Figure 2: Histograms of the average correlations of each gene with the ERCC total over the six datasets considered, with comparisons between the gene set of interest and the remaining genes. Note that the x-axis ranges from -0.15 to 0.275.

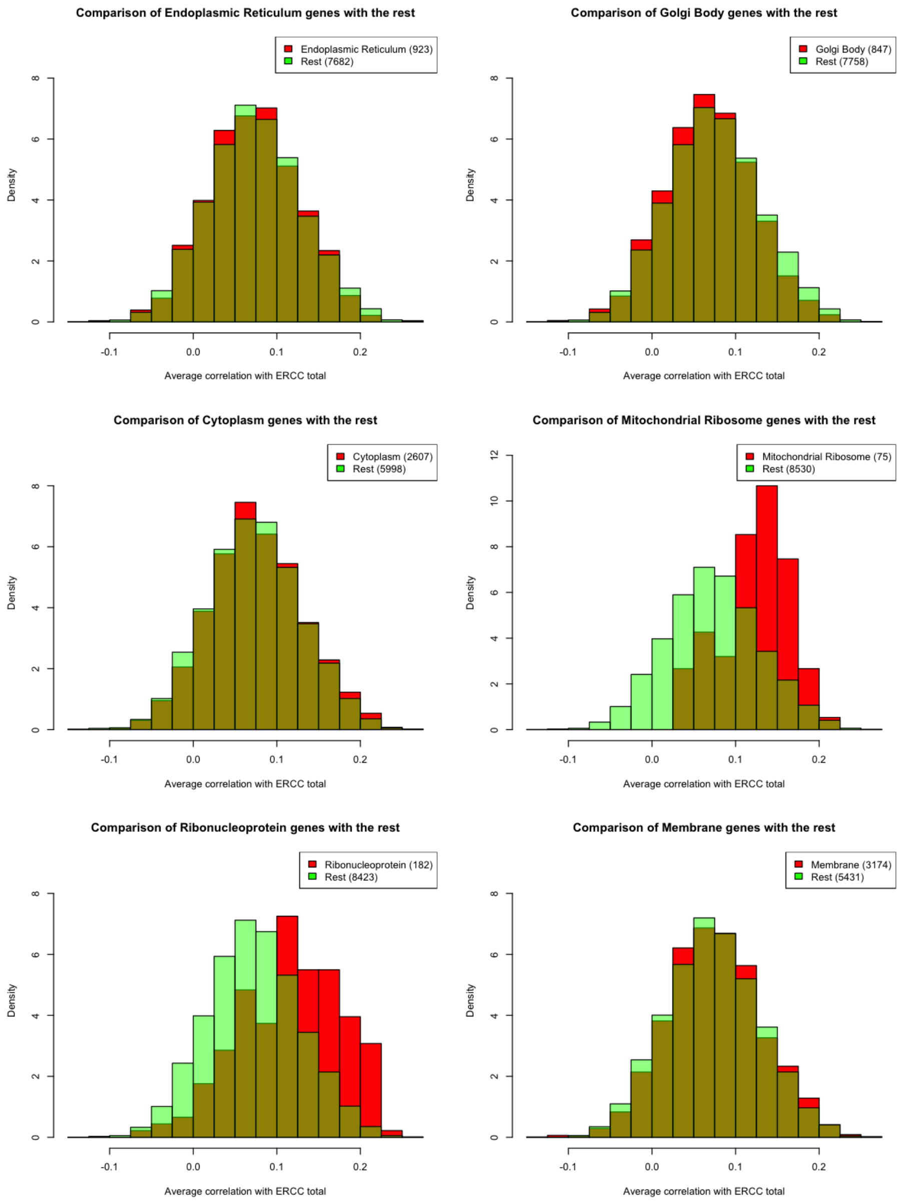


Figure 3: Histograms of the average correlations of each gene with the ERCC total over the six datasets considered, with comparisons between the gene set of interest and the remaining genes. Note that the x-axis ranges from -0.15 to 0.275.

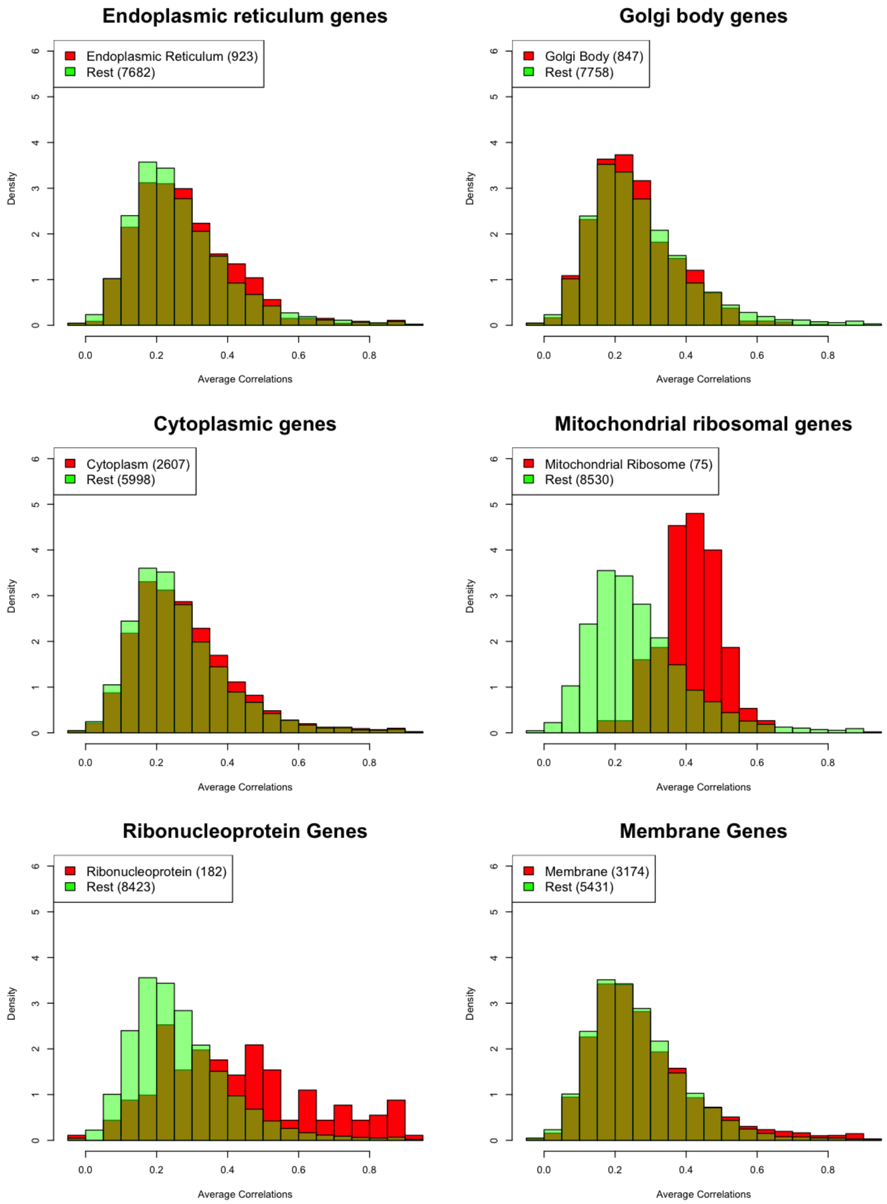


Figure 4: Histograms of the average correlations of each gene over the six datasets considered, with comparisons between the set of genes of interest and the remaining genes.

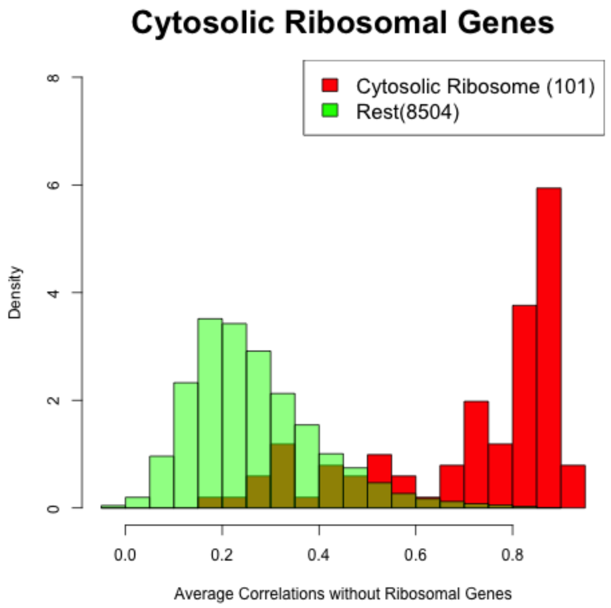


Figure 5: Histogram of the average correlation of cytosolic ribosomal genes compared to the remaining genes when correlated with the adjusted cell total with respect to the ribosomal genes. Because the cytosolic ribosomal genes are a subset of the ribosomal genes, we consider the possibility that the correlations seen in Figure 3 could be inflated due to the correlation between cytosolic ribosomal genes and the remaining ribosomal genes. Using the adjusted cell total removing the ribosomal genes should reduce the artificial inflation. The figure here displays a similar distribution to the one in Figure 3, indicating that any inflation was minimal.

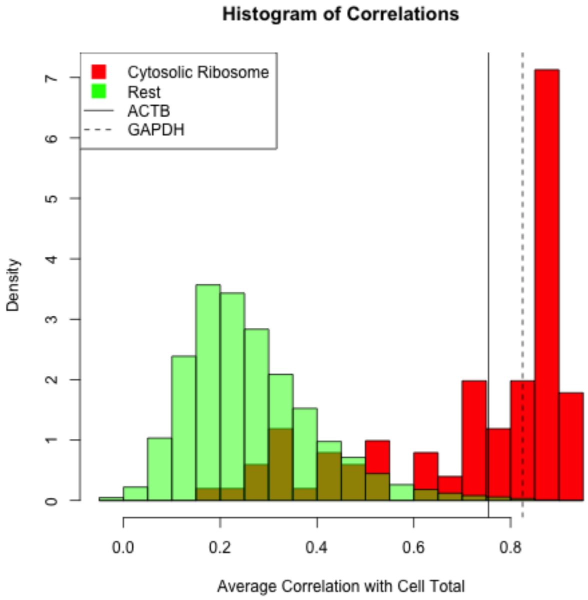


Figure 6: The average correlations of Beta-actin (ACTB) and GAPDH with the cell total over the six datasets. Note that, unlike in Figure 3, the correlations here are with the unadjusted cell total rather than an adjusted cell total. This allows correlations between ACTB, GAPDH, and the cytosolic ribosomal genes to be comparable.

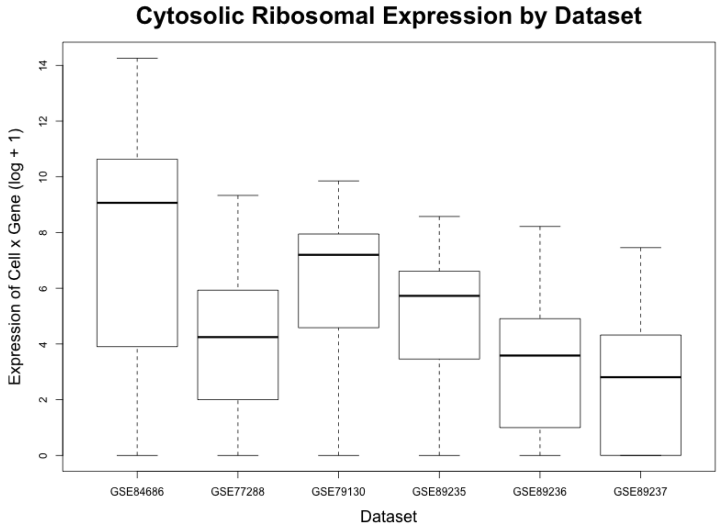


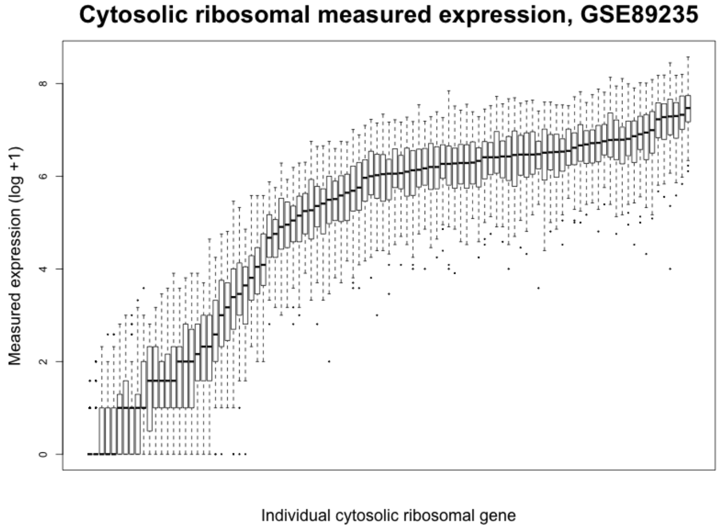


Figure 7: Summary characteristics of the expression of cytosolic ribosomal genes across cells and across datasets. The boxplot on the top shows all of the expression measurements by dataset for the cytosolic ribosomal genes, transformed with the log + 1. For comparison, the proportion of zeros in each of these datasets is 80%, 41%, 42%, 63%, 76%, and 82%, respectively and means (on the log + 1 scale) of 1.23, 1.26, 1.37, 0.71, 0.45, and 0.30, respectively. The boxplot on the bottom looks at the expression (log + 1) of each of the cytosolic ribosomal genes within GSE89235. Analogous boxplots for each of the datasets can be found in Supplementary Figure 8.

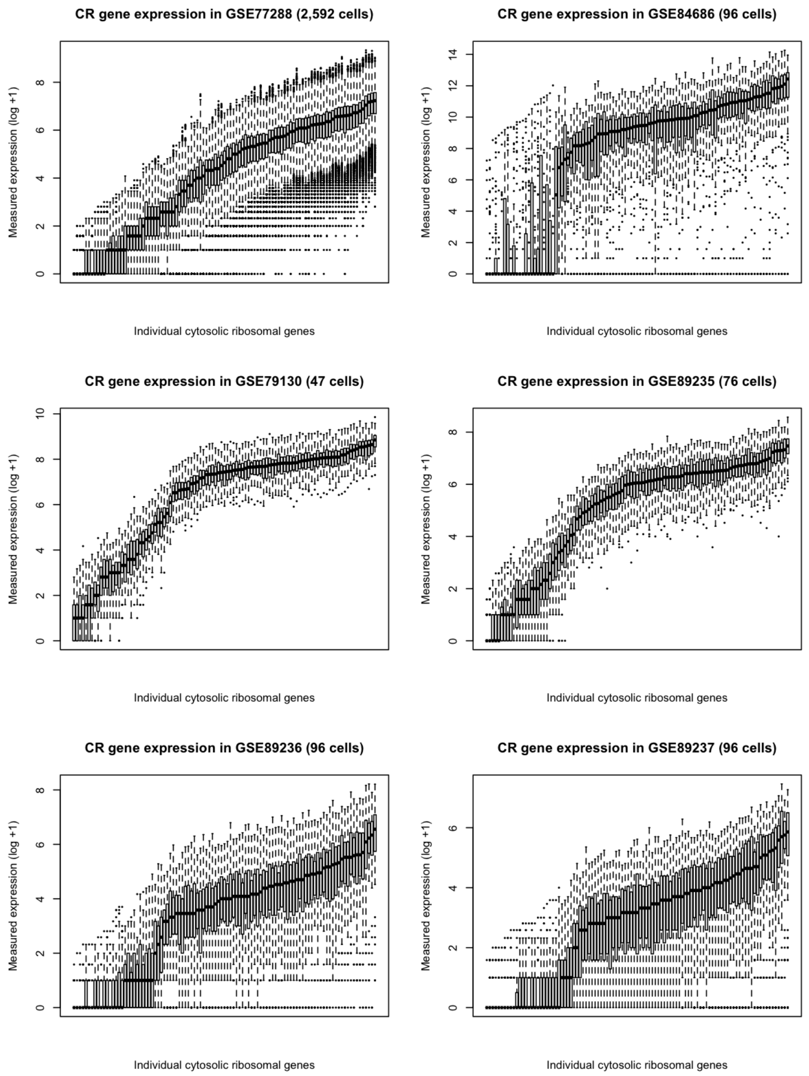


Figure 8: The boxplots of the distributions of each cytosolic ribosomal gene in each dataset. This expands the bottom panel of Supplementary Figure 7 to all datasets. We can observe the variation and relative stability in the expression of cytosolic ribosomal genes.

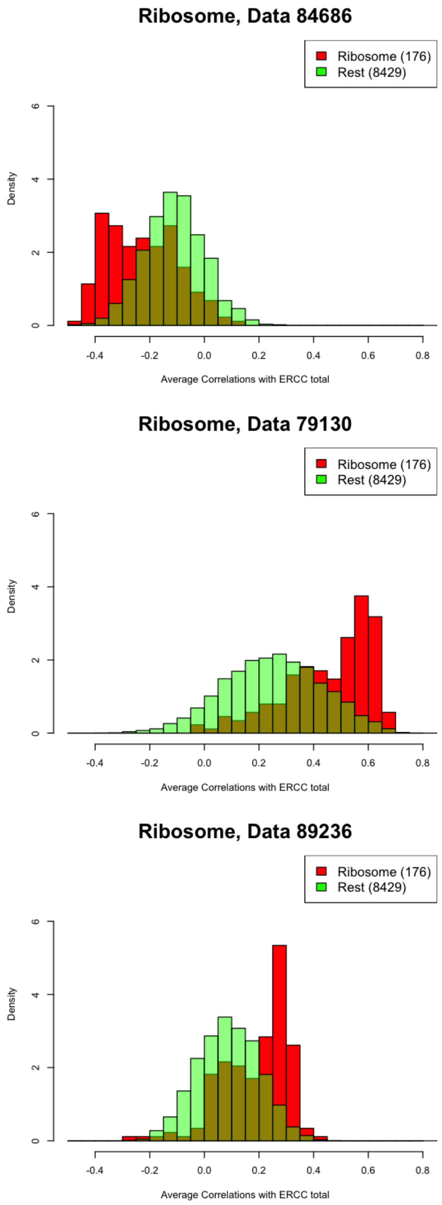


Figure 9: Histograms of the correlations of each gene with the ERCC total for each dataset, with comparisons between the ribosomal genes and the remaining genes.

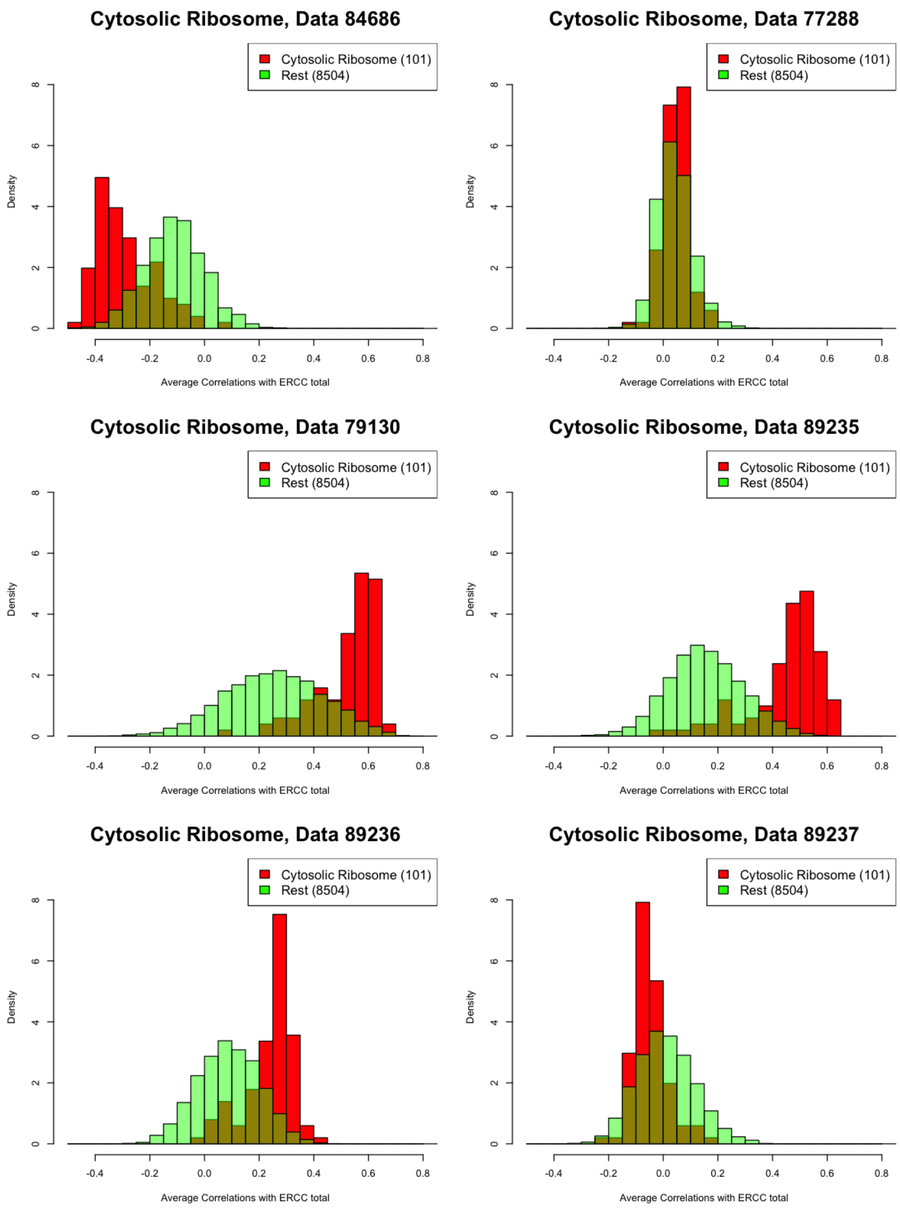


Figure 10: Histogram of the correlations of each gene with the ERCC total for each dataset, with comparisons between the cytosolic ribosomal genes and the remaining genes.

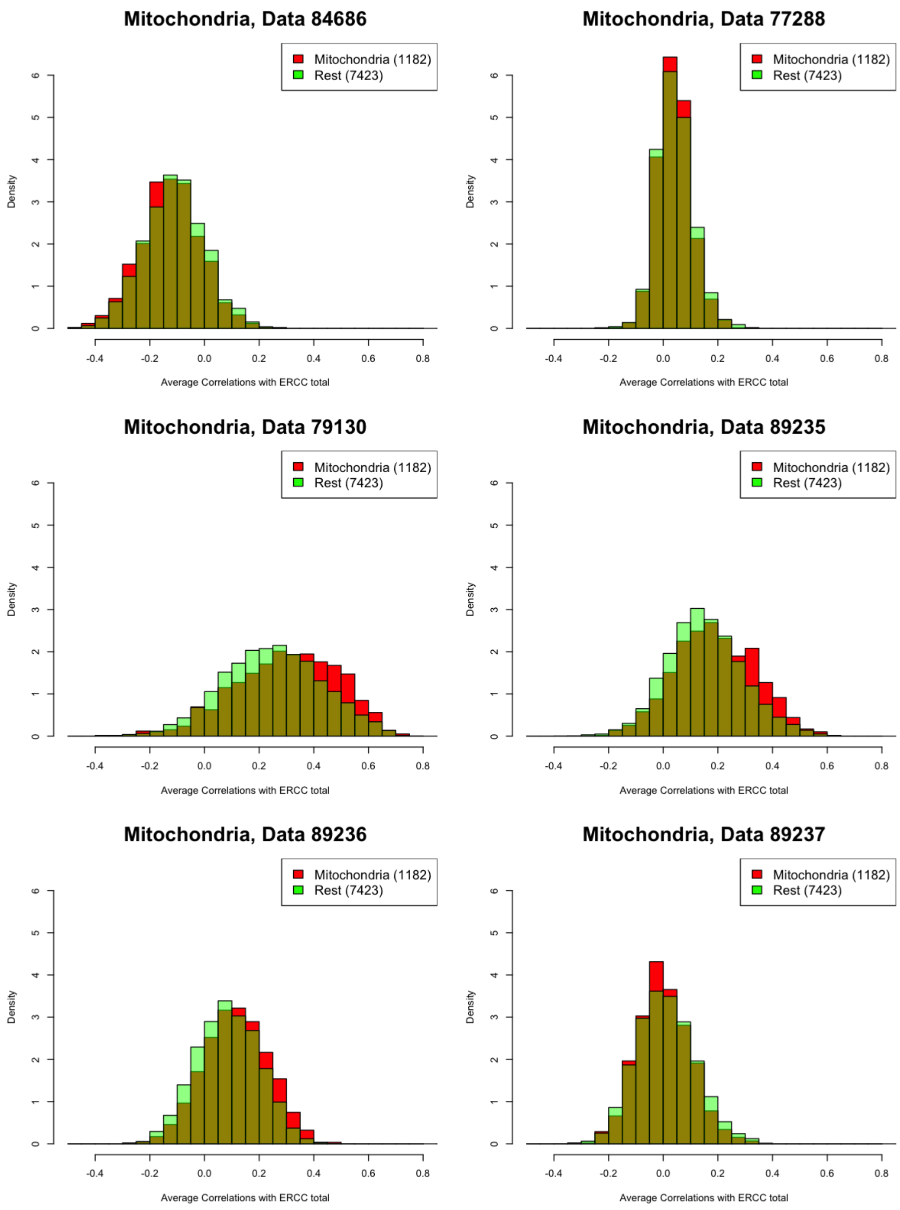


Figure 11: Histogram of the correlations of each gene with the ERCC total for each dataset, with comparisons between the mitochondrial genes and the remaining genes.

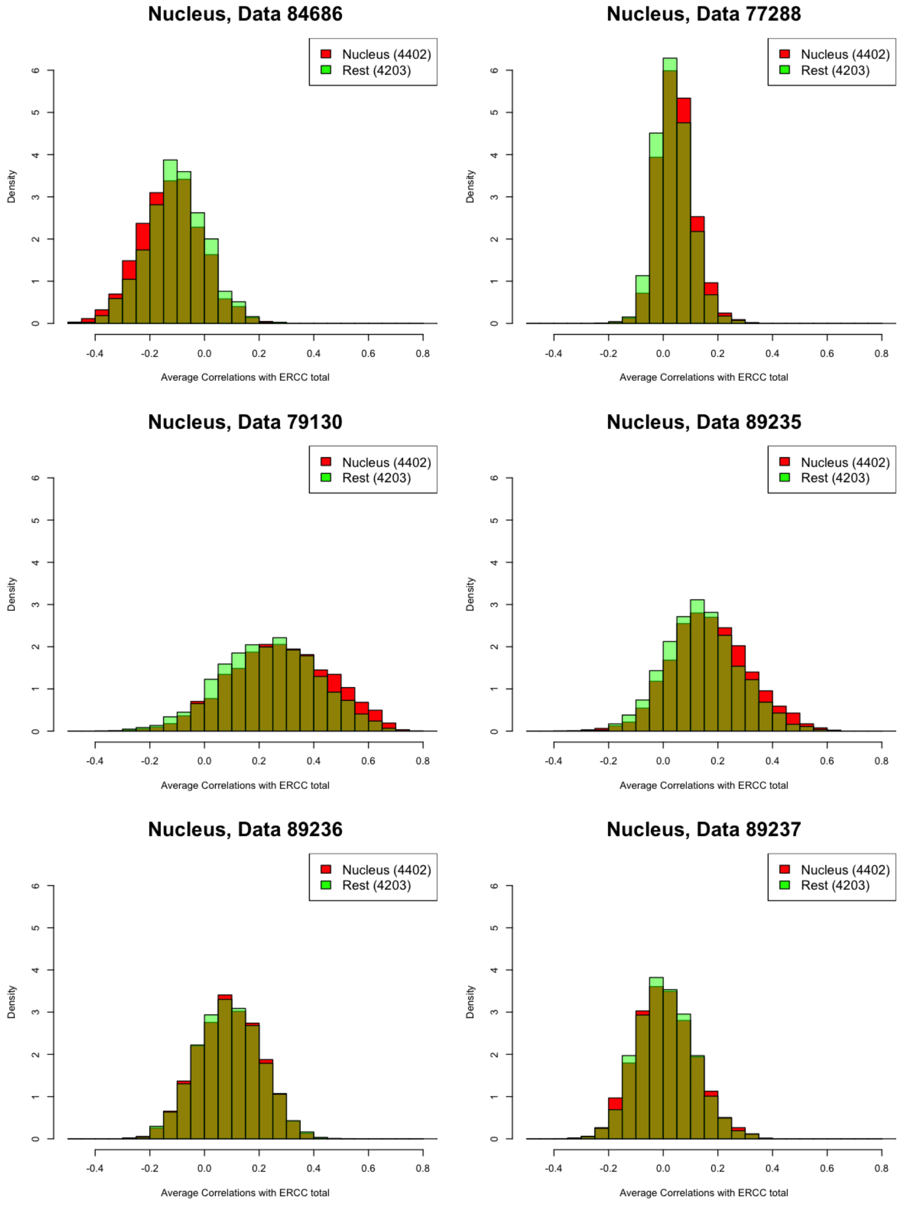


Figure 12: Histogram of the correlations of each gene with the ERCC total for each dataset, with comparisons between the nuclear genes and the remaining genes.

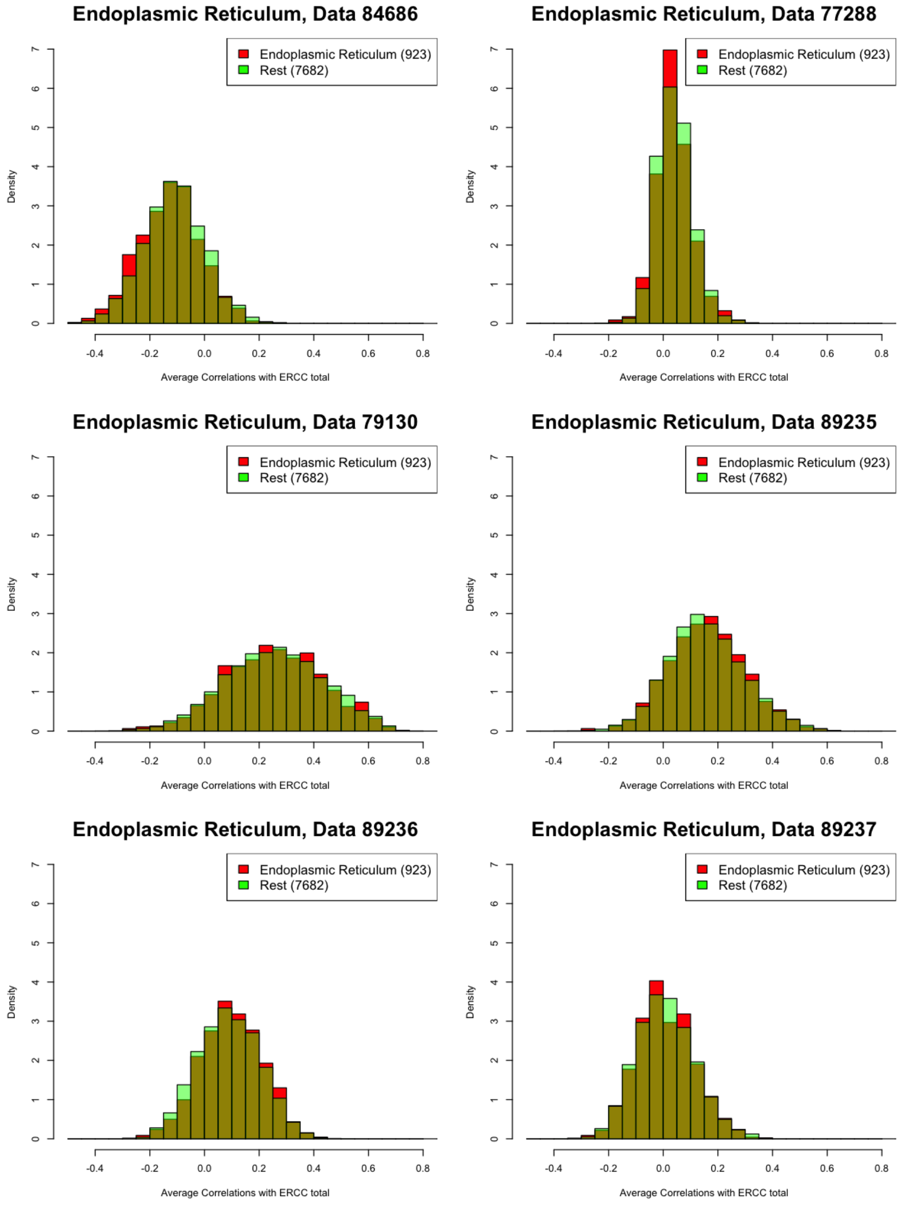


Figure 13: Histogram of the correlations of each gene with the ERCC total for each dataset, with comparisons between the endoplasmic reticulum genes and the remaining genes.

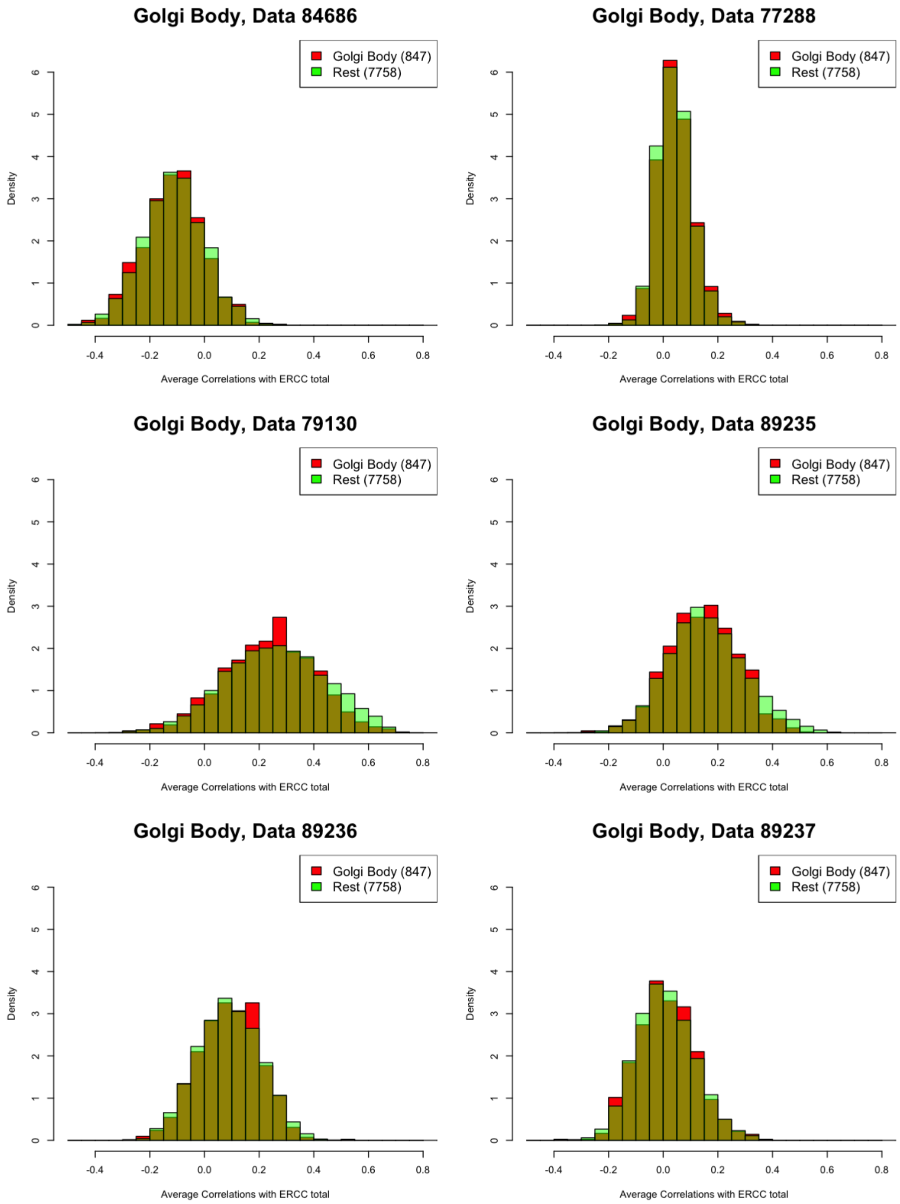


Figure 14: Histogram of the correlations of each gene with the ERCC total for each dataset, with comparisons between the Golgi body genes and the remaining genes.

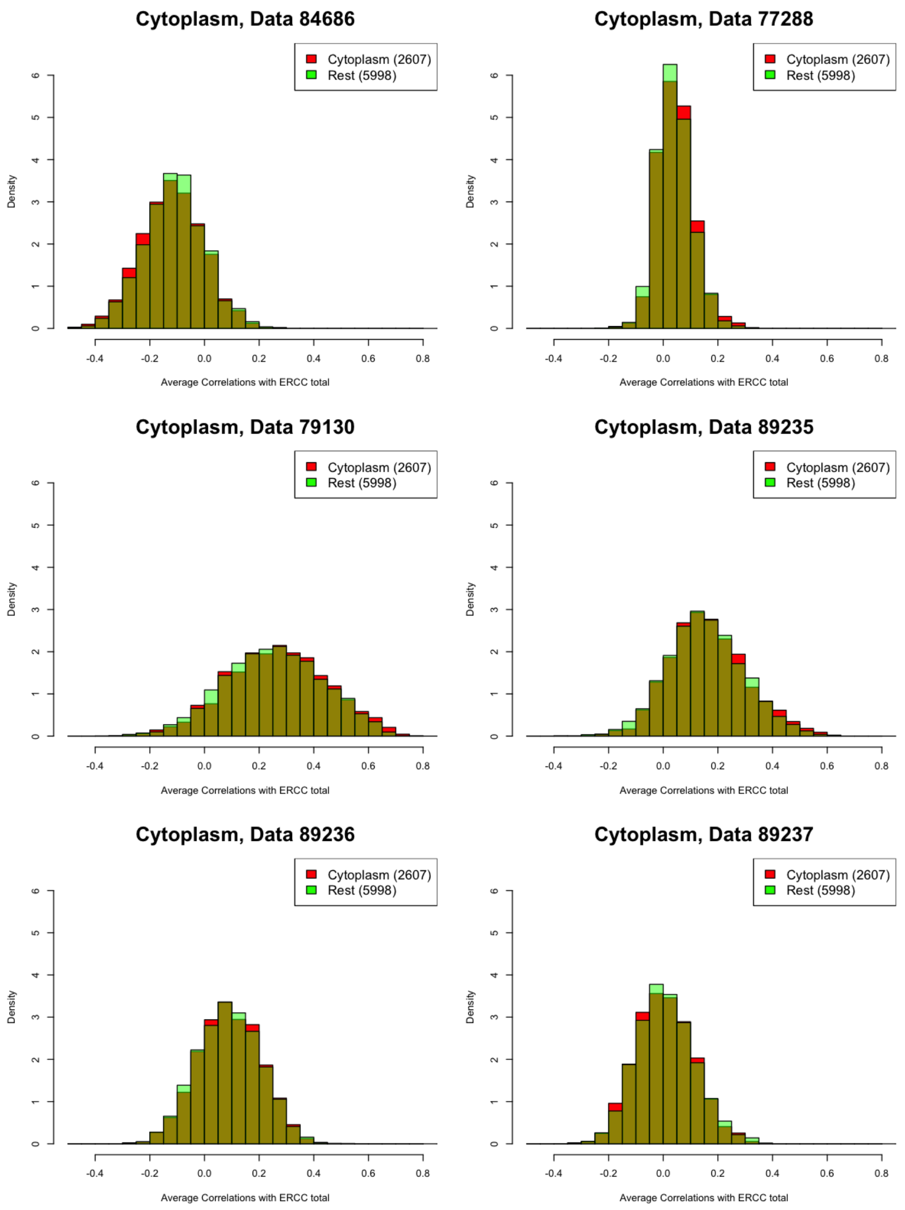


Figure 15: Histogram of the correlations of each gene with the ERCC total for each dataset, with comparisons between the cytoplasmic genes and the remaining genes.

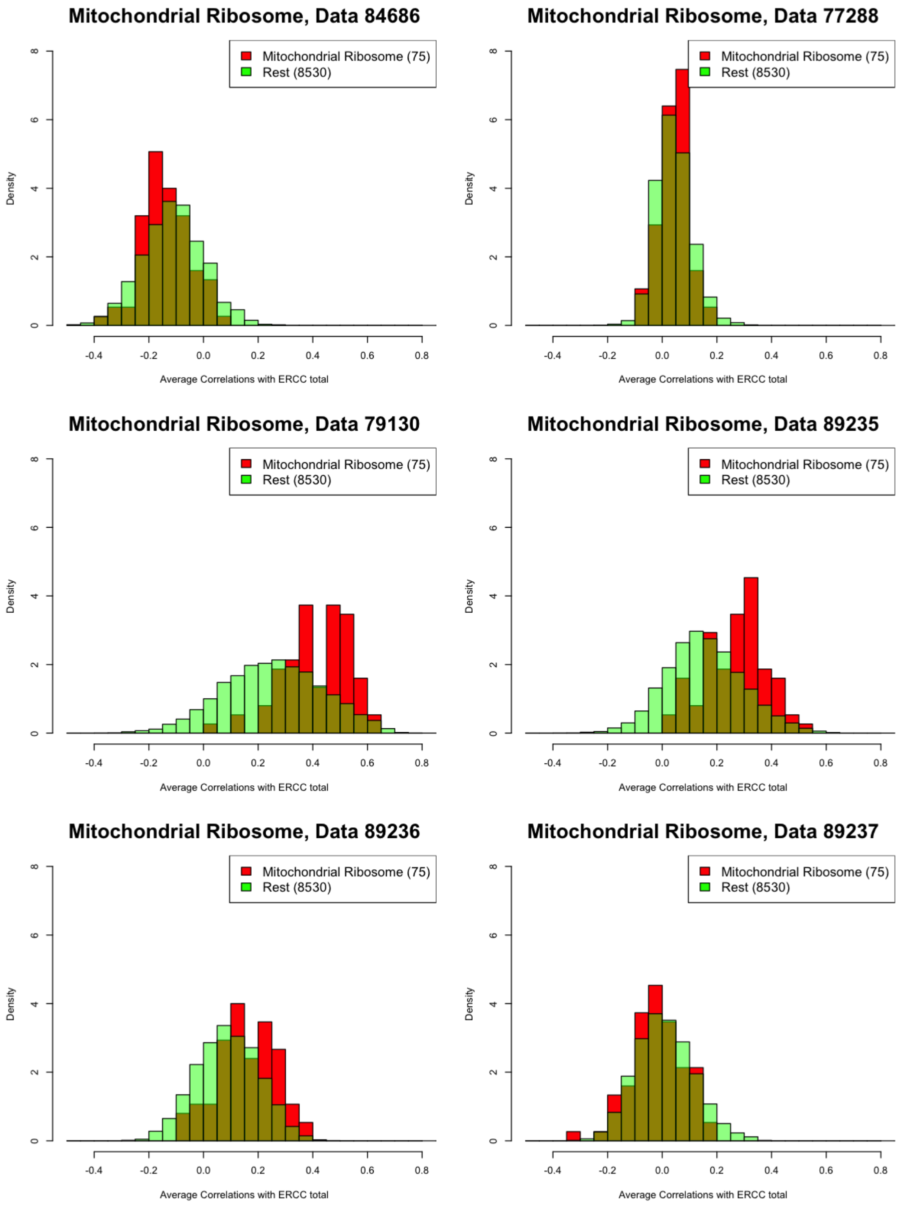


Figure 16: Histogram of the correlations of each gene with the ERCC total for each dataset, with comparisons between the mitochondrial ribosomal genes and the remaining genes.

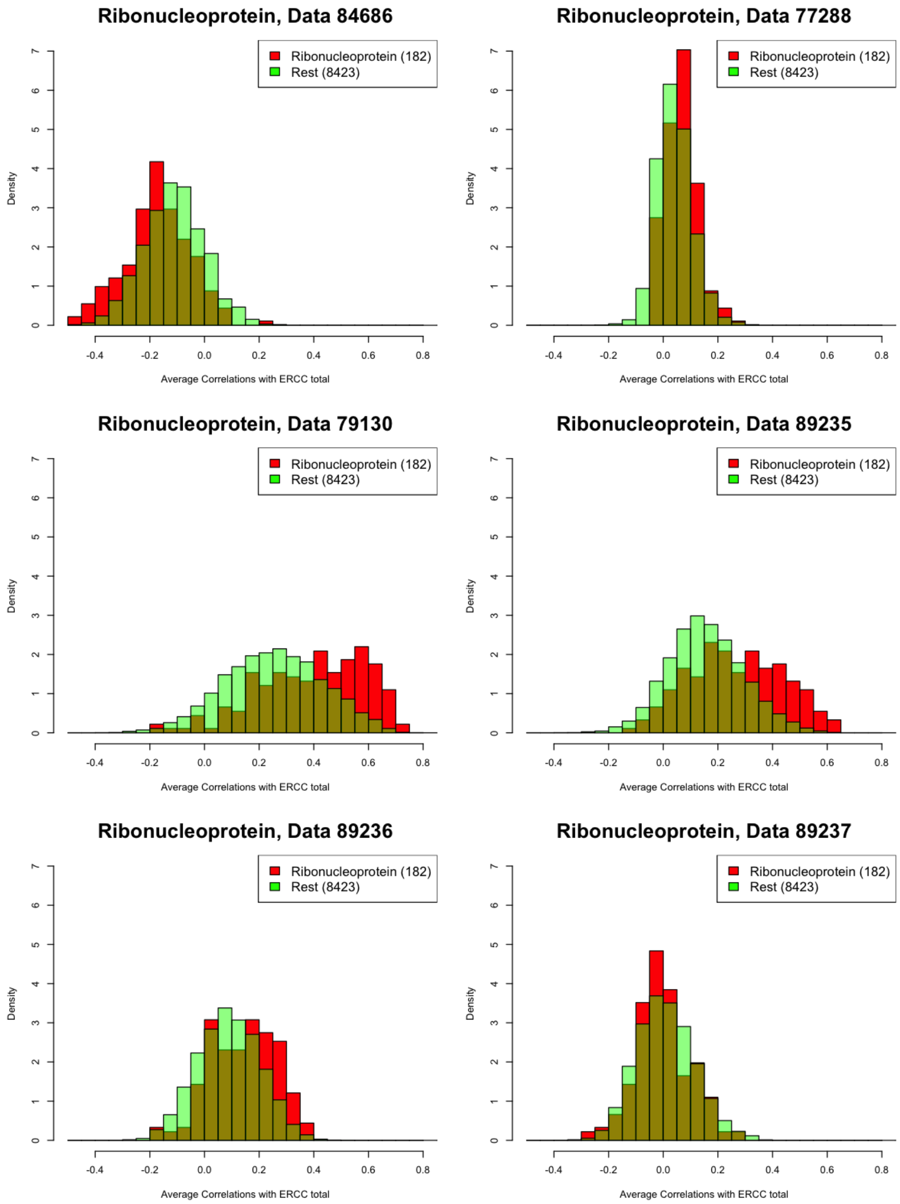


Figure 17: Histogram of the correlations of each gene with the ERCC total for each dataset, with comparisons between the ribonucleoprotein genes and the remaining genes.

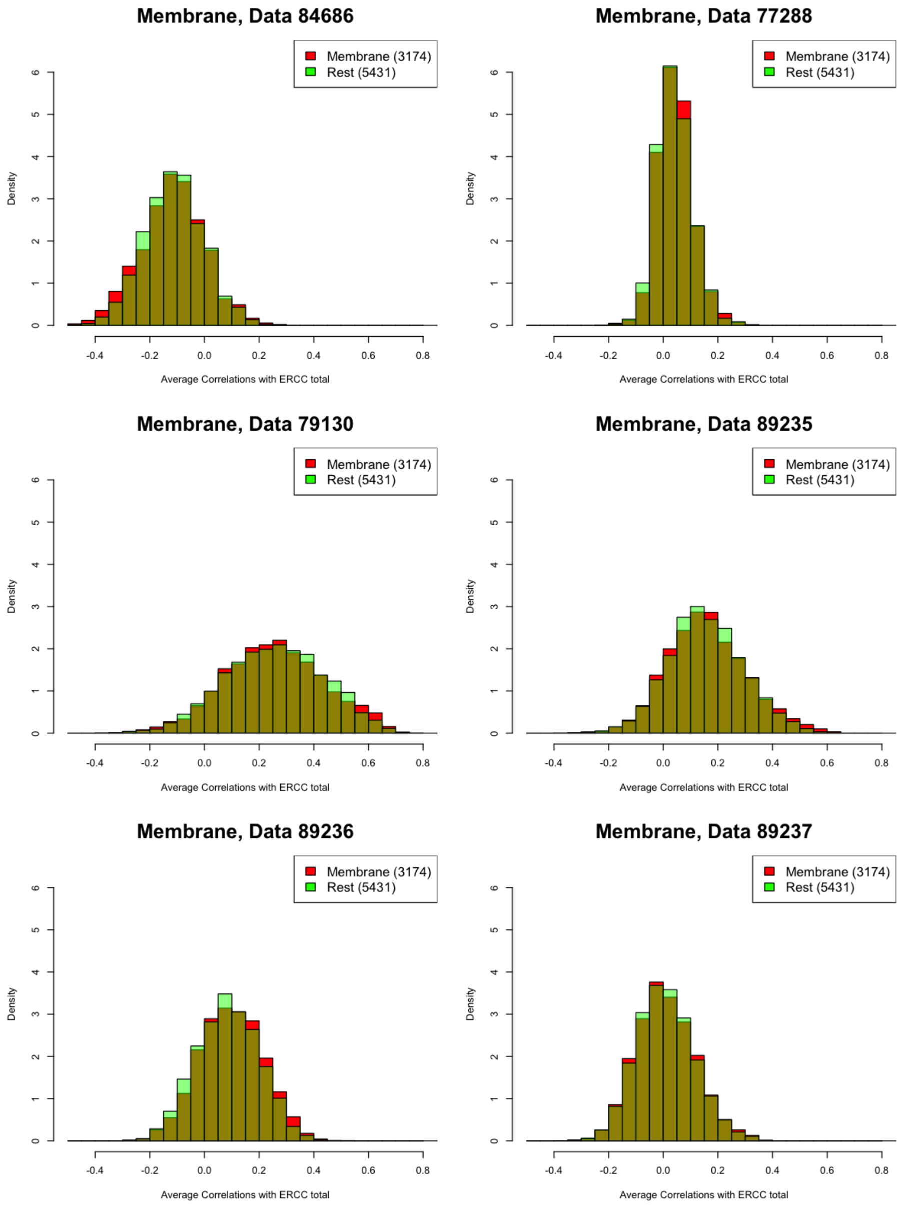


Figure 18: Histogram of the correlations of each gene with the ERCC total for each dataset, with comparisons between the membrane genes and the remaining genes.

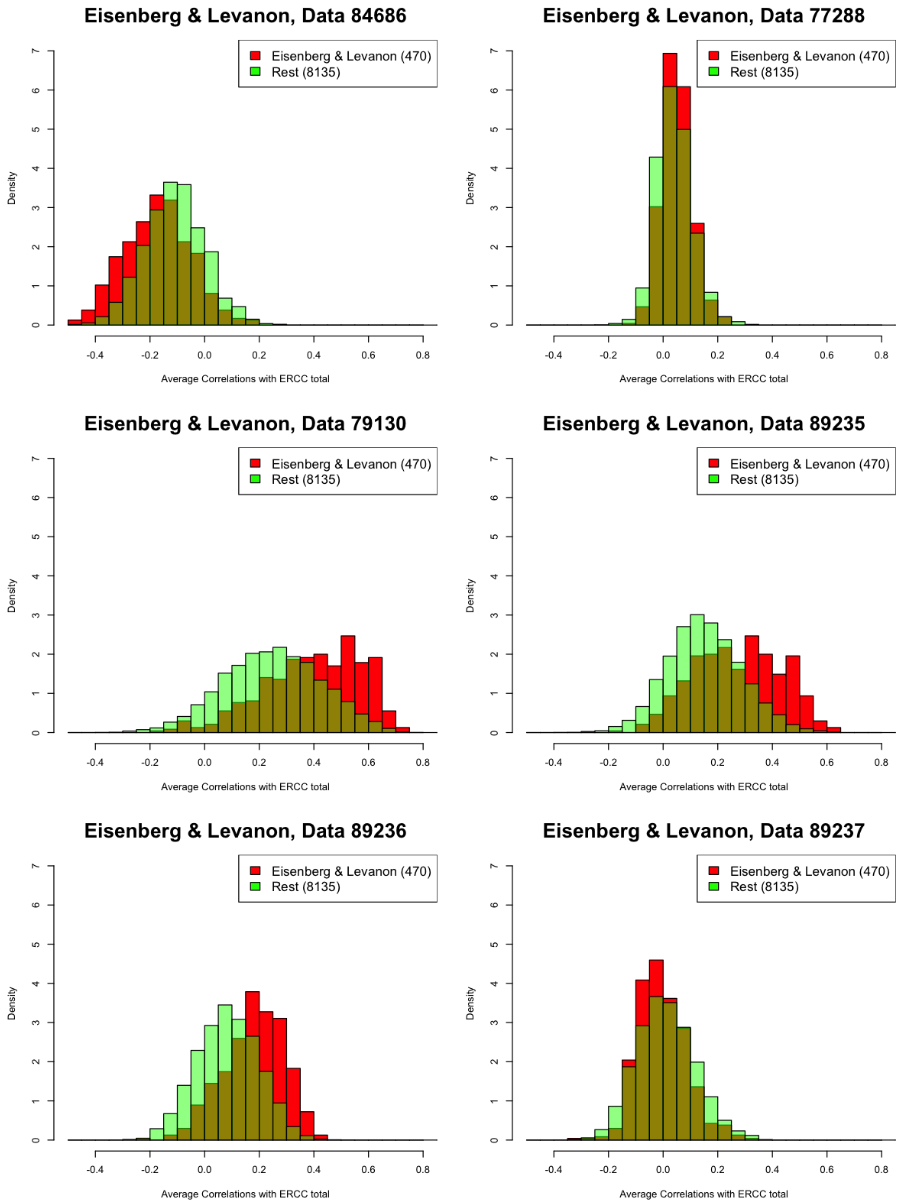


Figure 19: Histogram of the correlations of each gene with the ERCC total for each dataset, with comparisons between the stably expressed genes from Eisenberg and Levanon [2003] and the remaining genes.

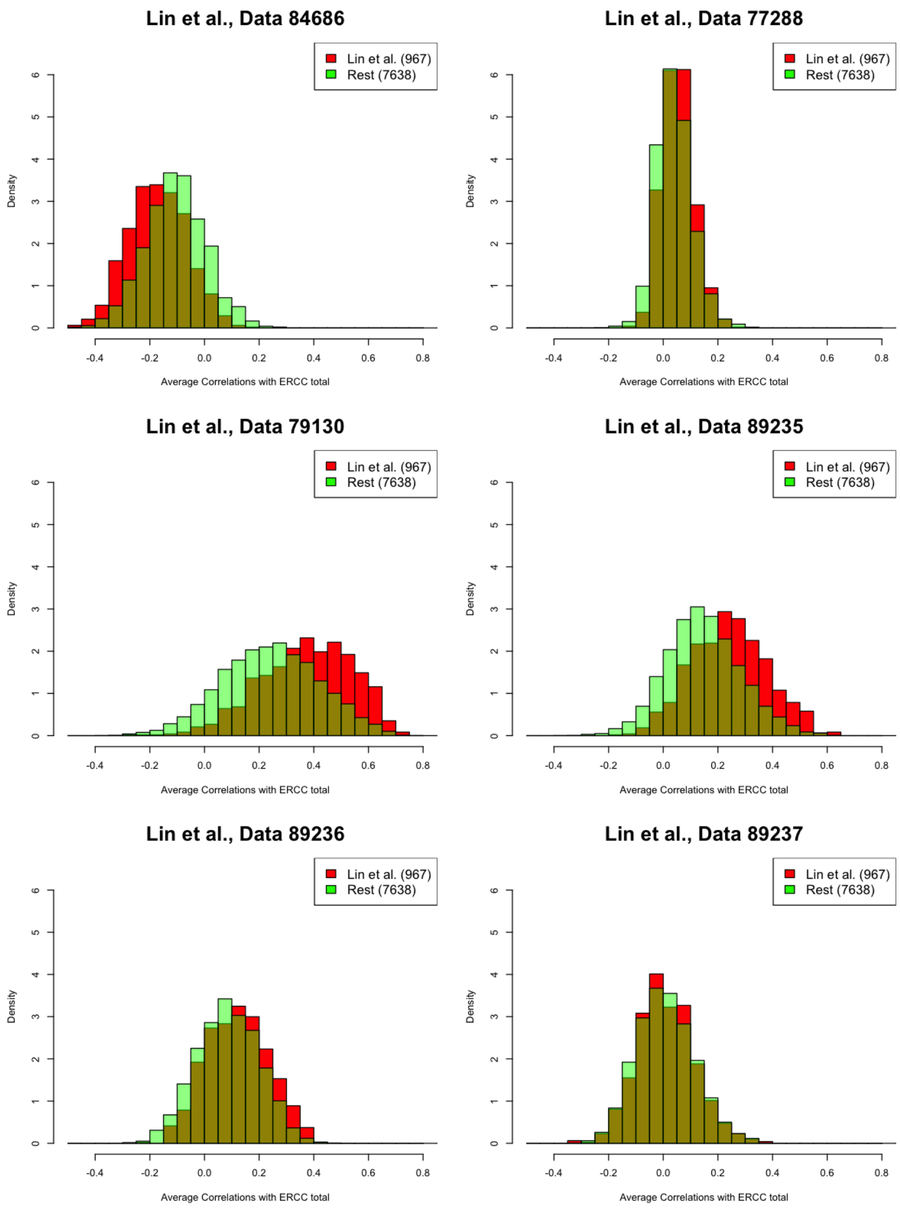


Figure 20: Histogram of the correlations of each gene with the ERCC total for each dataset, with comparisons between the single cell stably expressed genes from Lin *et al.* [2017] and the remaining genes.

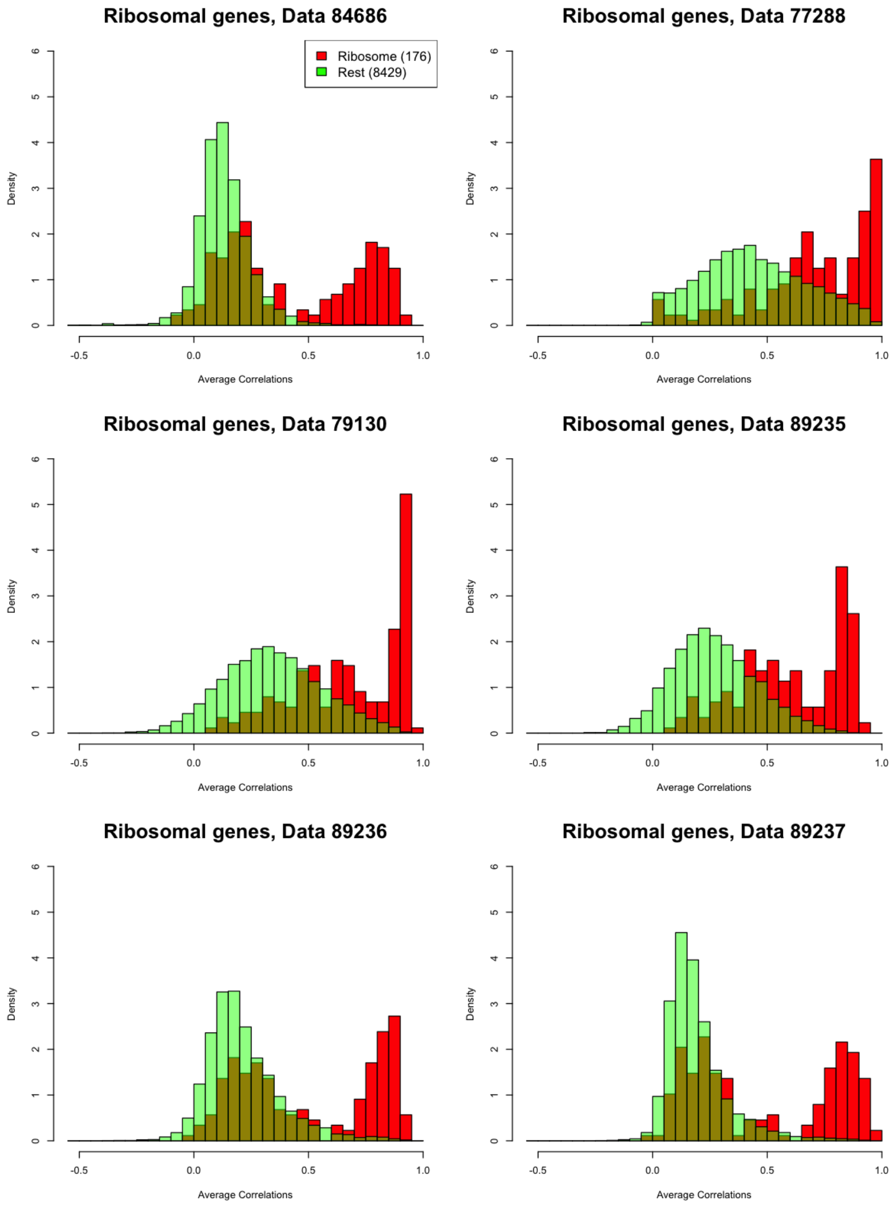


Figure 21: Histograms of the correlations of each gene with the adjusted cell total for ribosomal genes for each dataset, with comparisons between the ribosomal genes and the remaining genes.

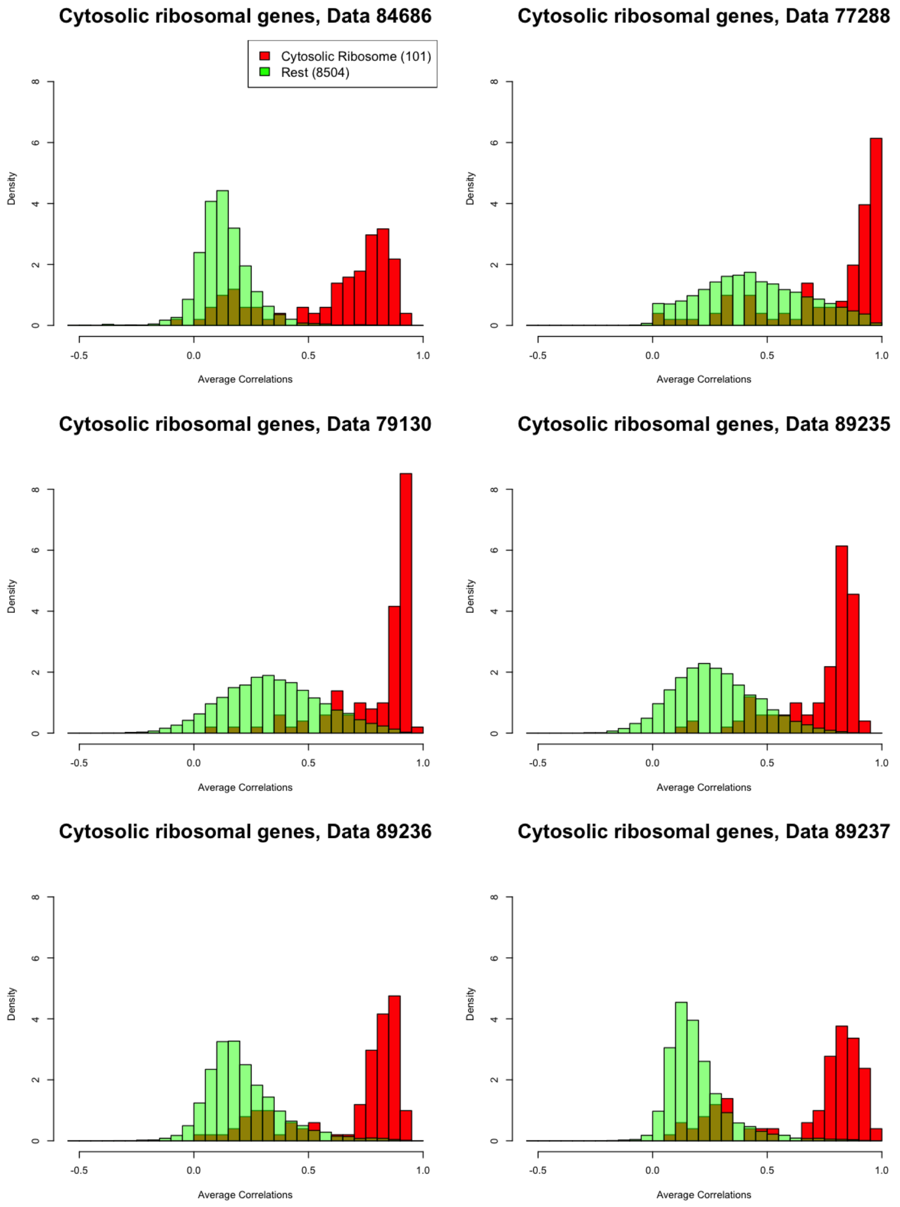


Figure 22: Histogram of the correlations of each gene with the adjusted cell total for cytosolic ribosomal genes for each dataset,, with comparisons between the cytosolic ribosomal genes and the remaining genes.

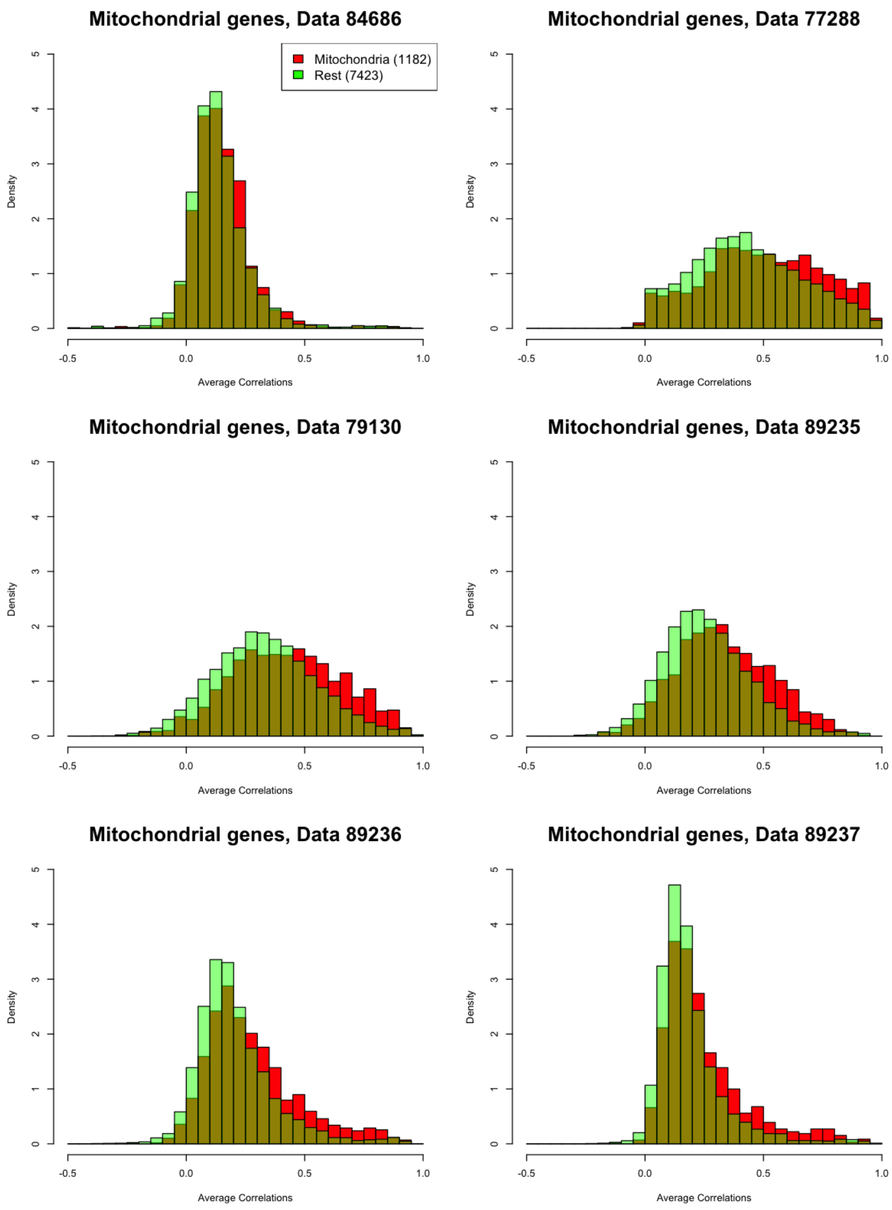


Figure 23: Histogram of the correlations of each gene with the adjusted cell total for mitochondrial genes for each dataset, with comparisons between the mitochondrial genes and the remaining genes.

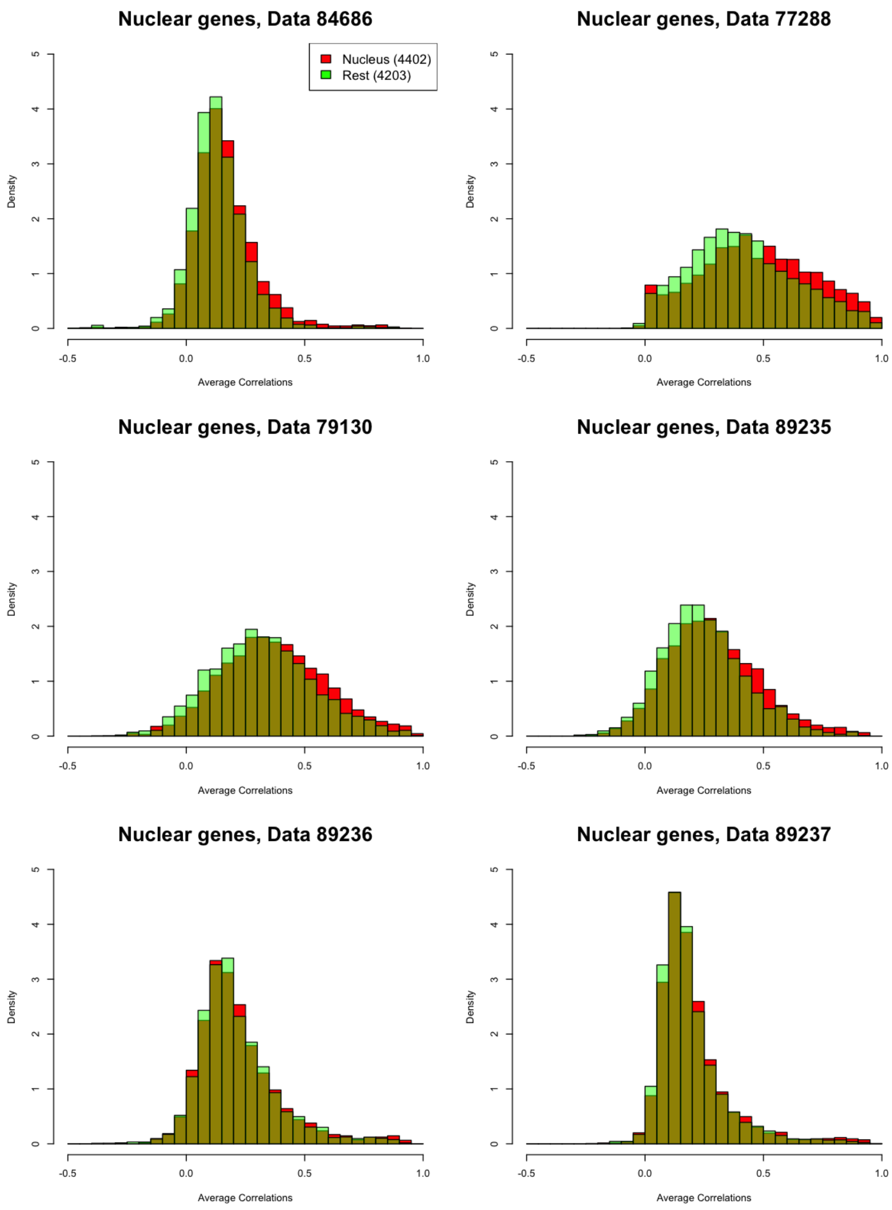


Figure 24: Histogram of the correlations of each gene with the adjusted cell total for nuclear genes for each dataset, with comparisons between the nuclear genes and the remaining genes.

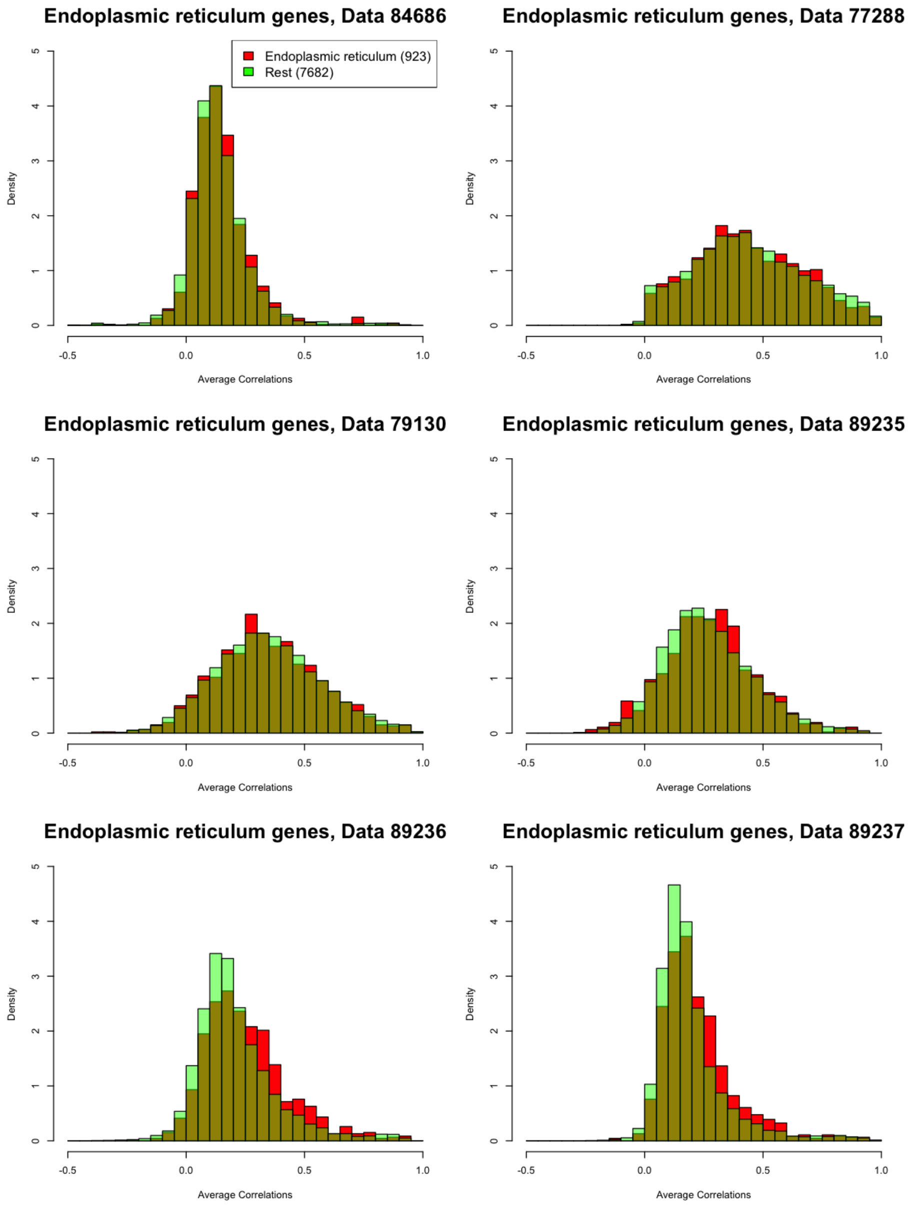


Figure 25: Histogram of the correlations of each gene with the adjusted cell total for endoplasmic reticulum genes for each dataset, with comparisons between the endoplasmic reticulum genes and the remaining genes.

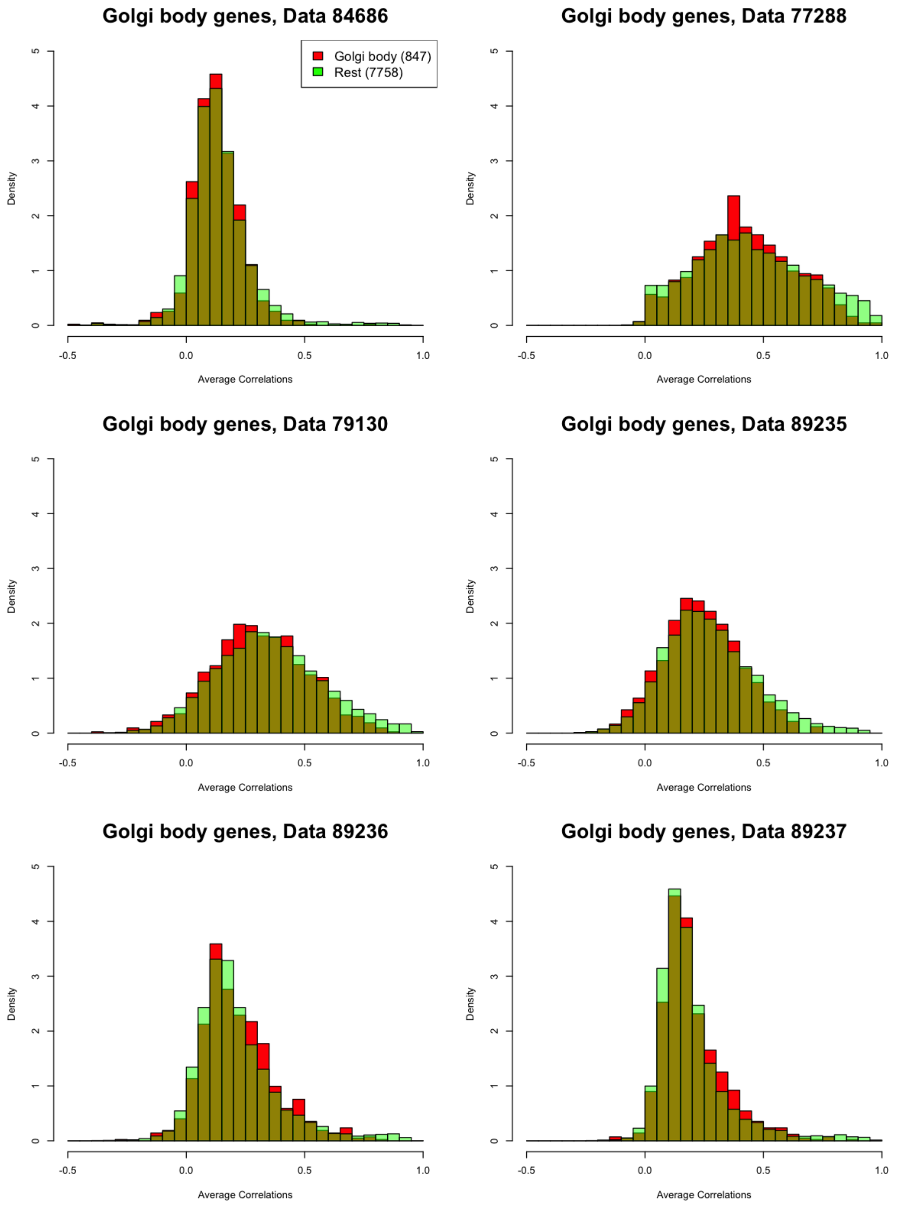


Figure 26: Histogram of the correlations of each gene with the adjusted cell total for Golgi body genes for each dataset, with comparisons between the Golgi body genes and the remaining genes.

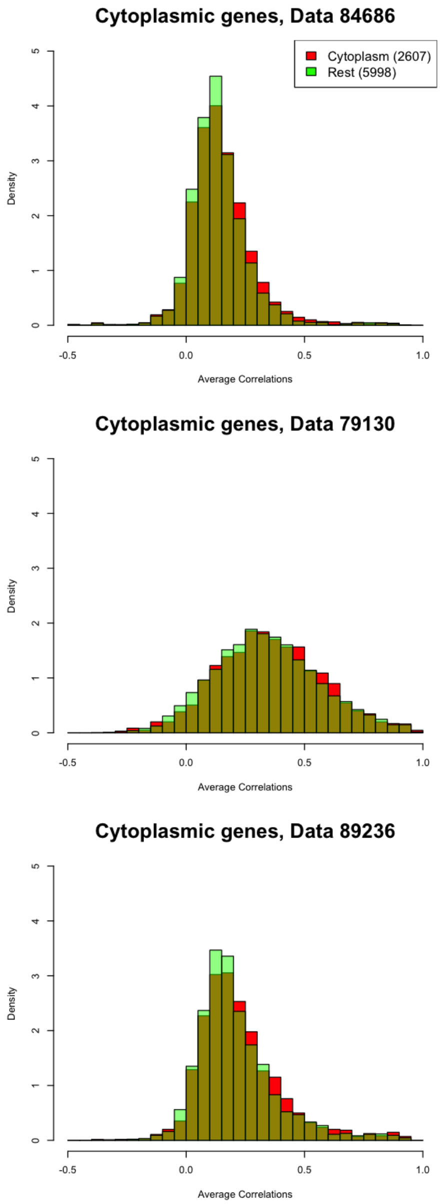


Figure 27: Histogram of the correlations of each gene with the adjusted cell total for cytoplasmic genes for each dataset, with comparisons between the cytoplasmic genes and the remaining genes.

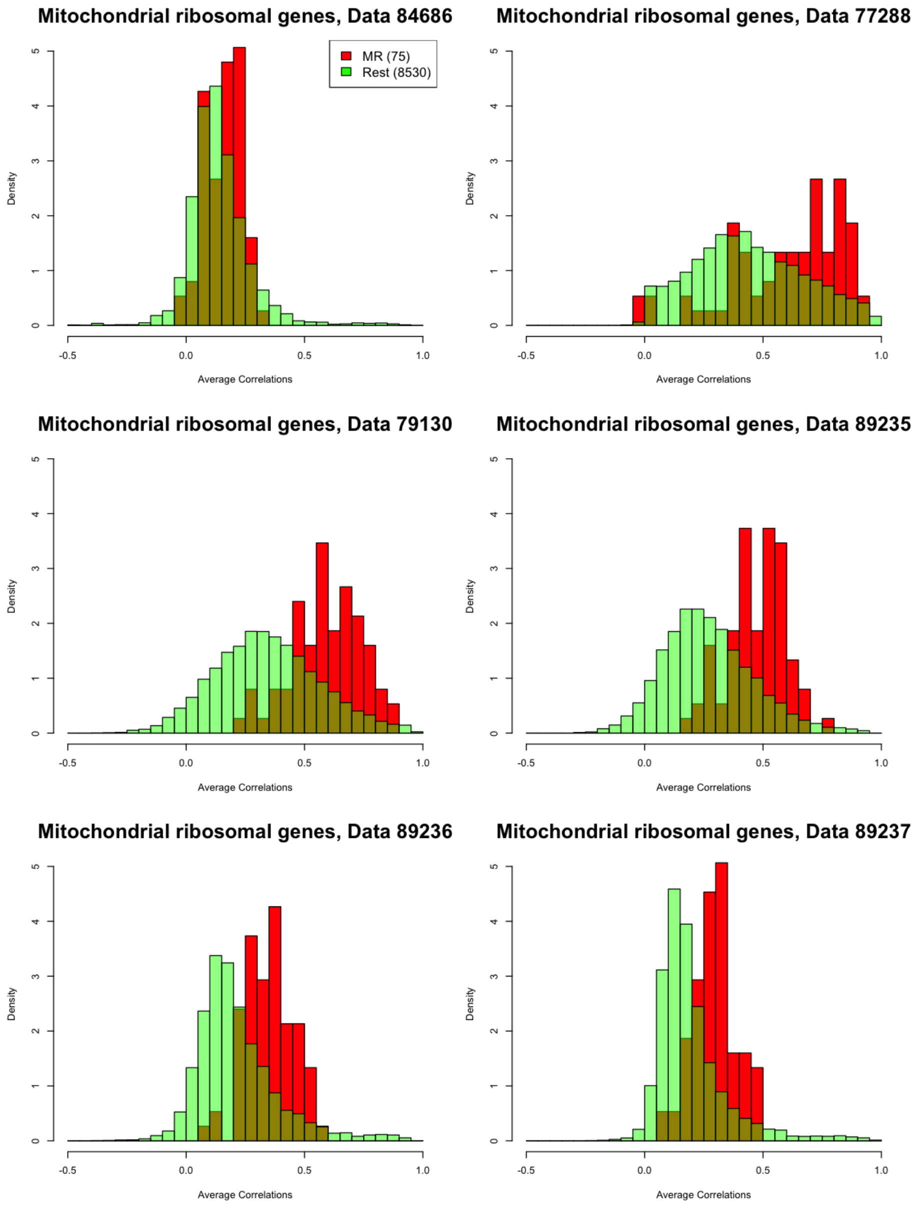


Figure 28: Histogram of the correlations of each gene with the adjusted cell total for mitochondrial ribosomal genes for each dataset, with comparisons between the mitochondrial ribosomal genes and the remaining genes.

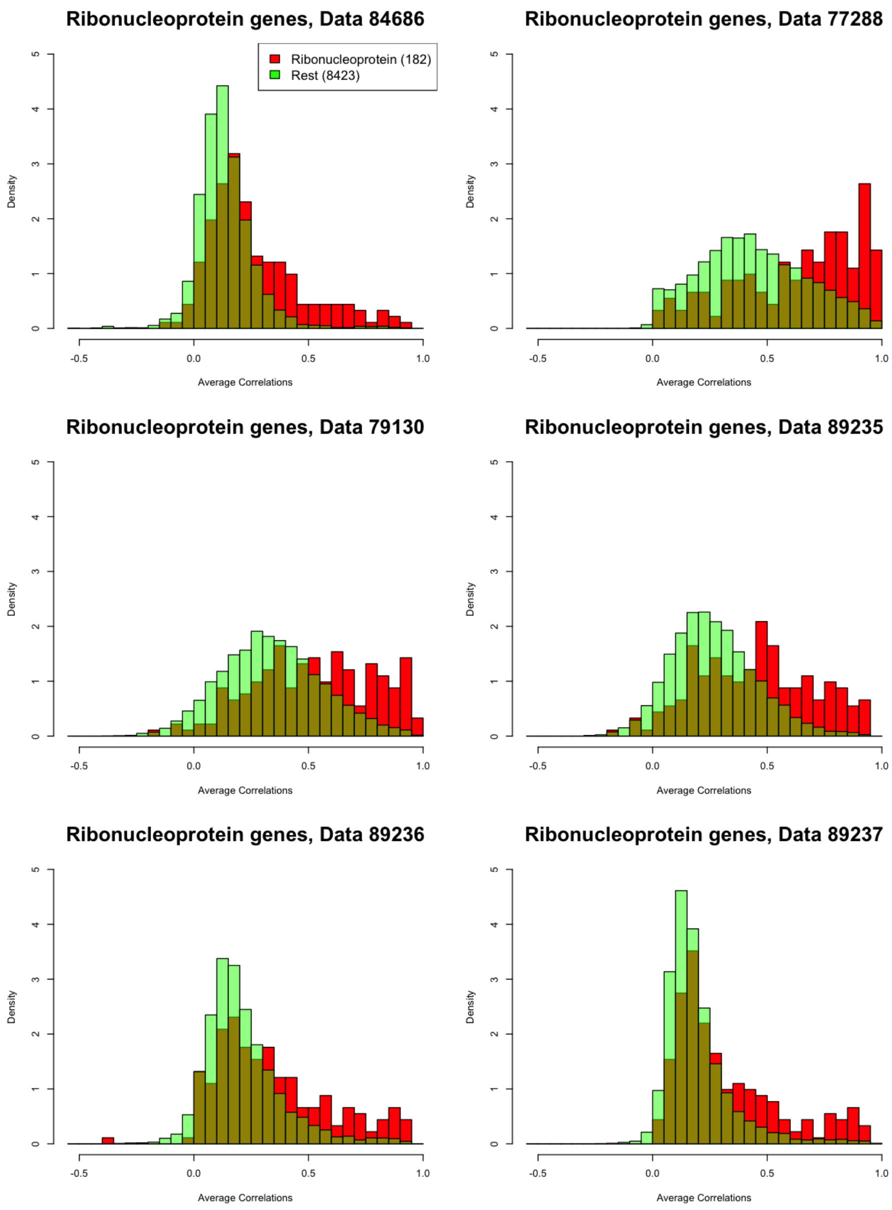


Figure 29: Histogram of the correlations of each gene with the adjusted cell total for ribonucleoprotein genes for each dataset, with comparisons between the ribonucleoprotein genes and the remaining genes.

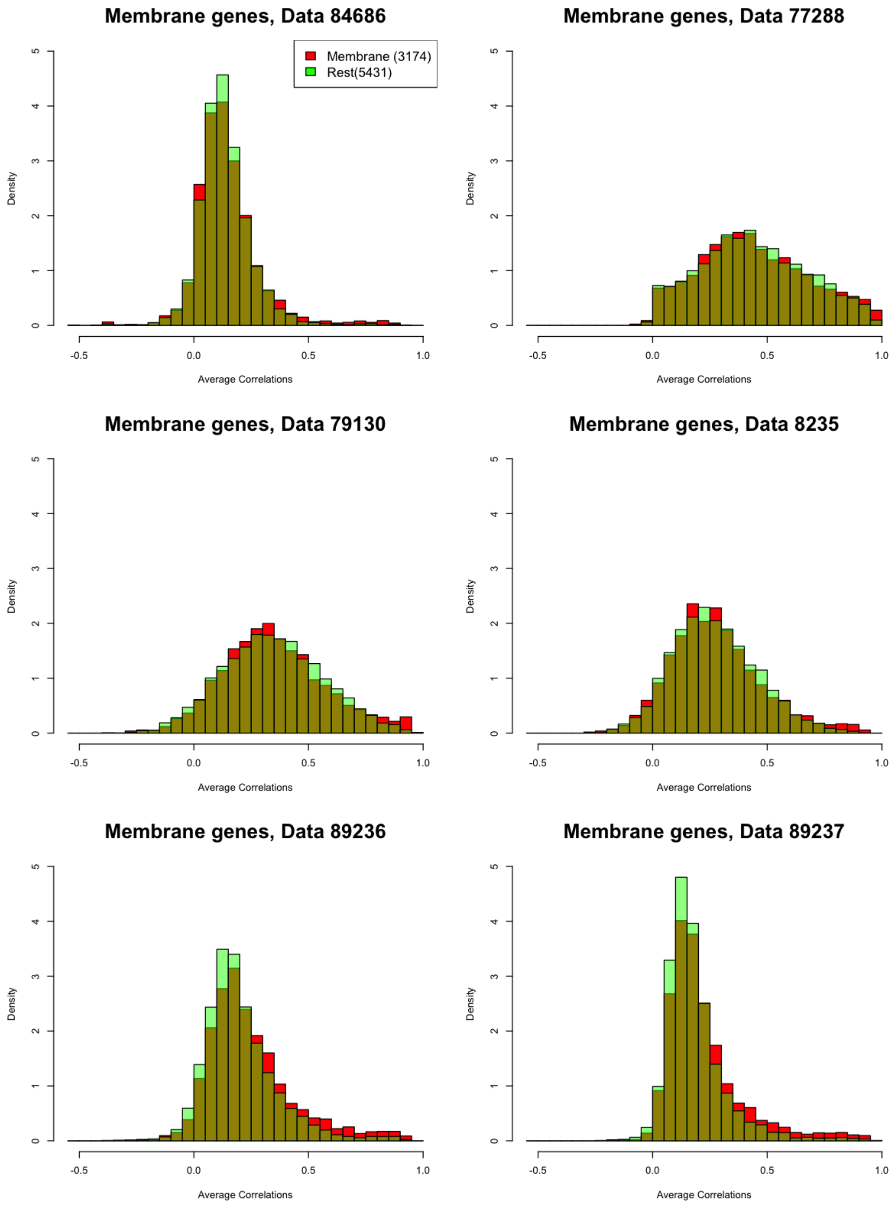


Figure 30: Histogram of the correlations of each gene with the adjusted cell total for membrane genes for each dataset, with comparisons between the membrane genes and the remaining genes.

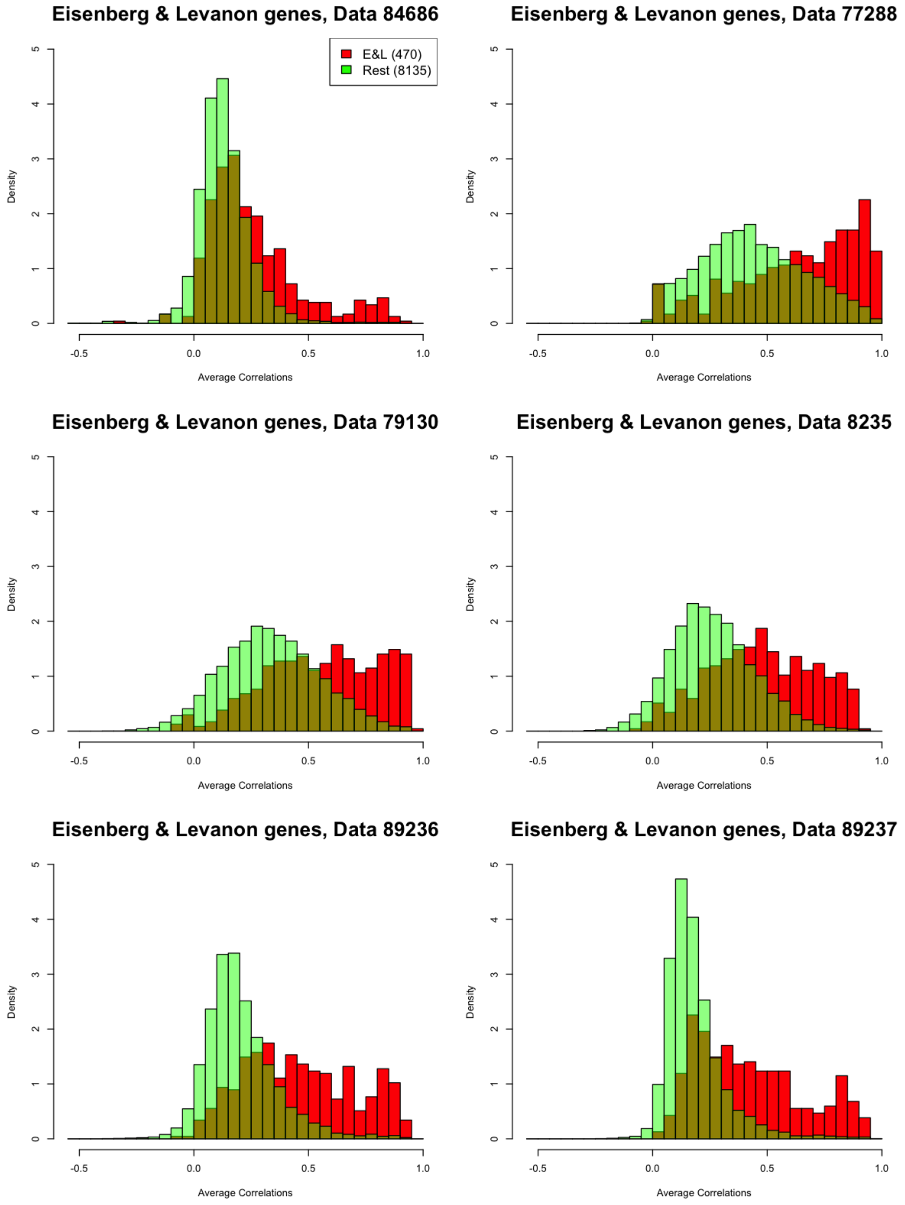


Figure 31: Histogram of the correlations of each gene with the adjusted cell total for stably expressed genes from Eisenberg and Levanon [2013] for each dataset, with comparisons between the stably expressed genes and the remaining genes.

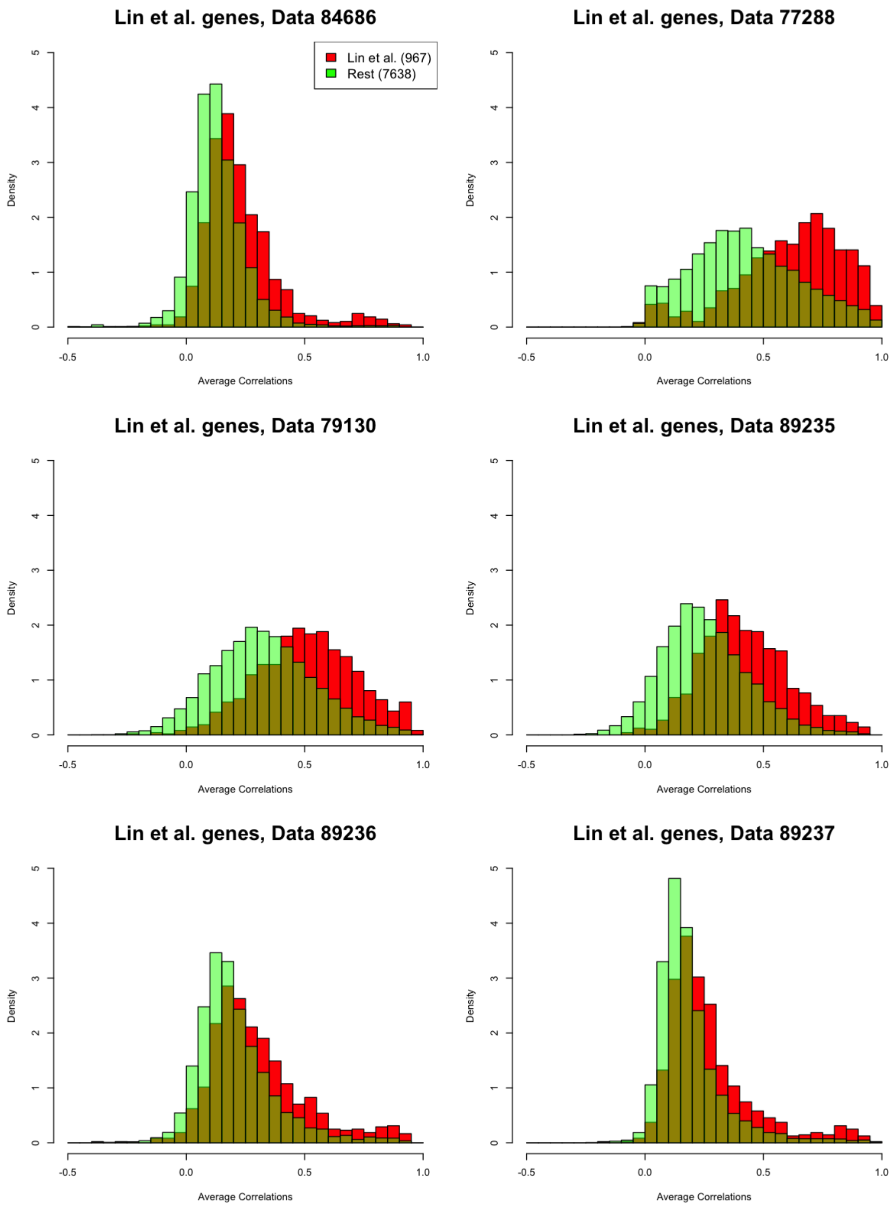


Figure 32: Histogram of the correlations of each gene with the adjusted cell total for single cell stably expressed genes from Lin *et al.* [2017] for each dataset, with comparisons between the single cell stably expressed genes and the remaining genes.

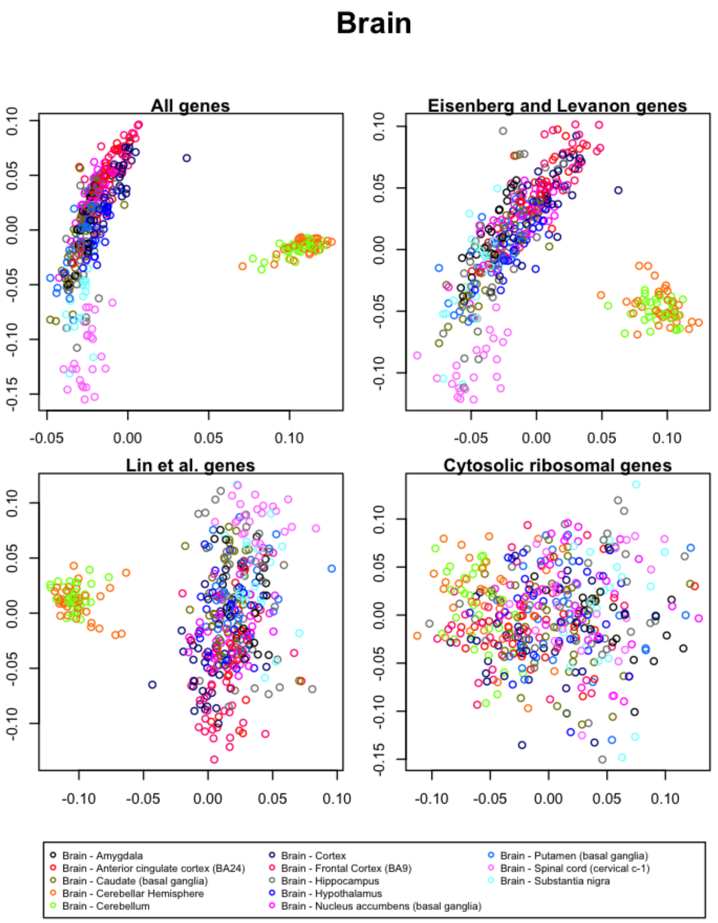


Figure 33: The first and second singular vectors of the GTEx data from the brain.

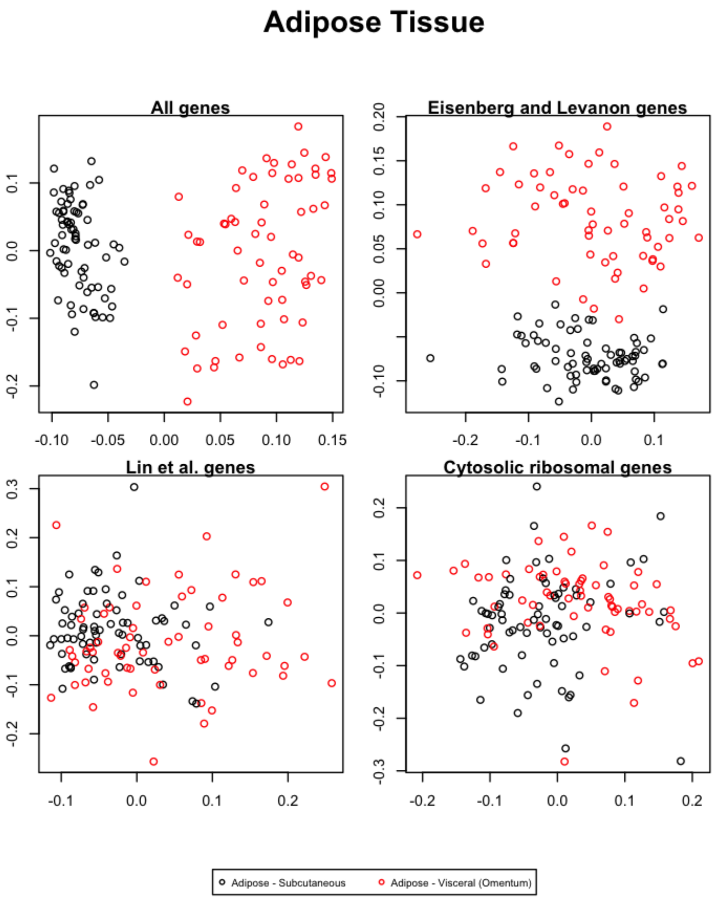


Figure 34: The first and second singular vectors of the GTEx data from the adipose tissue.

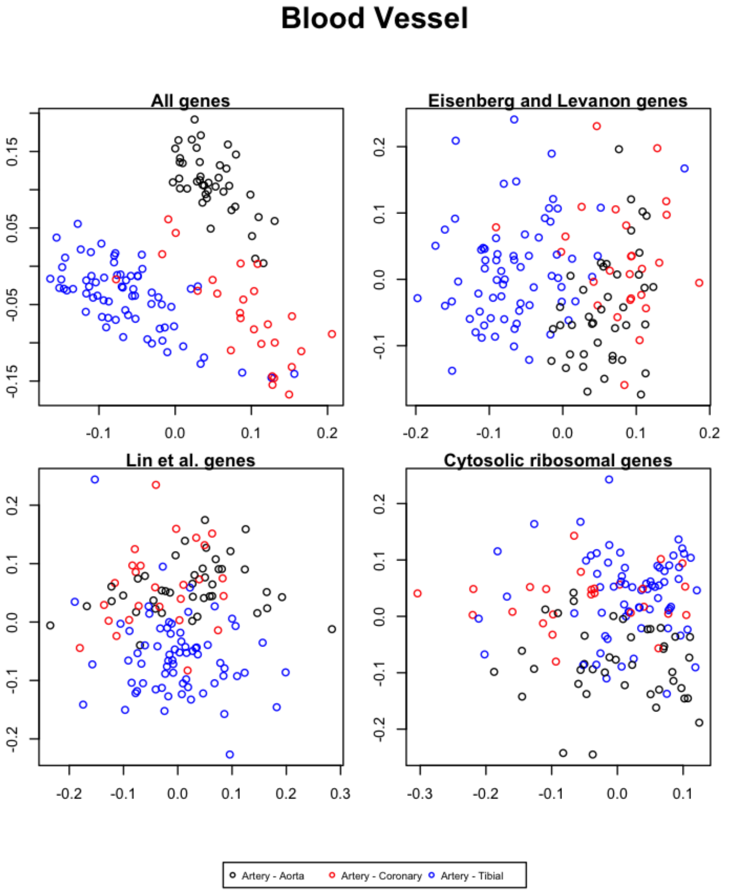


Figure 35: The first and second singular vectors of the GTEx data from the blood vessels.

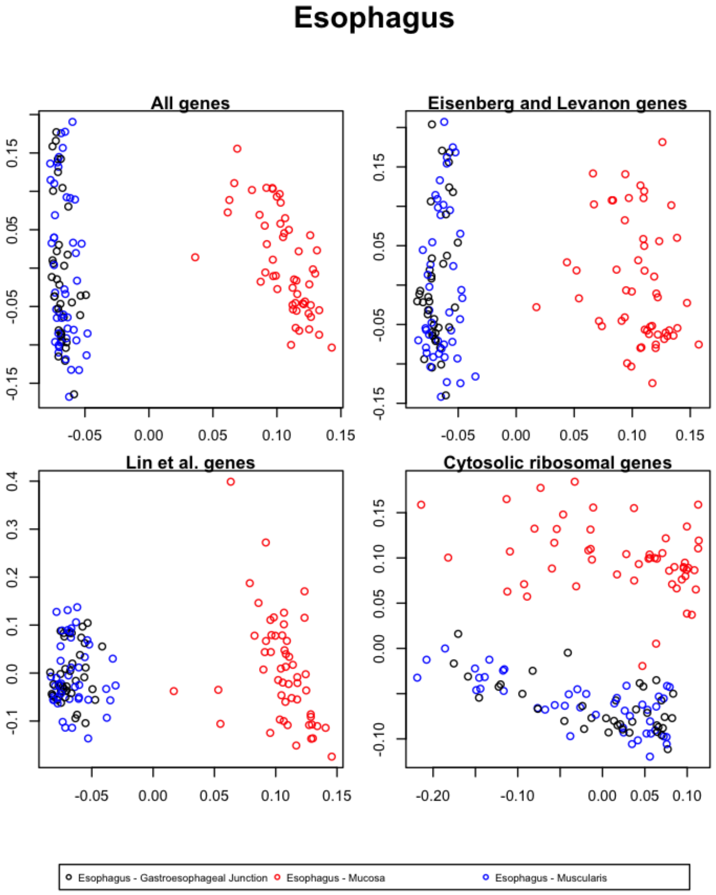


Figure 36: The first and second singular vectors of the GTEx data from the esophagus.

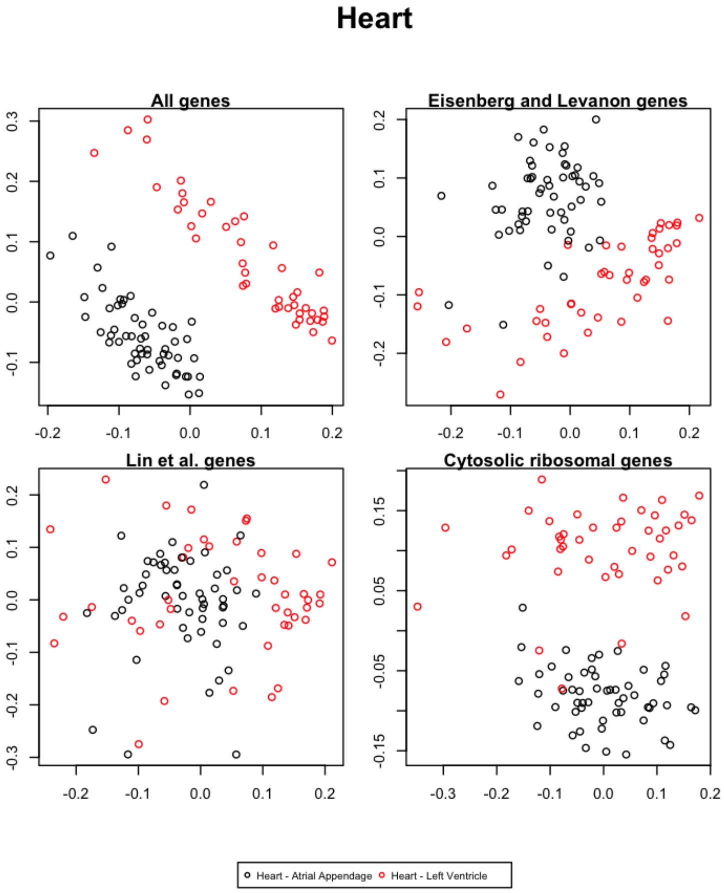


Figure 37: The first and second singular vectors of the GTEx data from the heart.

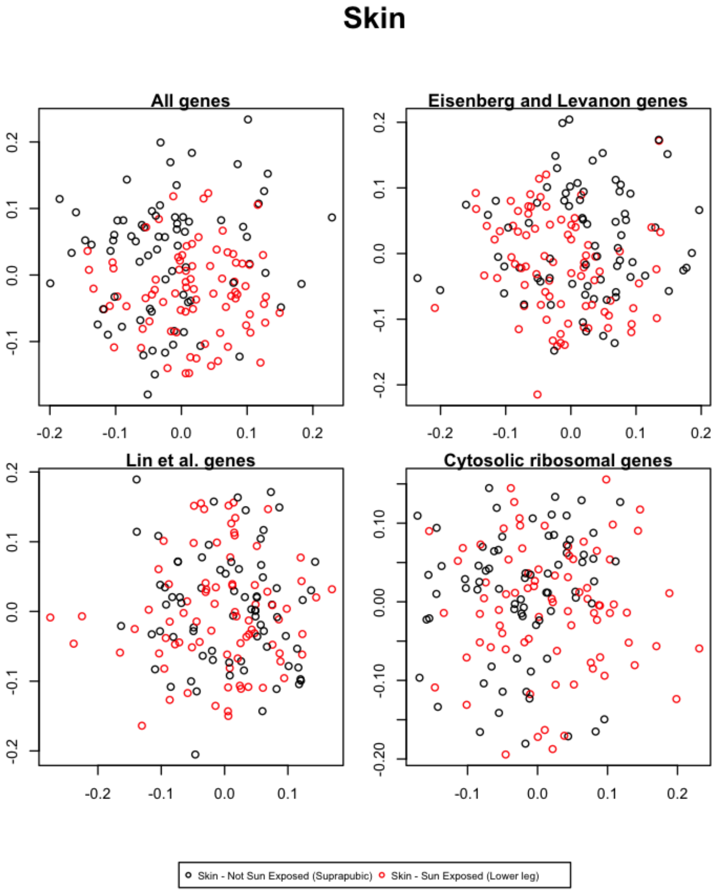


Figure 38: The first and second singular vectors of the GTEx data from the skin.

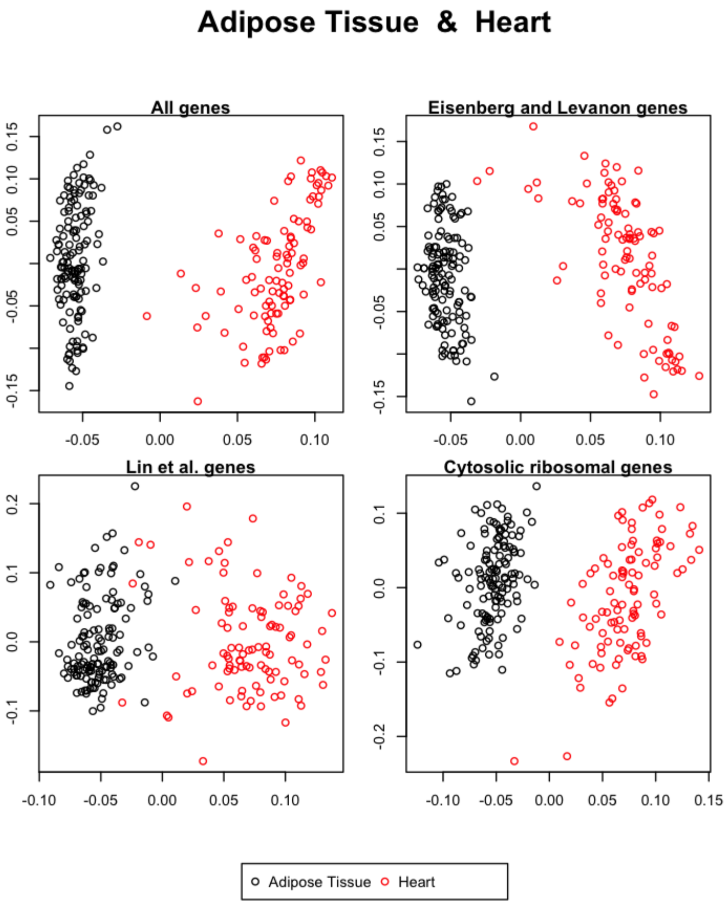


Figure 39: The first and second singular vectors of the GTEx data from the adipose tissue and the heart.

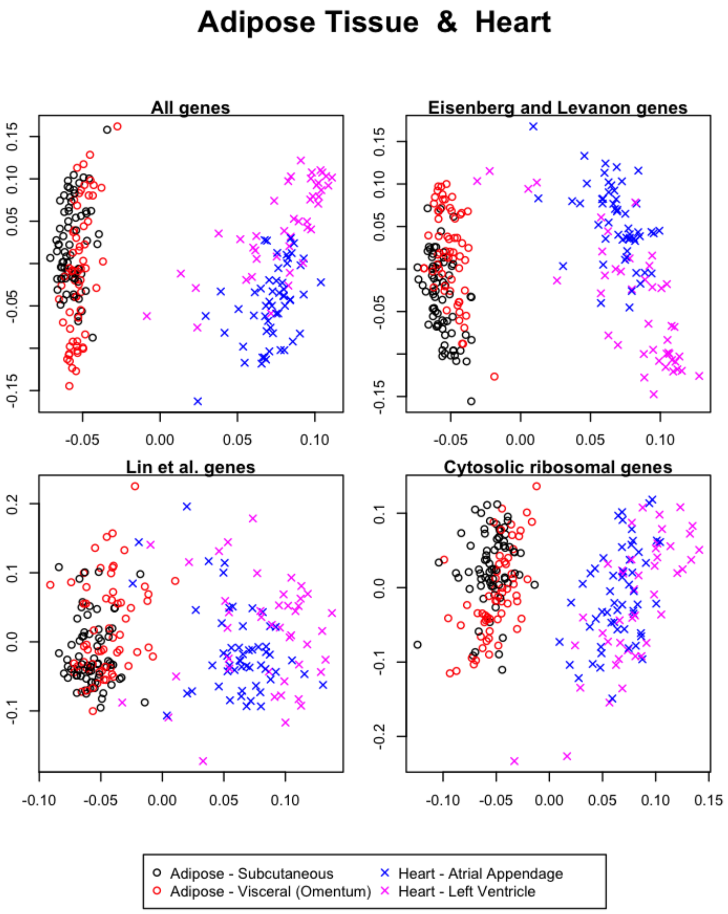


Figure 40: The first and second singular vectors of the GTEx data from the adipose tissue and the heart.

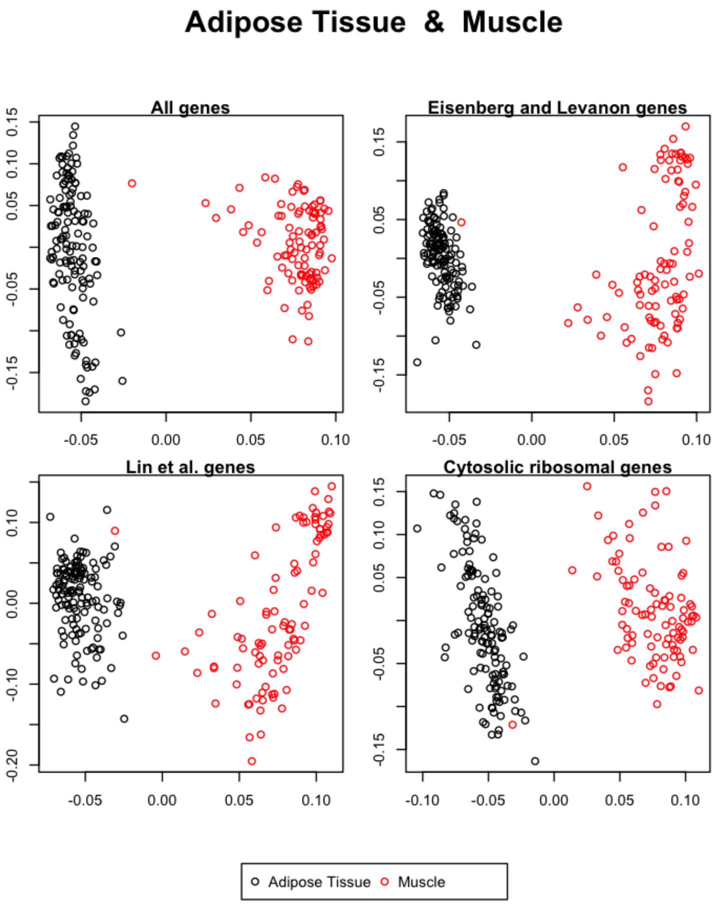


Figure 41: The first and second singular vectors of the GTEx data from the adipose tissue and the muscle.

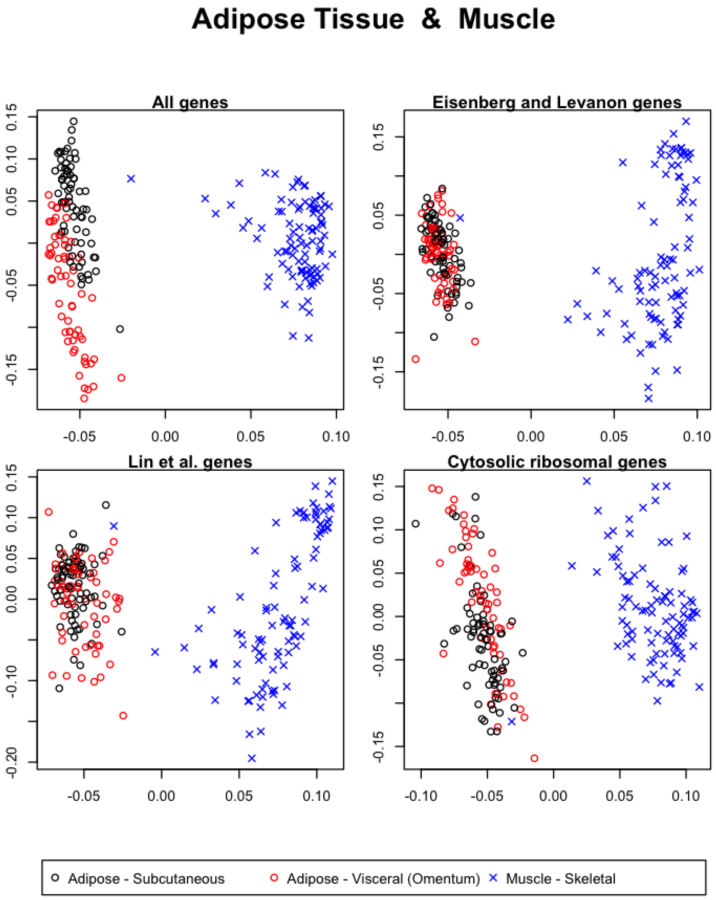


Figure 42: The first and second singular vectors of the GTEx data from the adipose tissue and the muscle.

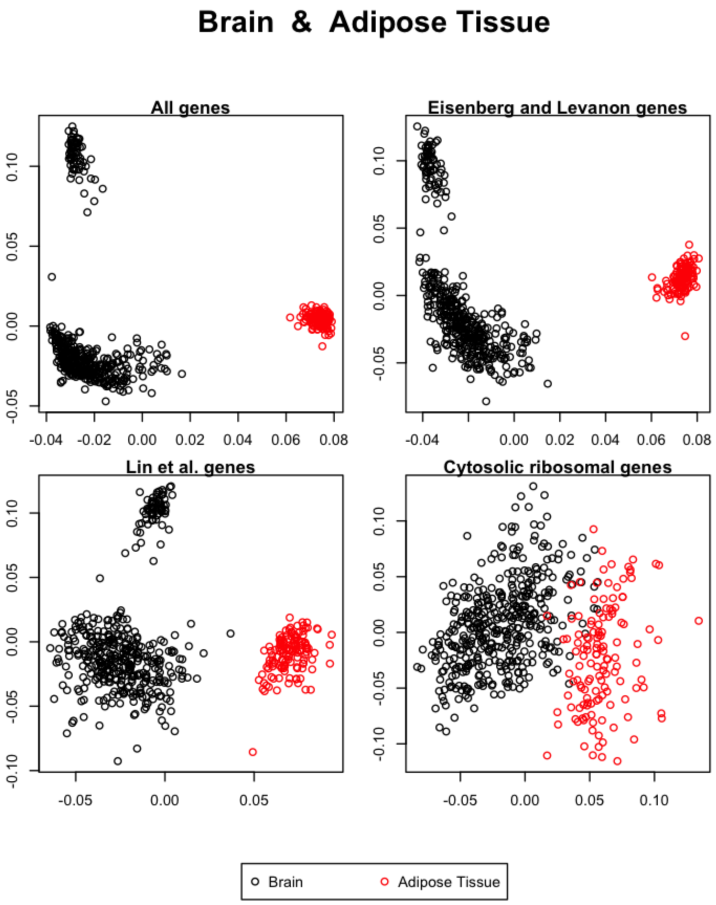


Figure 43: The first and second singular vectors of the GTEx data from the brain and the adipose tissue.

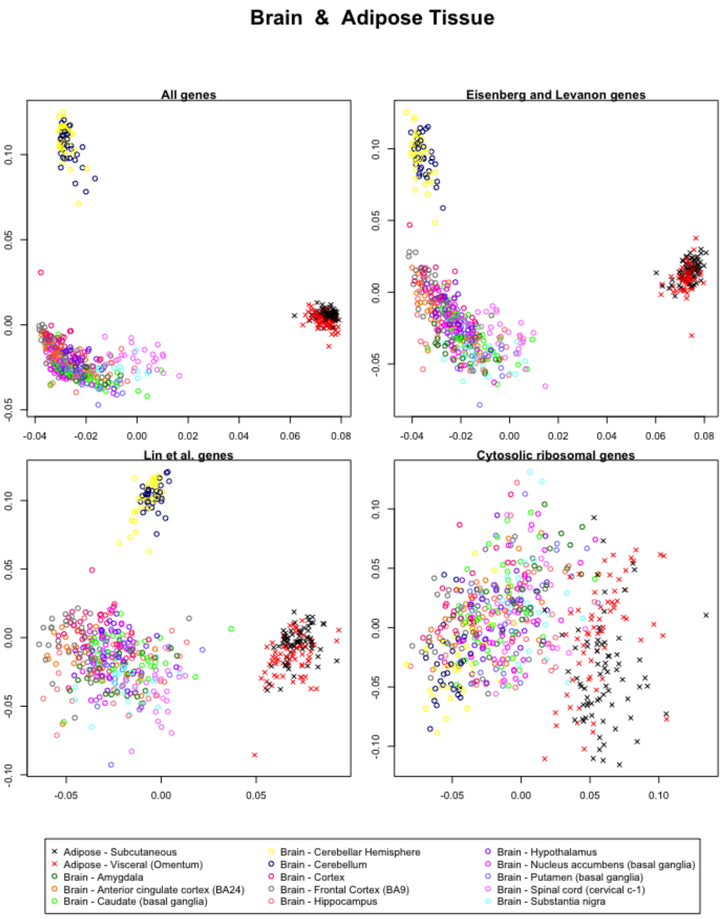


Figure 44: The first and second singular vectors of the GTEx data from the brain and the adipose tissue.

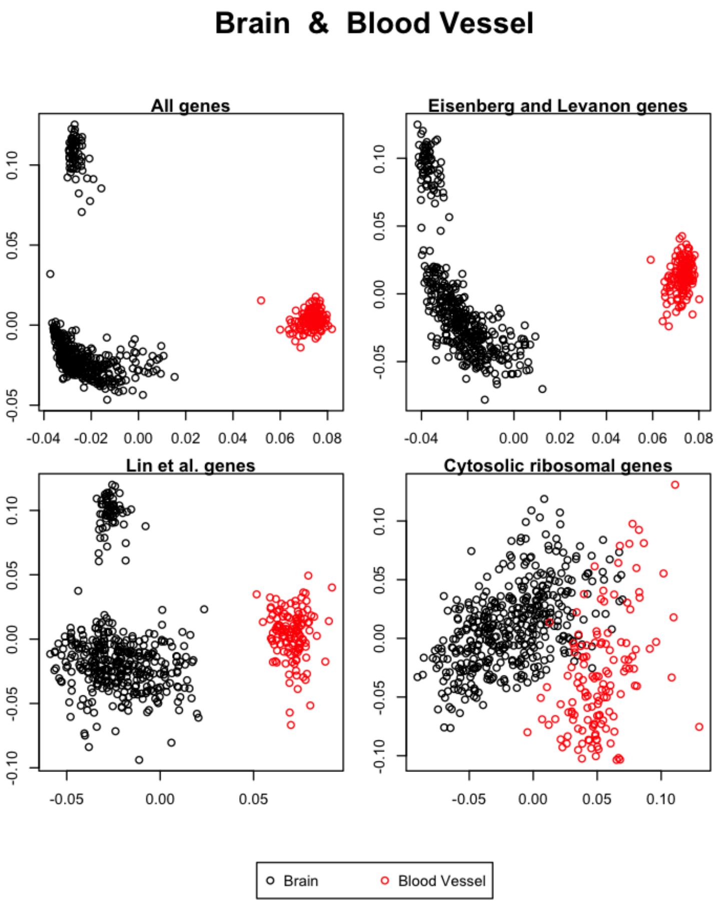


Figure 45: The first and second singular vectors of the GTEx data from the brain and the blood vessel.

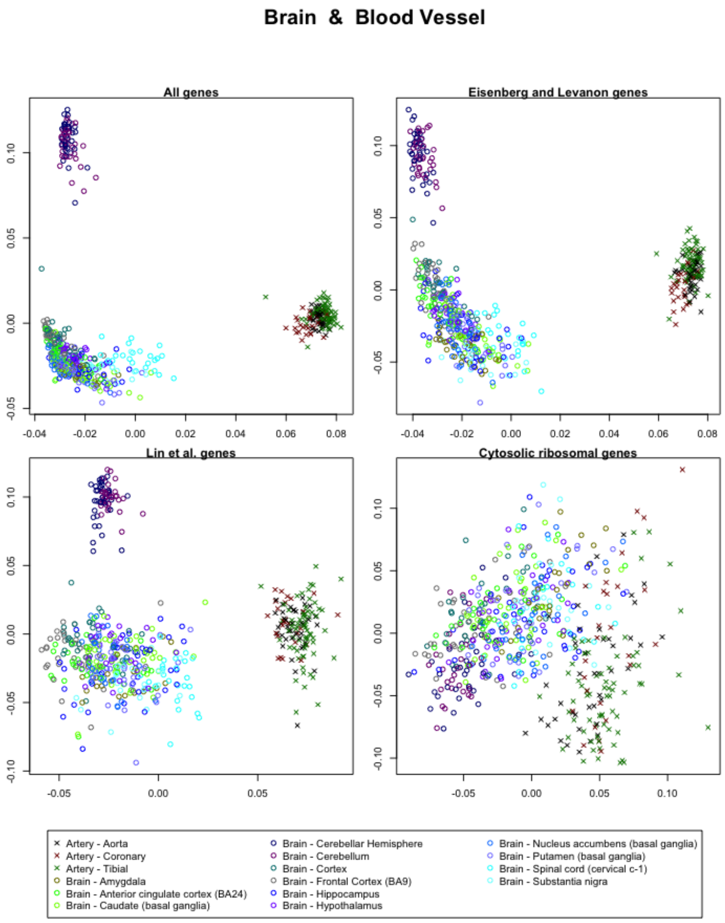


Figure 46: The first and second singular vectors of the GTEx data from the brain and the blood vessel.

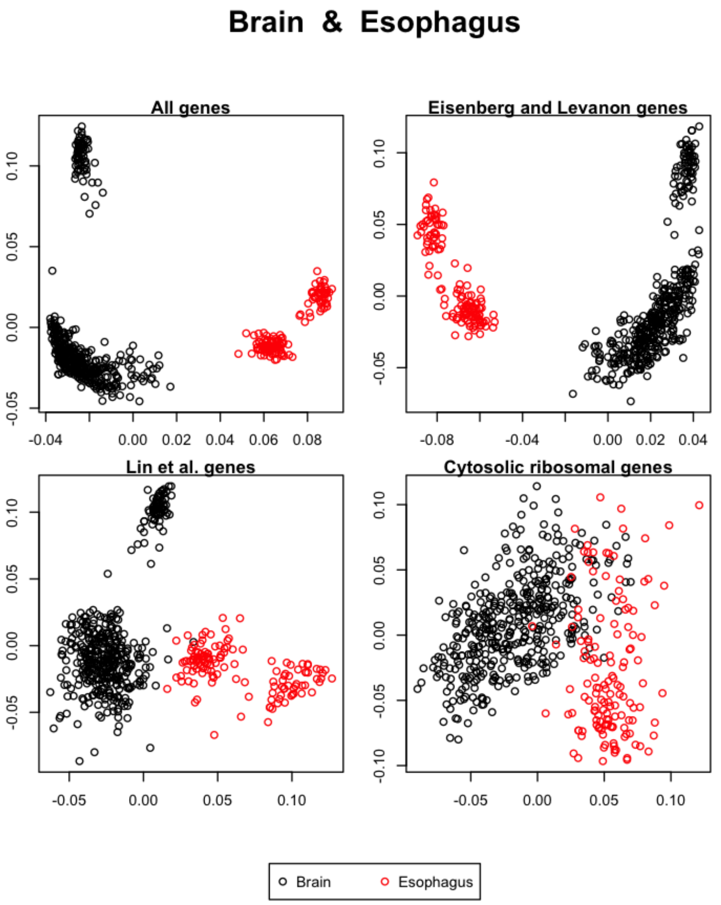


Figure 47: The first and second singular vectors of the GTEx data from the brain and the esophagus.

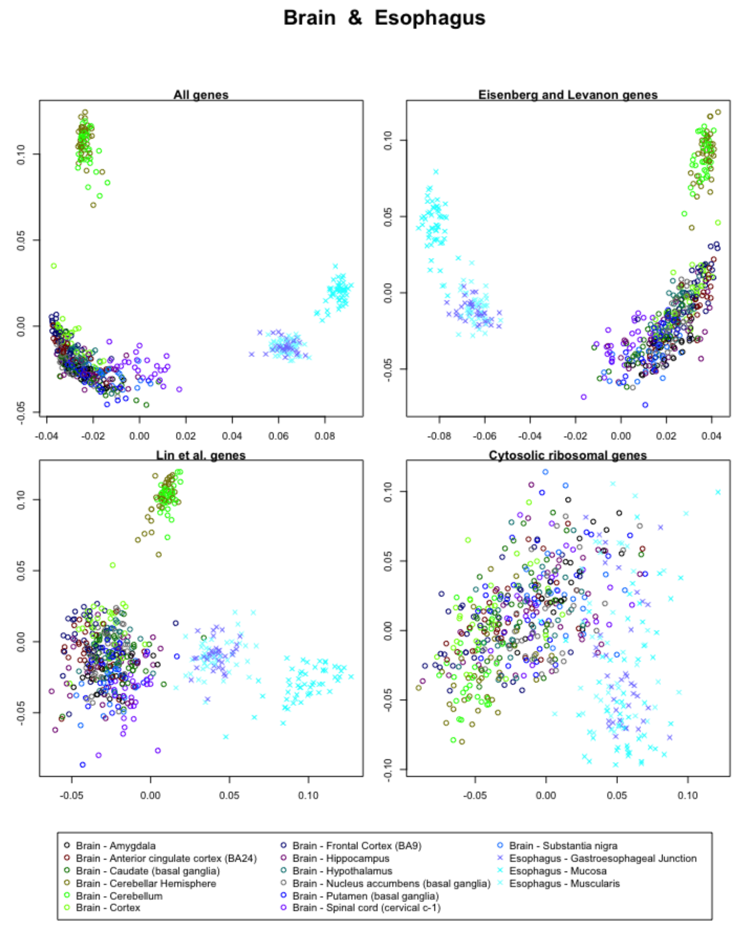


Figure 48: The first and second singular vectors of the GTEx data from the brain and the esophagus.

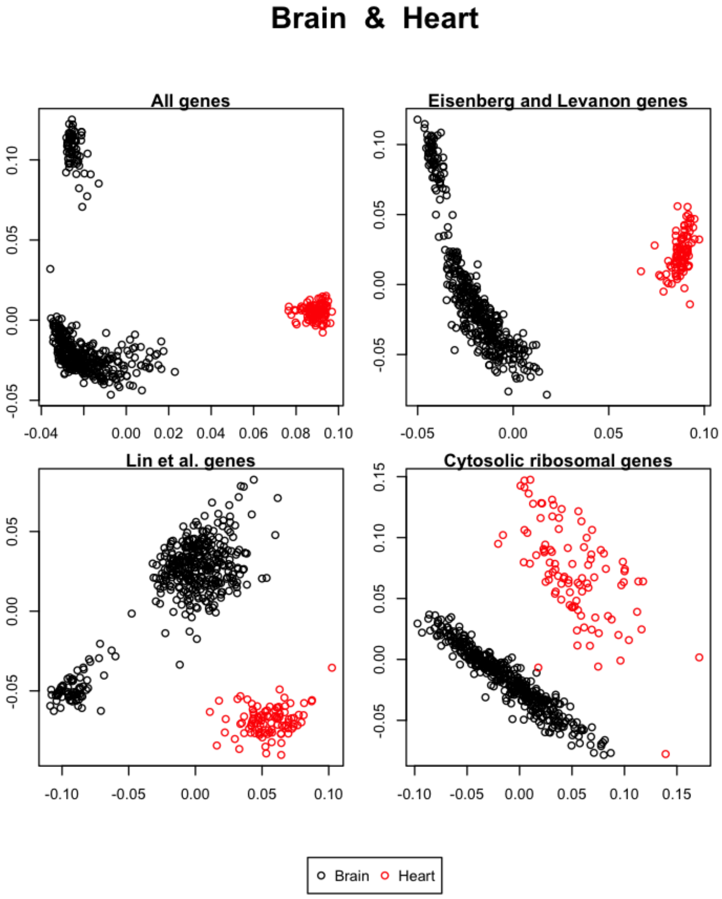


Figure 49: The first and second singular vectors of the GTEx data from the brain and the heart.

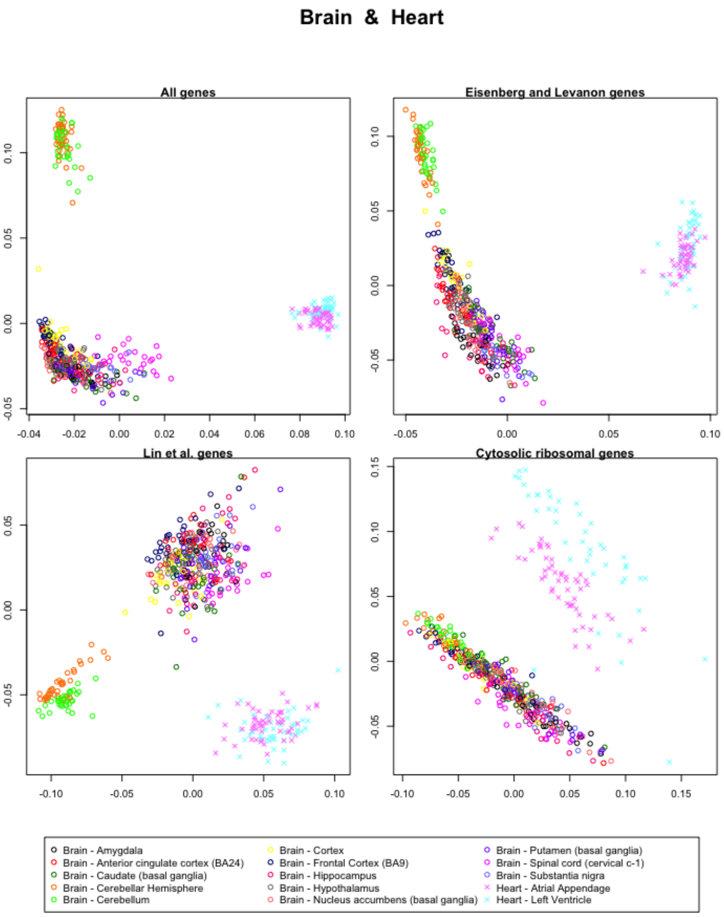


Figure 50: The first and second singular vectors of the GTEx data from the brain and the heart.

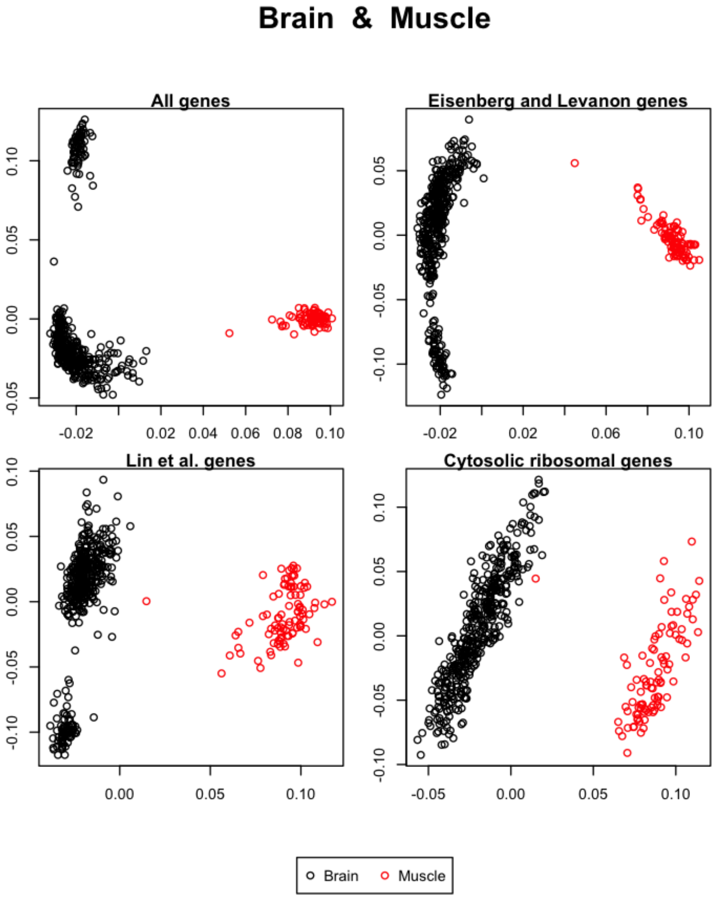


Figure 51: The first and second singular vectors of the GTEx data from the brain and the muscle.

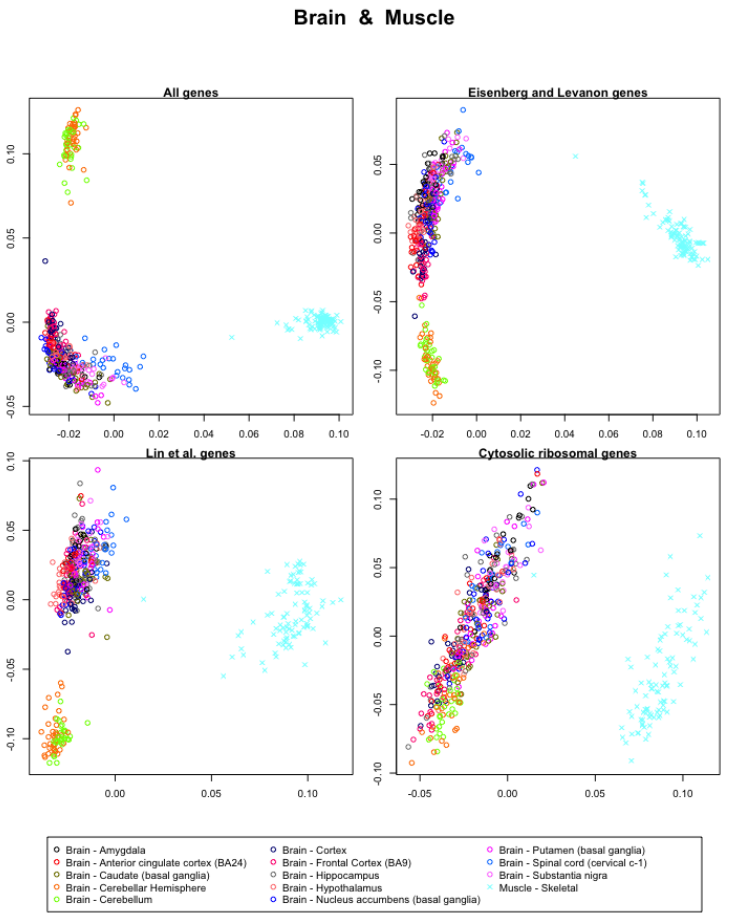


Figure 52: The first and second singular vectors of the GTEx data from the brain and the muscle.

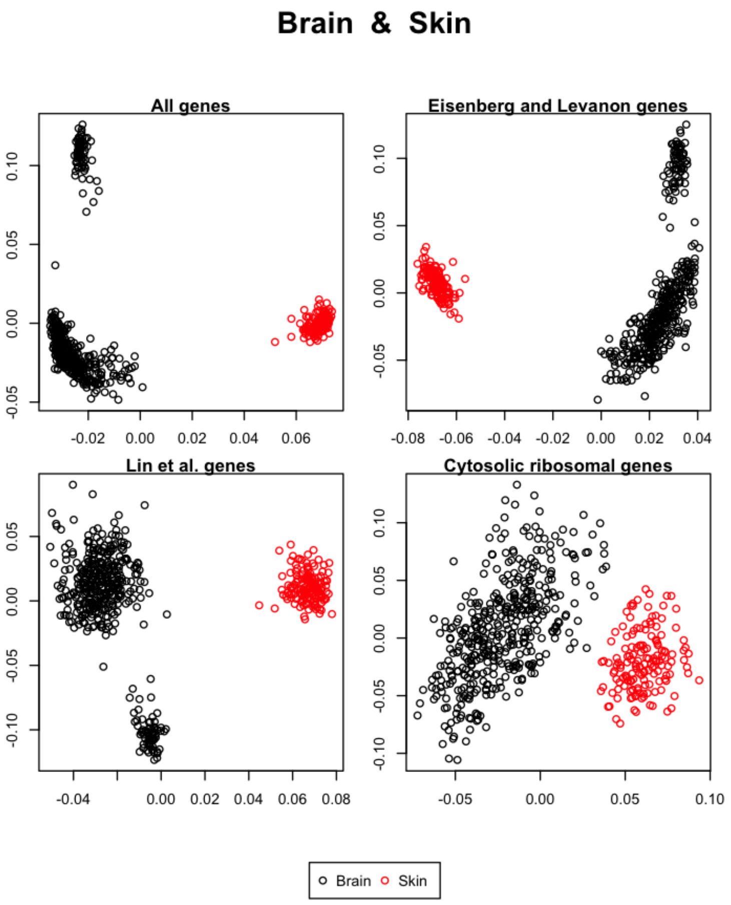


Figure 53: The first and second singular vectors of the GTEx data from the brain and the skin.

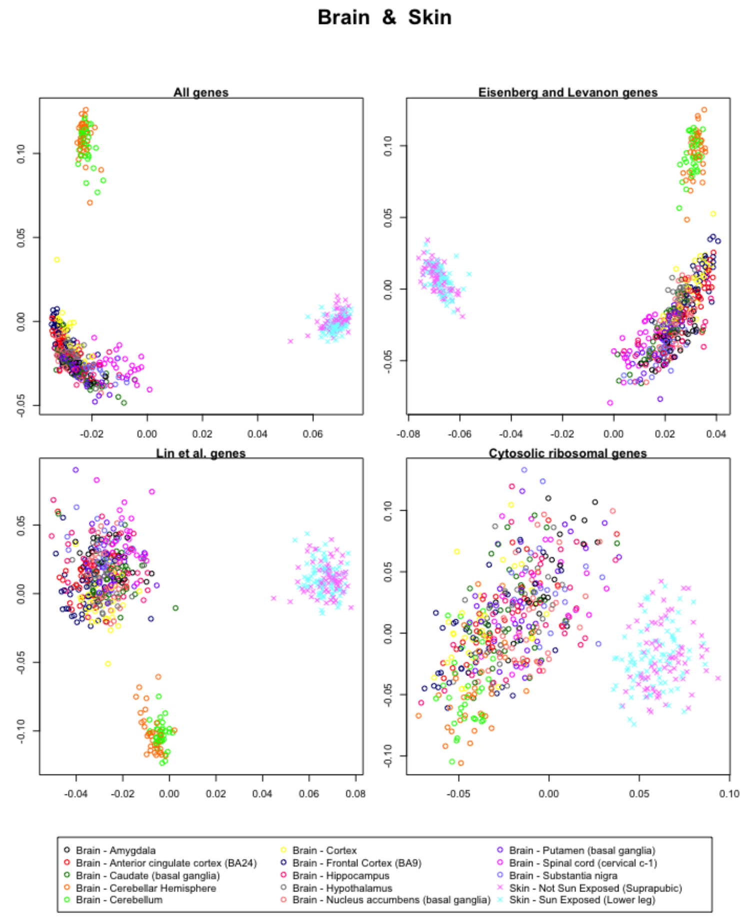


Figure 54: The first and second singular vectors of the GTEx data from the brain and the skin.

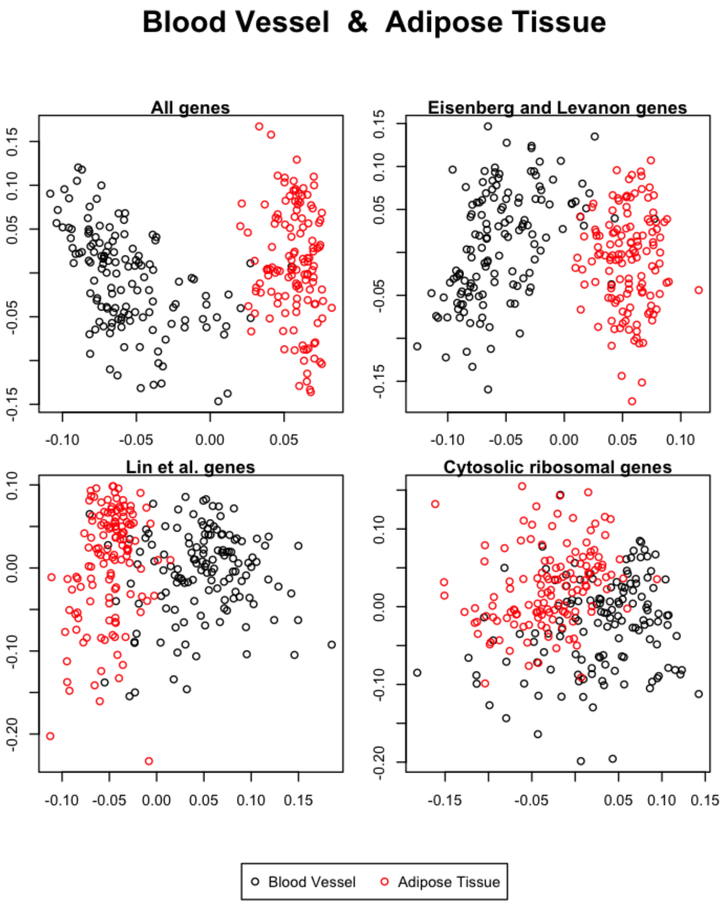


Figure 55: The first and second singular vectors of the GTEx data from the blood vessel and the adipose tissue.

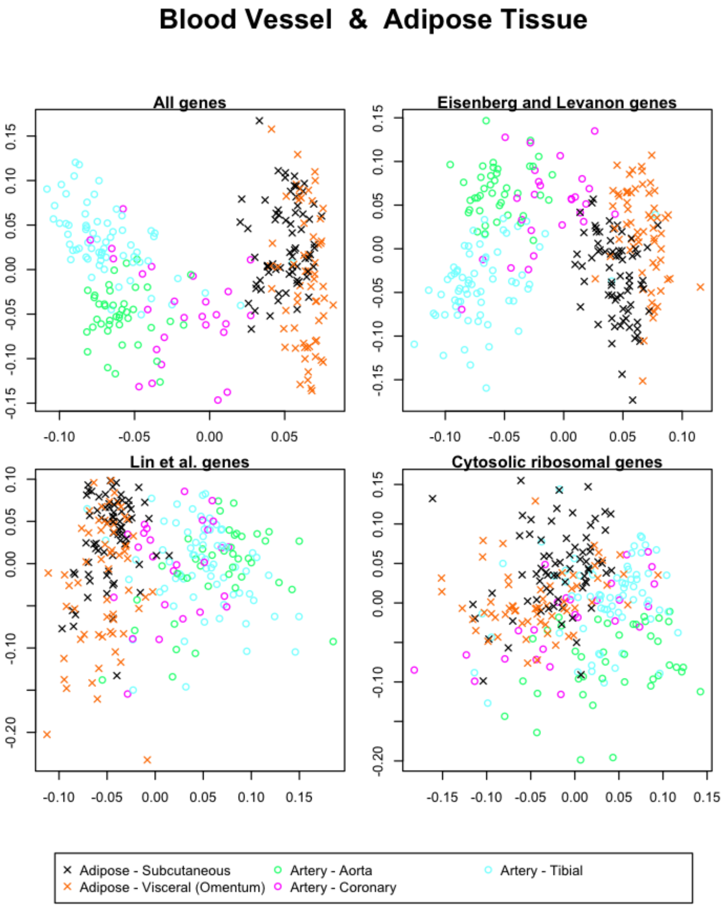


Figure 56: The first and second singular vectors of the GTEx data from the blood vessel and the adipose tissue.

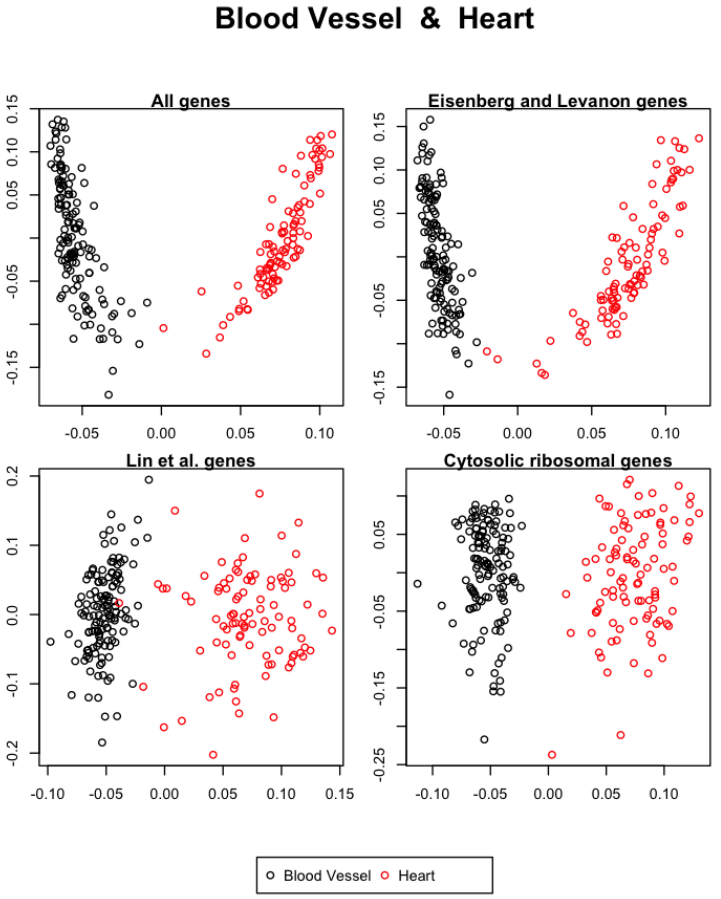


Figure 57: The first and second singular vectors of the GTEx data from the blood vessel and the heart.

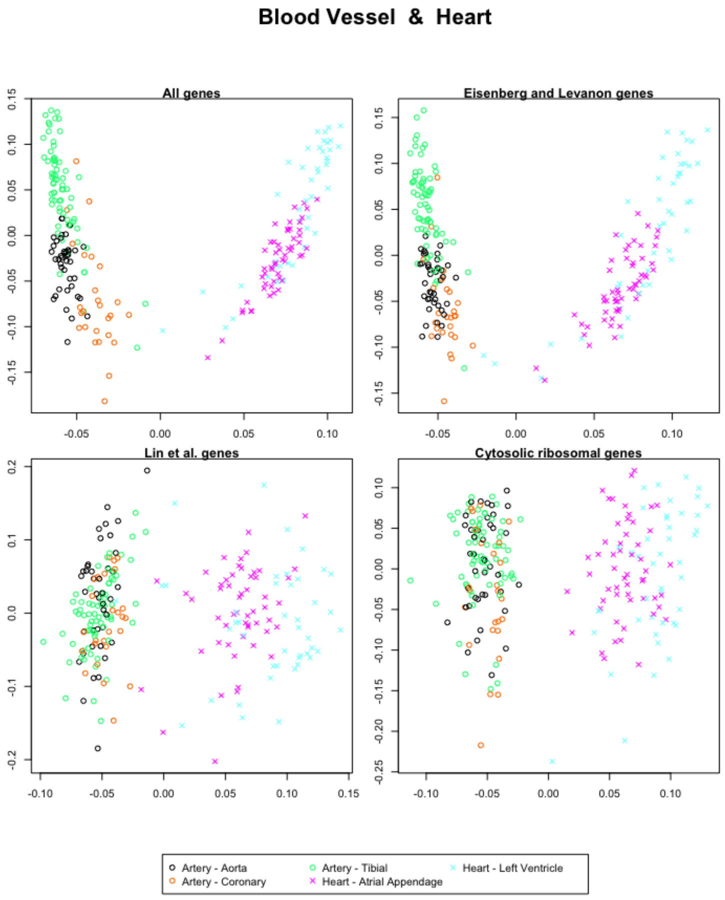


Figure 58: The first and second singular vectors of the GTEx data from the blood vessel and the heart.

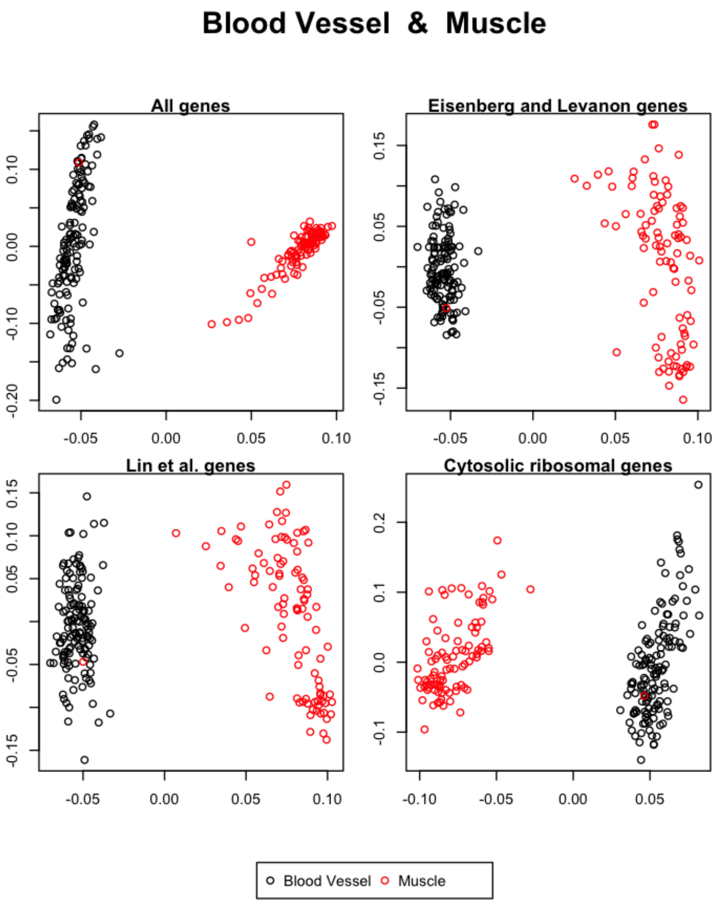


Figure 59: The first and second singular vectors of the GTEx data from the blood vessel and the muscle.

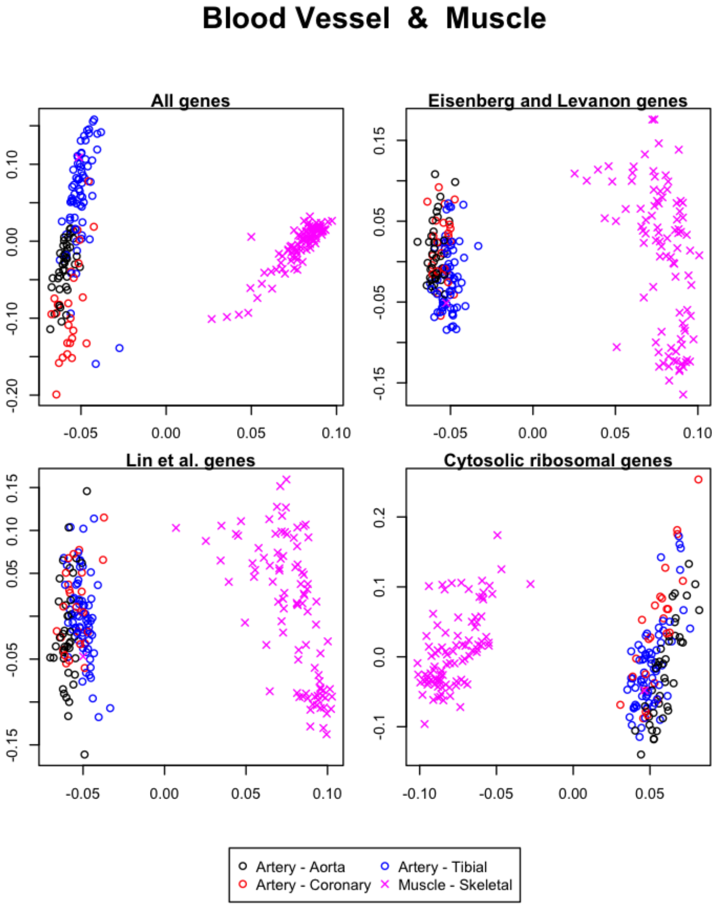


Figure 60: The first and second singular vectors of the GTEx data from the blood vessel and the muscle.

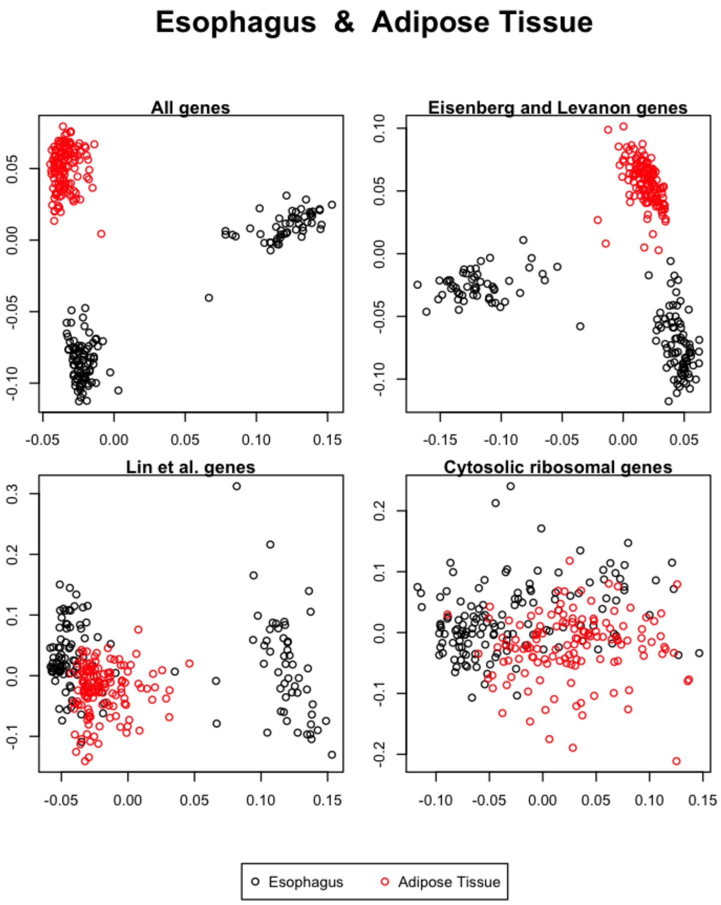


Figure 61: The first and second singular vectors of the GTEx data from the esophagus and the adipose tissue.

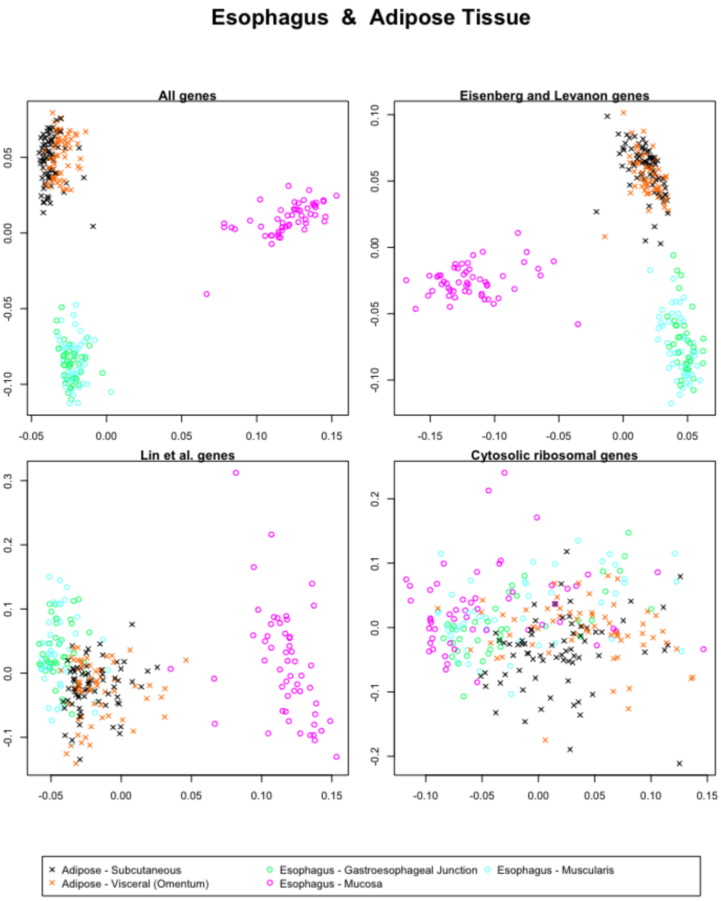


Figure 62: The first and second singular vectors of the GTEx data from the esophagus and the adipose tissue.

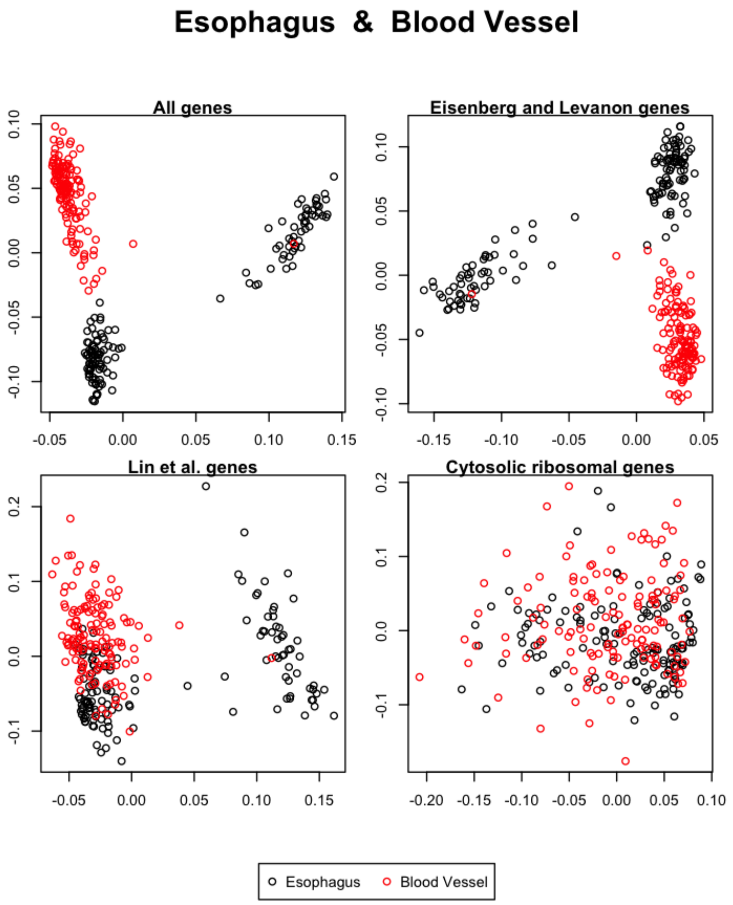


Figure 63: The first and second singular vectors of the GTEx data from the esophagus and the blood vessel.

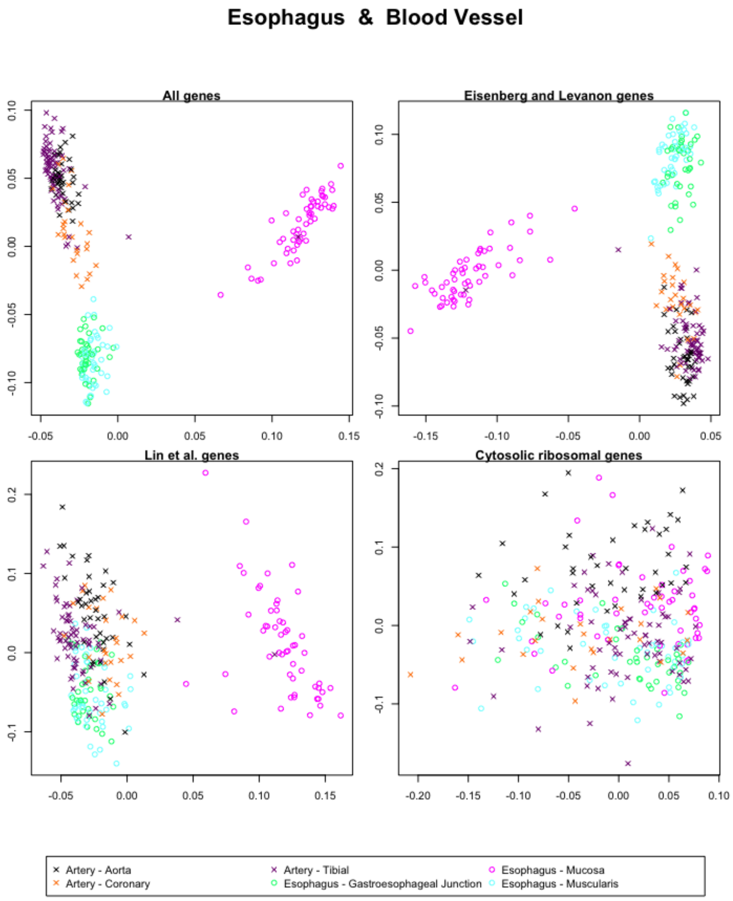


Figure 64: The first and second singular vectors of the GTEx data from the esophagus and the blood vessel.

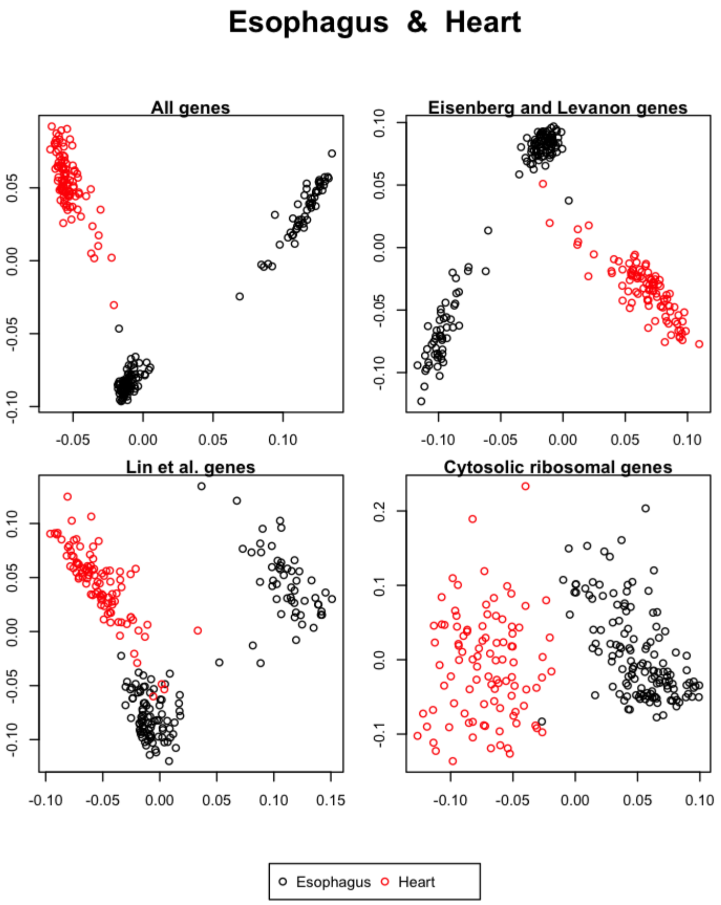


Figure 65: The first and second singular vectors of the GTEx data from the esophagus and the heart.

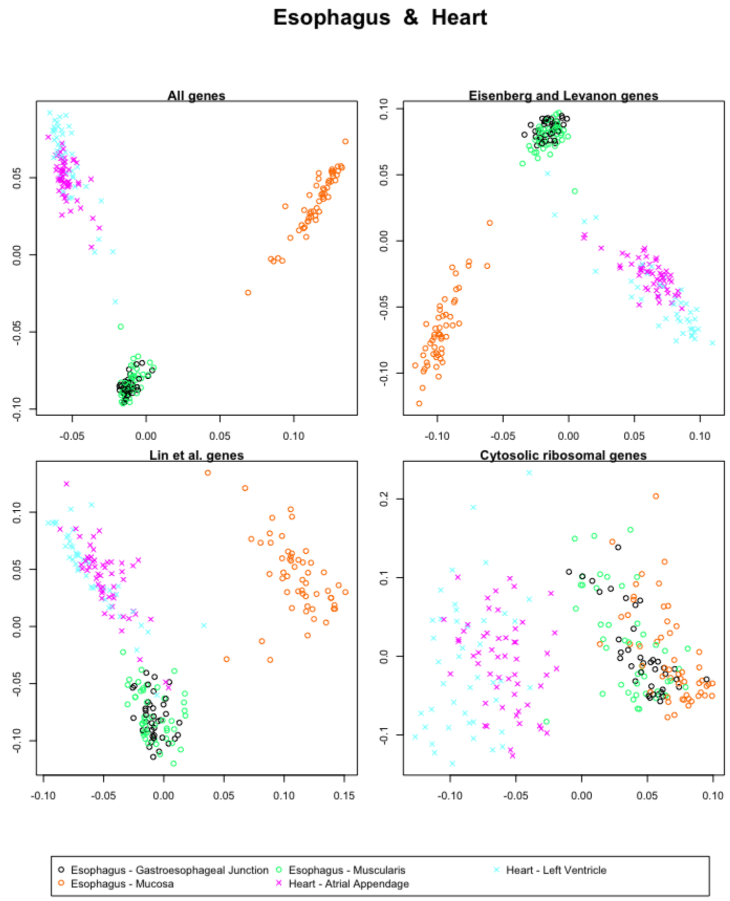


Figure 66: The first and second singular vectors of the GTEx data from the esophagus and the heart.

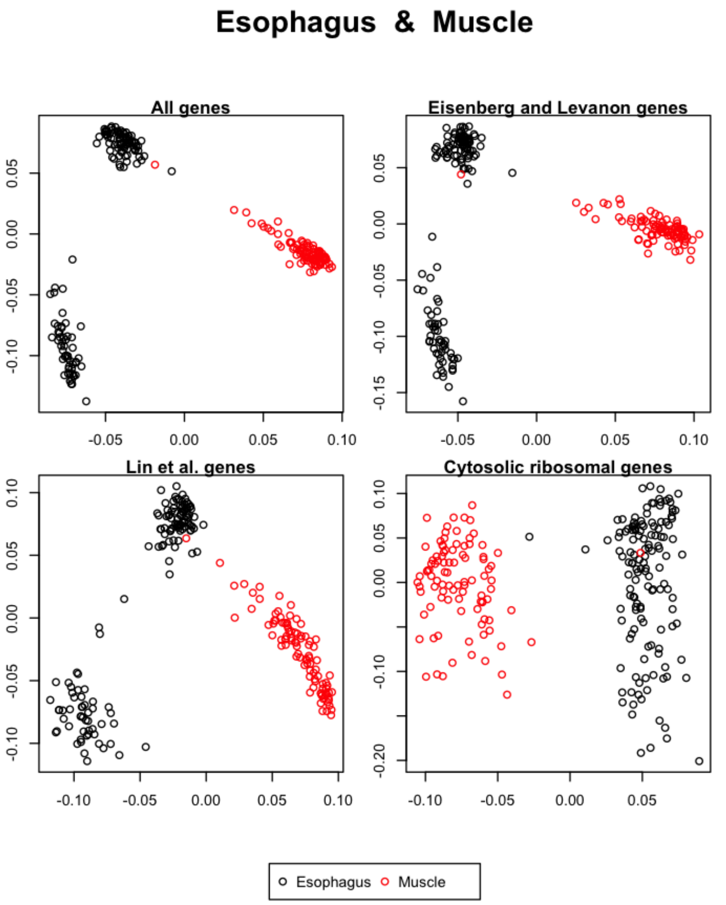


Figure 67: The first and second singular vectors of the GTEx data from the esophagus and the muscle.

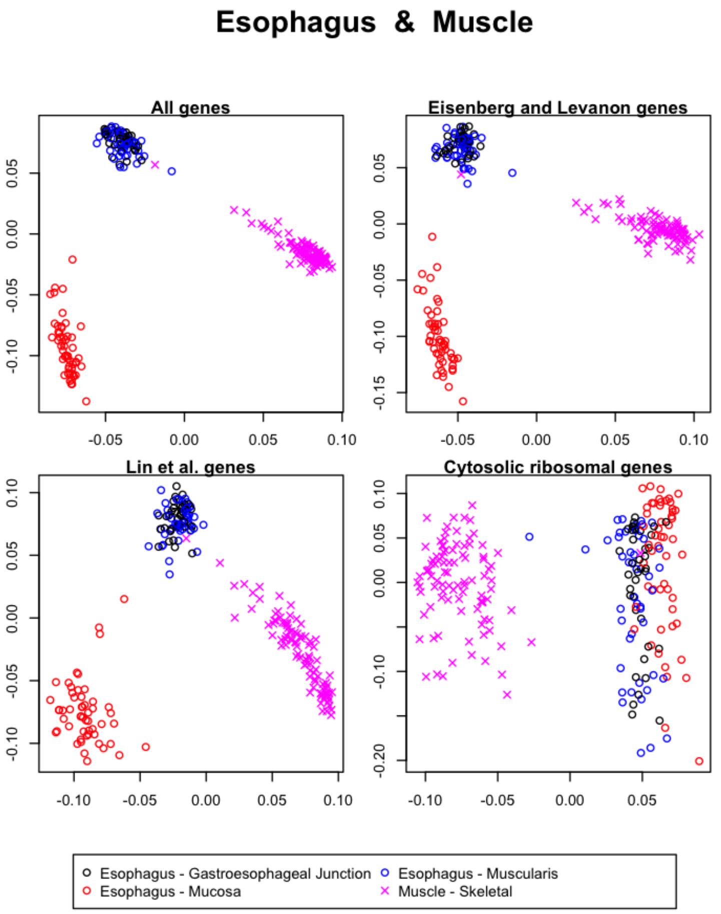


Figure 68: The first and second singular vectors of the GTEx data from the esophagus and the muscle.

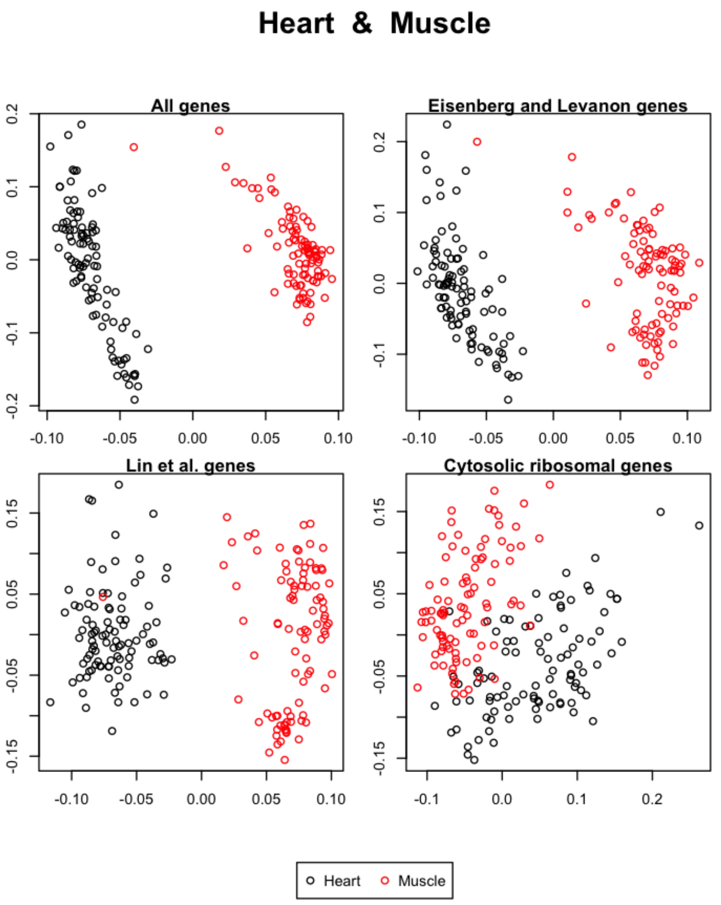


Figure 69: The first and second singular vectors of the GTEx data from the heart and the muscle.

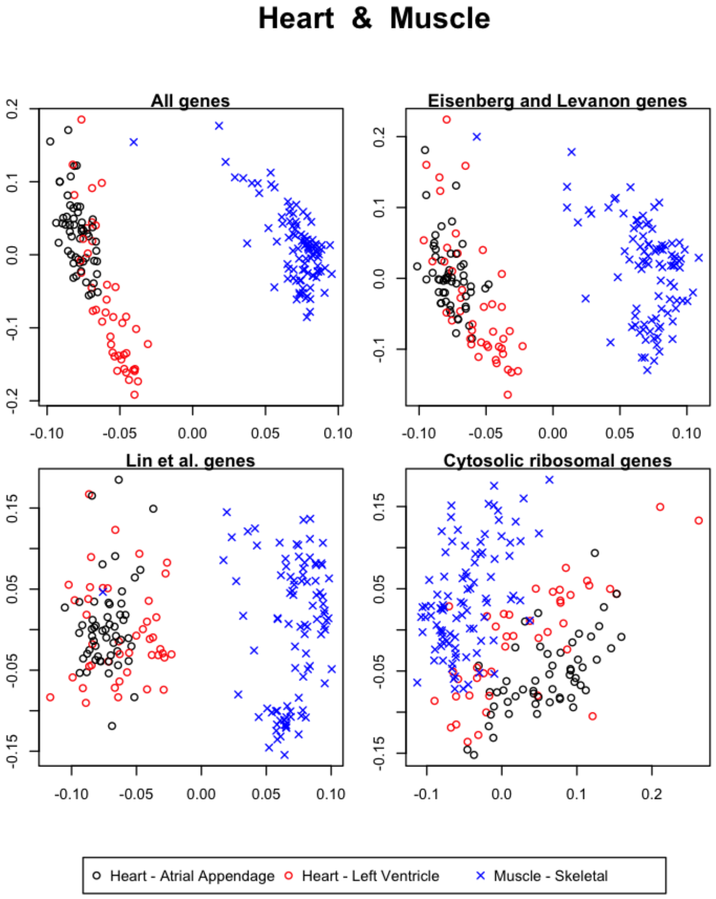


Figure 70: The first and second singular vectors of the GTEx data from the heart and the muscle.

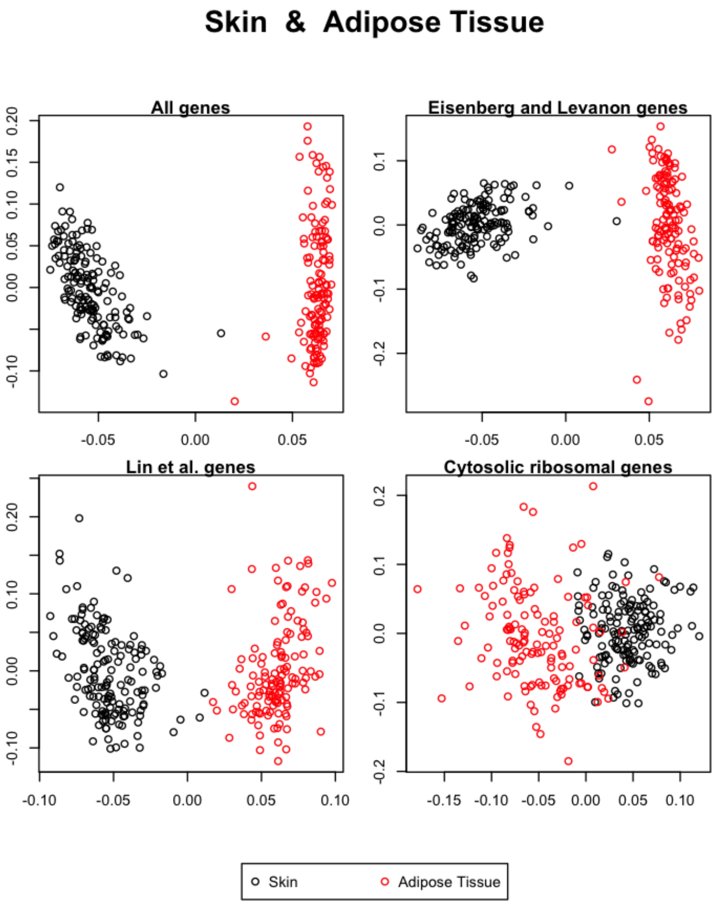


Figure 71: The first and second singular vectors of the GTEx data from the skin and the adipose tissue.

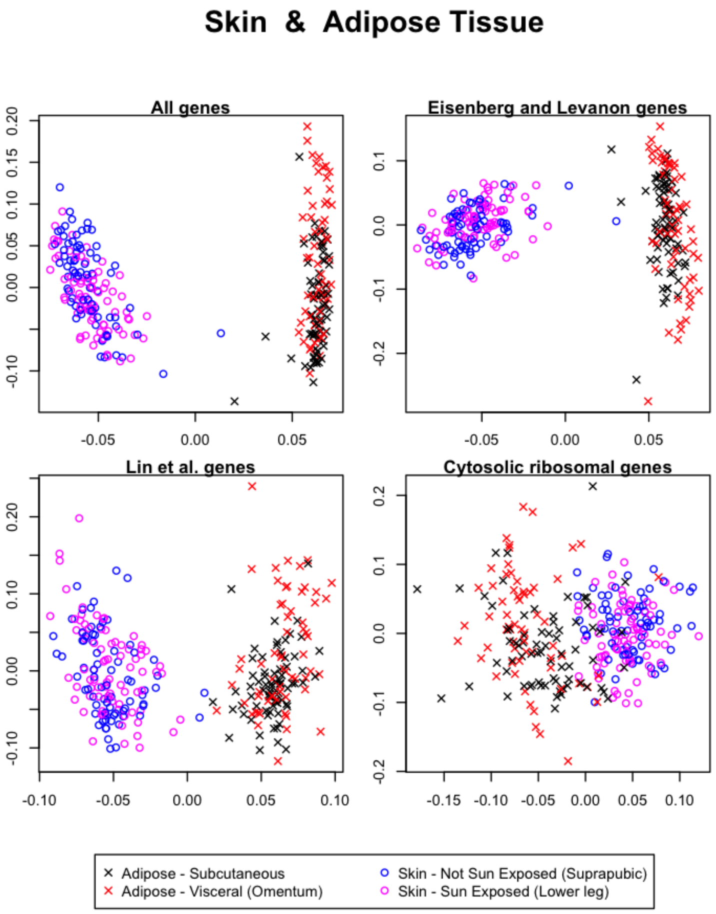


Figure 72: The first and second singular vectors of the GTEx data from the skin and the adipose tissue.

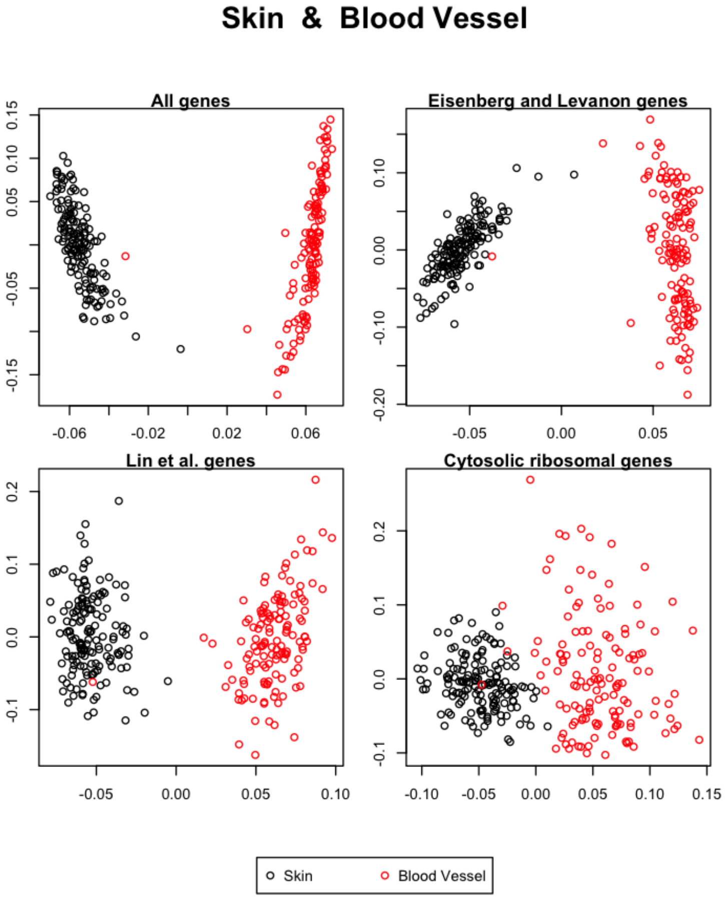


Figure 73: The first and second singular vectors of the GTEx data from the skin and the blood vessel.

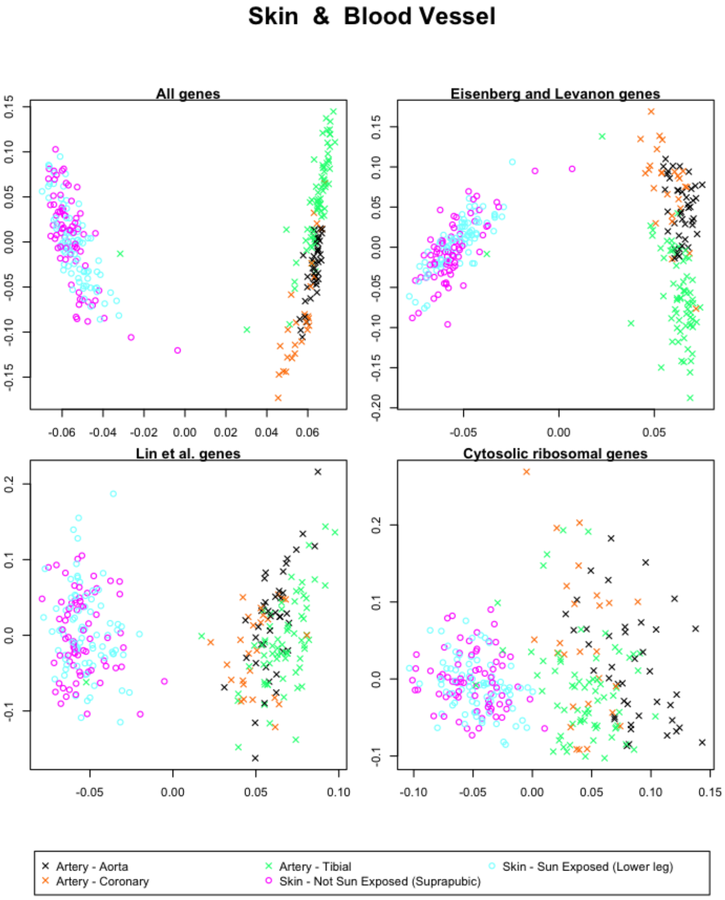


Figure 74: The first and second singular vectors of the GTEx data from the skin and the blood vessel.

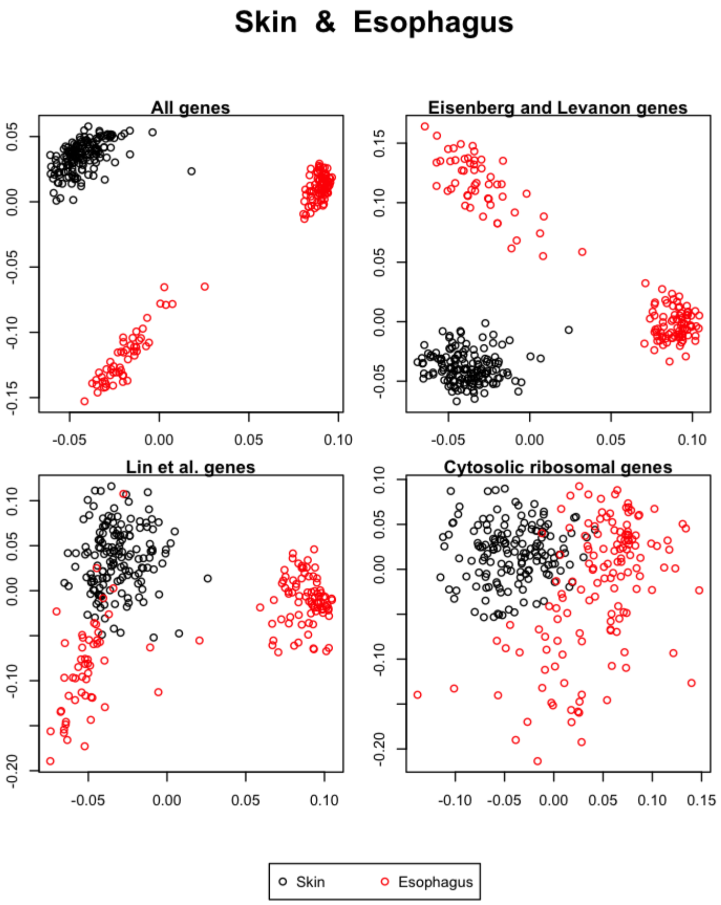


Figure 75: The first and second singular vectors of the GTEx data from the skin and the esophagus.

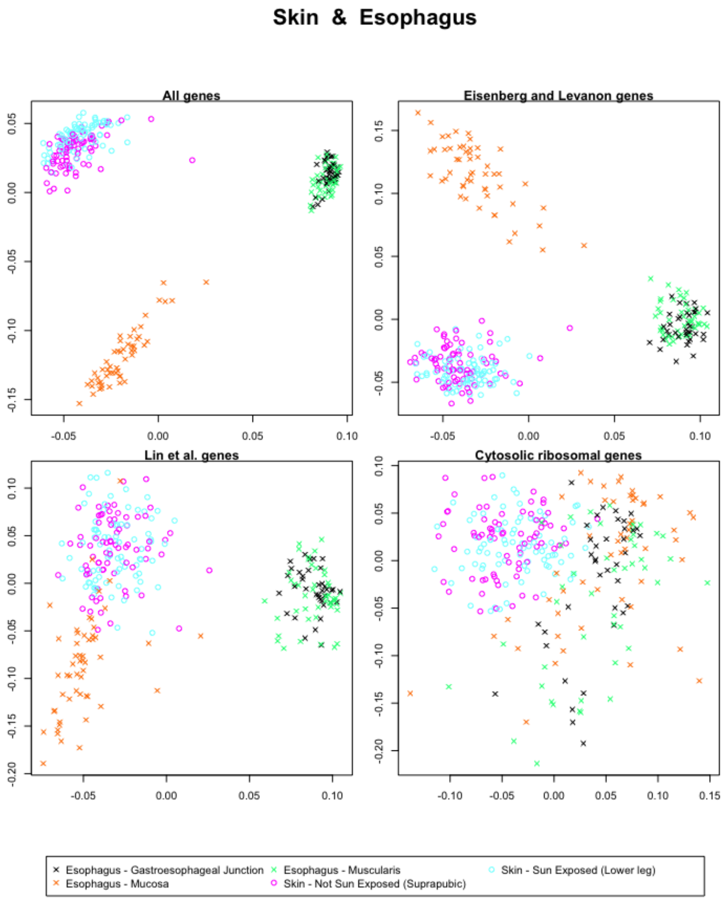


Figure 76: The first and second singular vectors of the GTEx data from the skin and the esophagus.

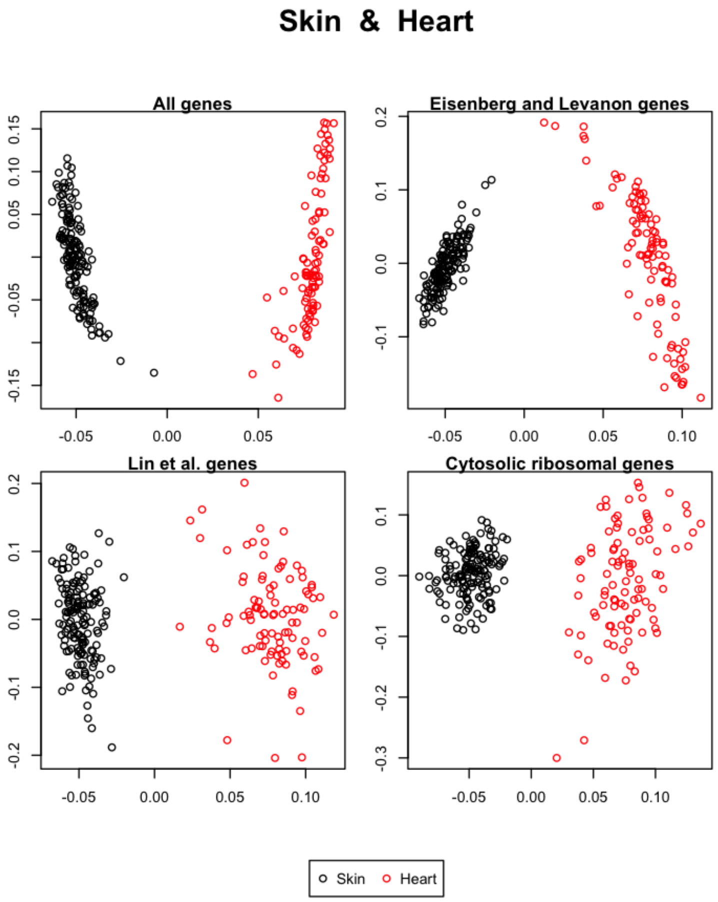


Figure 77: The first and second singular vectors of the GTEx data from the skin and the heart.

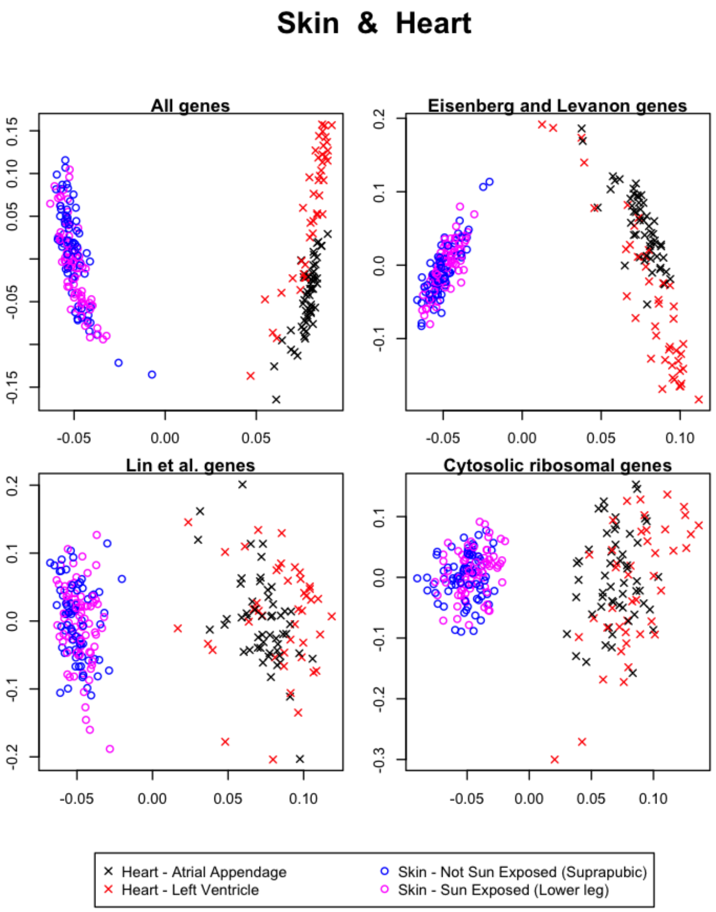


Figure 78: The first and second singular vectors of the GTEx data from the skin and the heart.

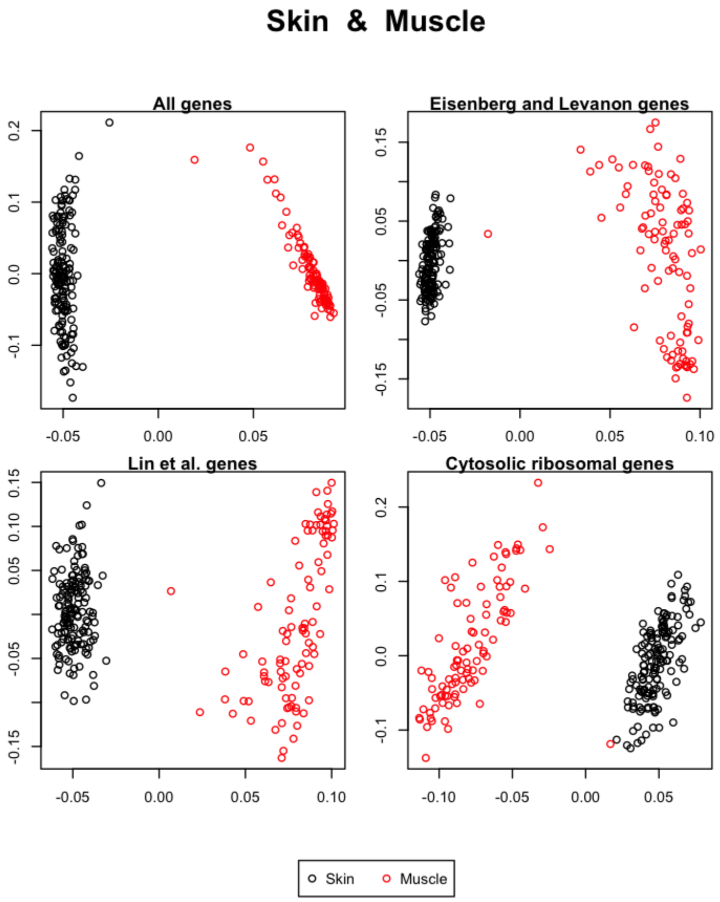


Figure 79: The first and second singular vectors of the GTEx data from the skin and the muscle.

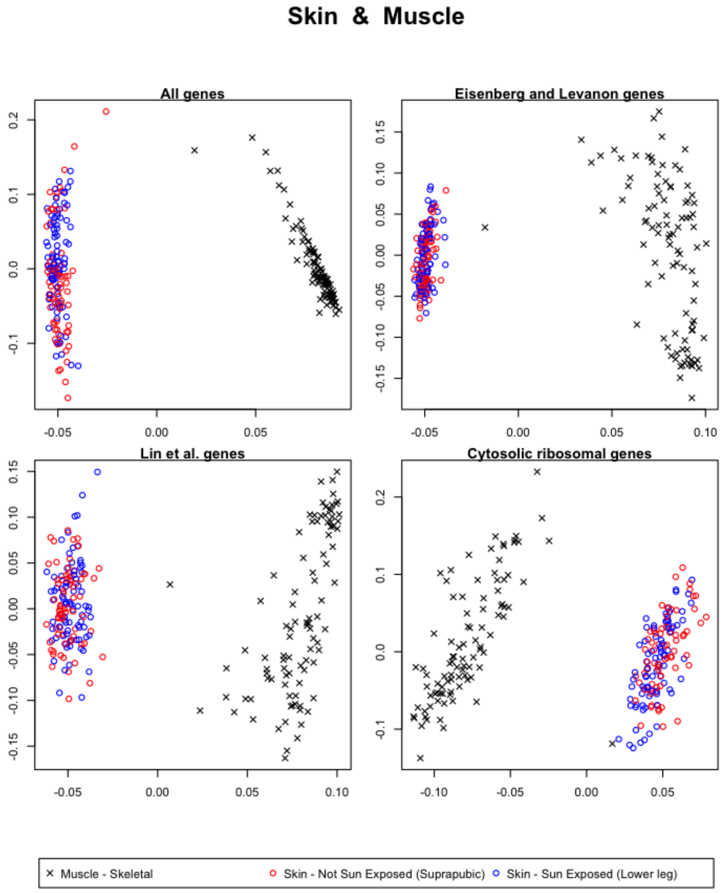


Figure 80: The first and second singular vectors of the GTEx data from the skin and the muscle.

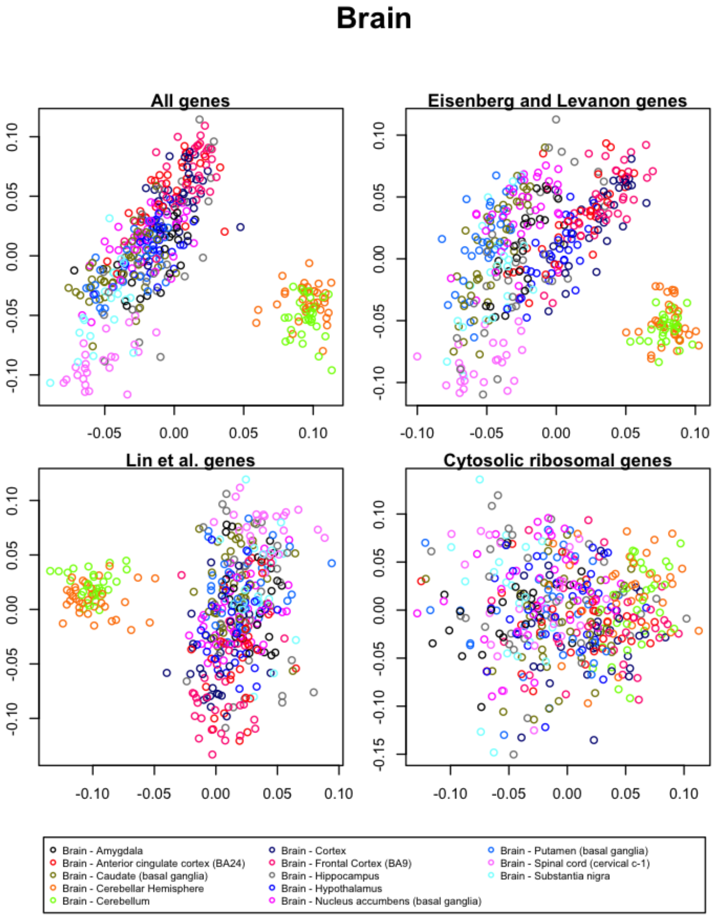


Figure 81: The first and second singular vectors of the GTEx data from the brain.

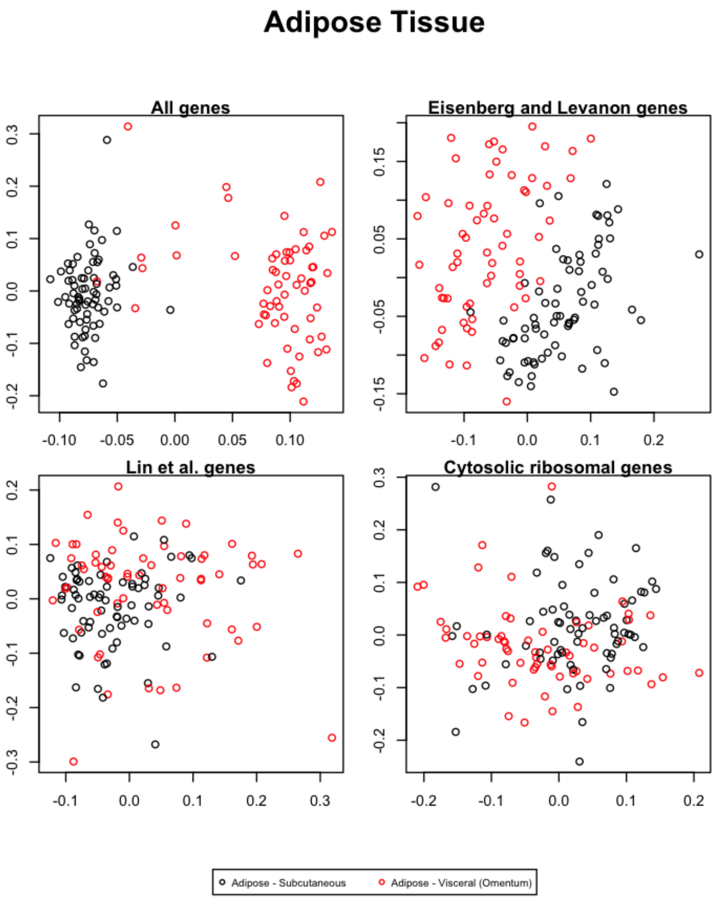


Figure 82: The first and second singular vectors of the GTEx data from the adipose tissue.

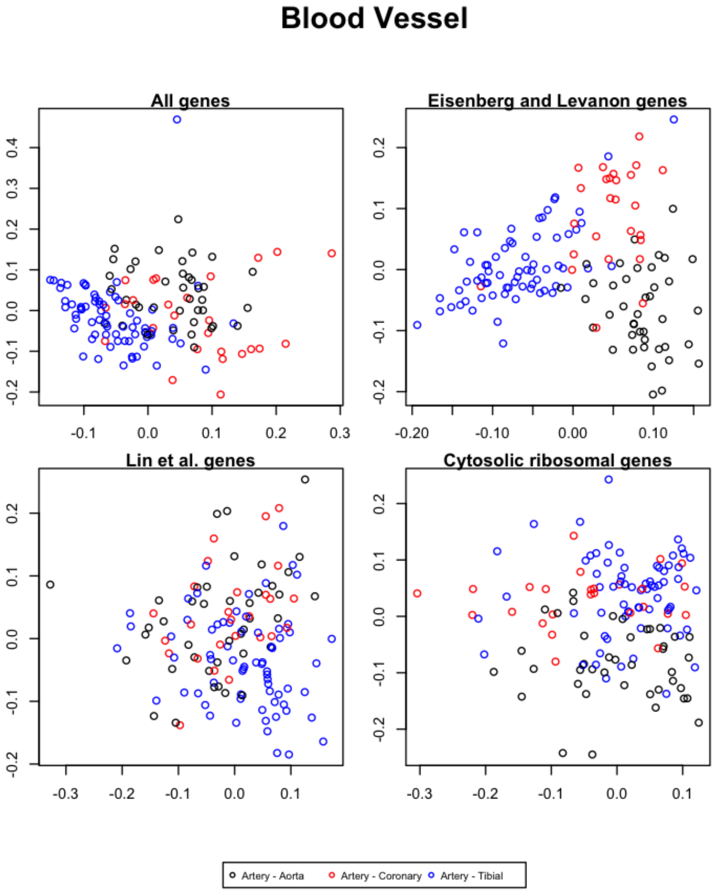


Figure 83: The first and second singular vectors of the GTEx data from the blood vessels.

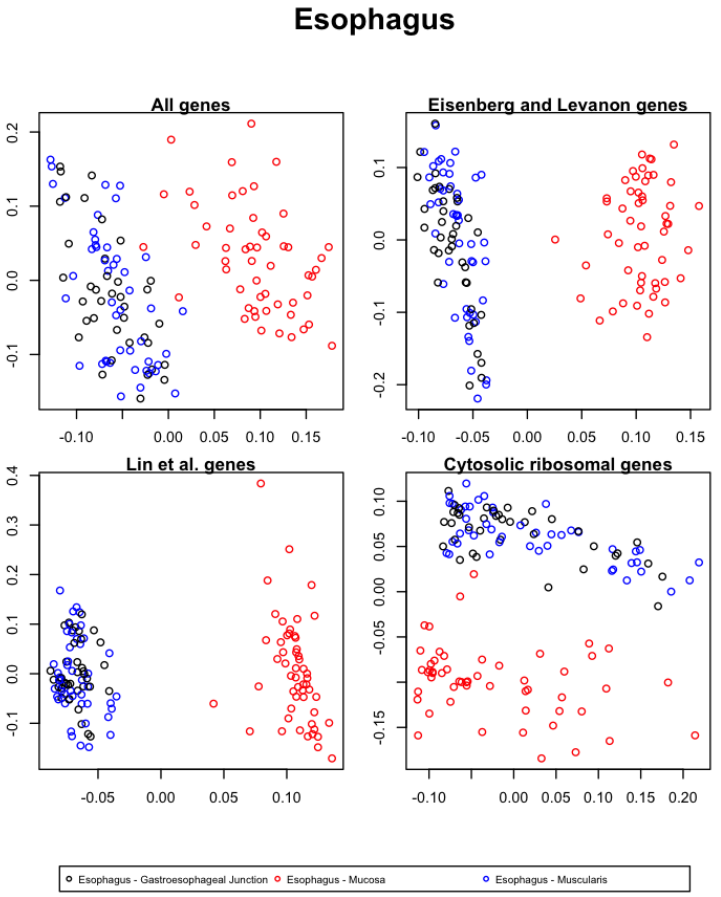


Figure 84: The first and second singular vectors of the GTEx data from the esophagus.

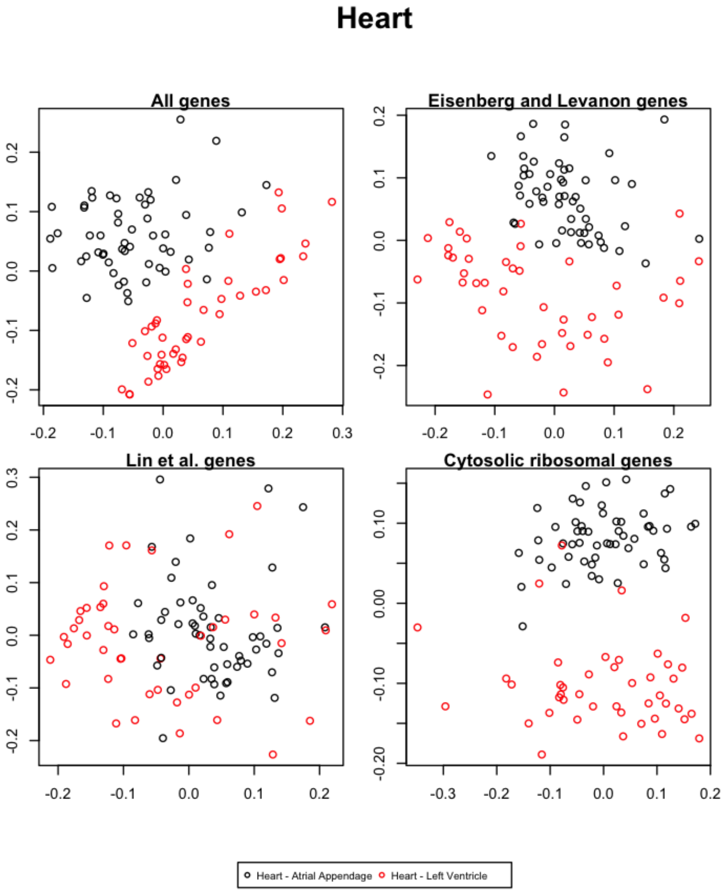


Figure 85: The first and second singular vectors of the GTEx data from the heart.

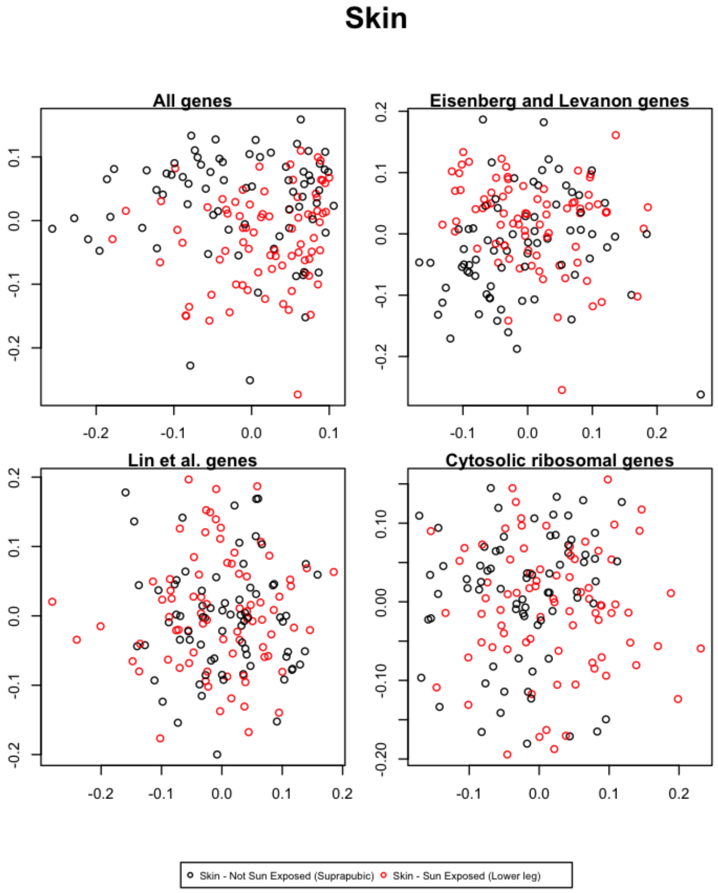


Figure 86: The first and second singular vectors of the GTEx data from the skin.

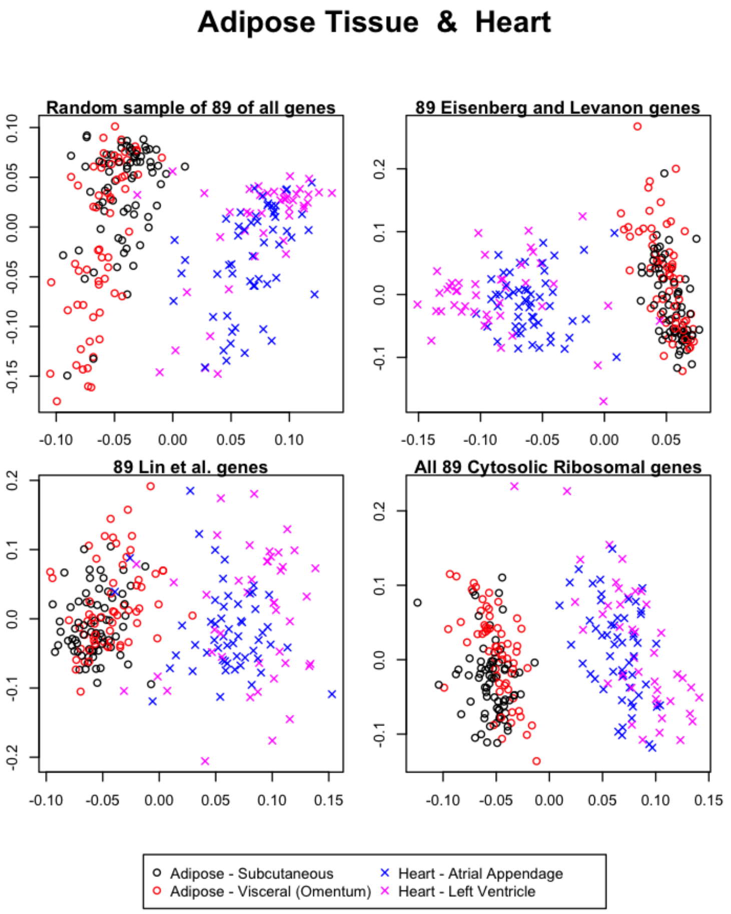


Figure 87: The first and second singular vectors of the GTEx data from the adipose tissue and the heart.

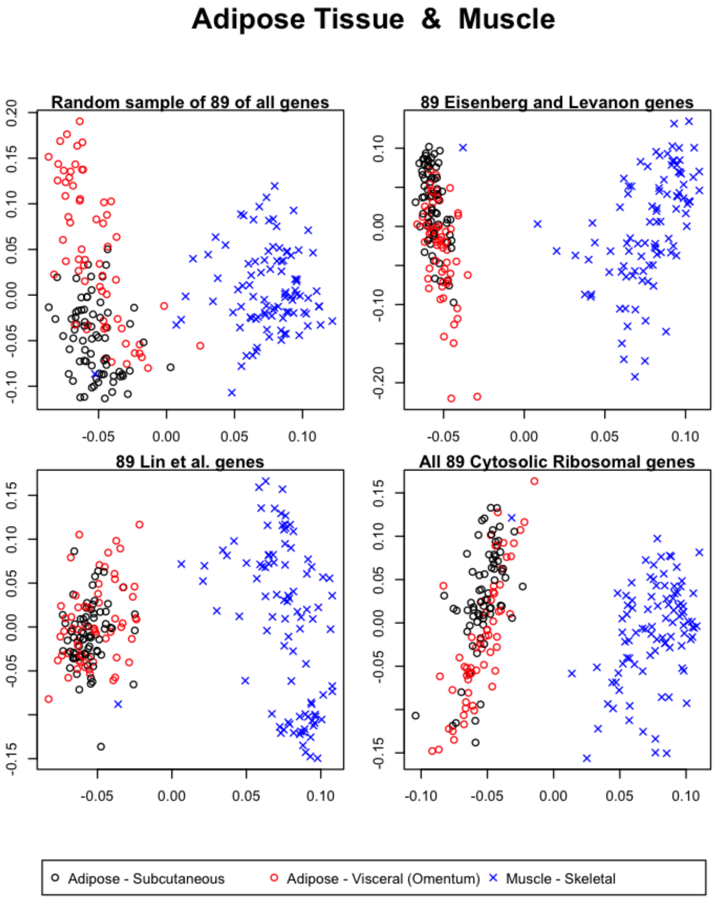


Figure 88: The first and second singular vectors of the GTEx data from the adipose tissue and the muscle.

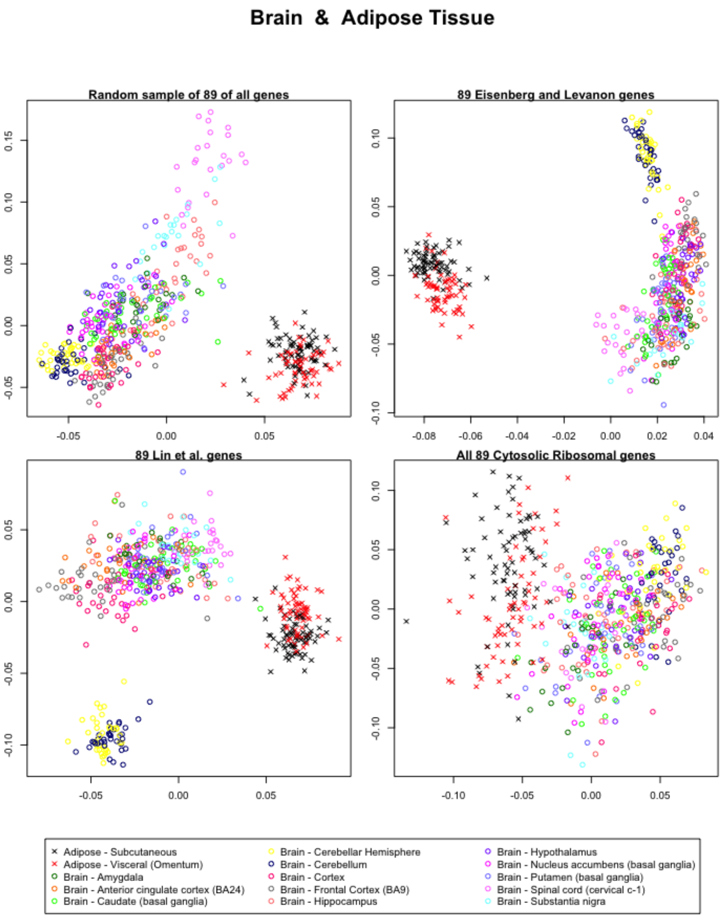


Figure 89: The first and second singular vectors of the GTEx data from the brain and the adipose tissue.

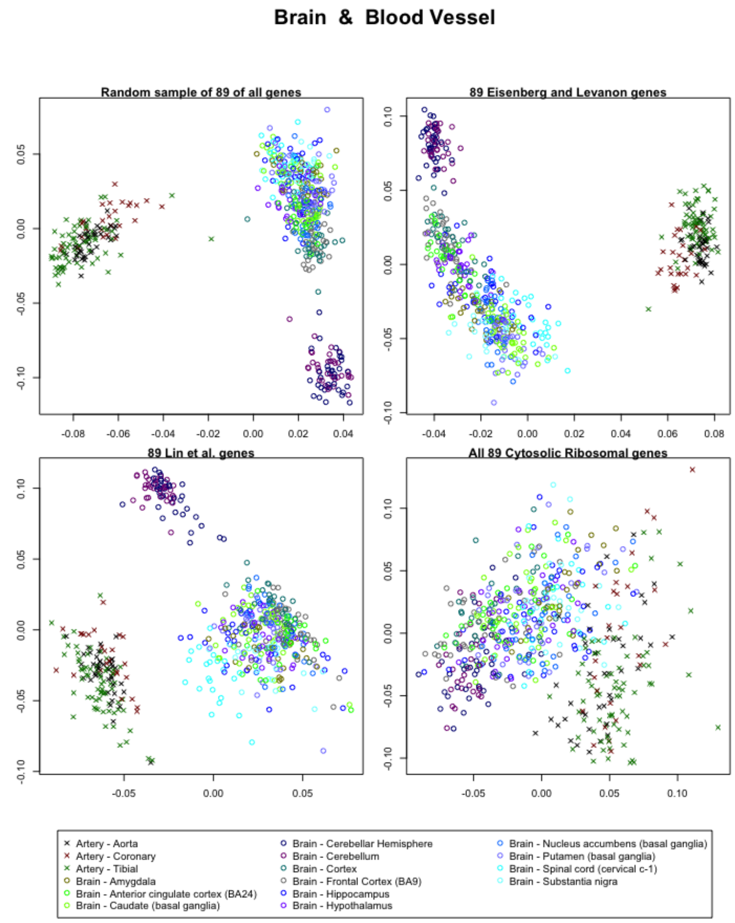


Figure 90: The first and second singular vectors of the GTEx data from the brain and the blood vessel.

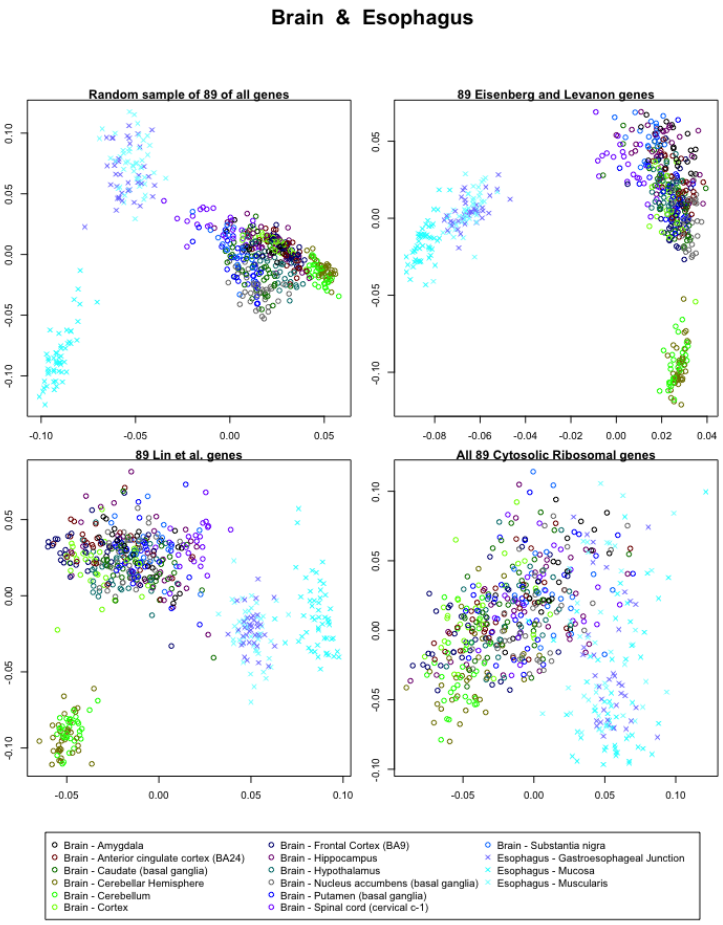


Figure 91: The first and second singular vectors of the GTEx data from the brain and the esophagus.

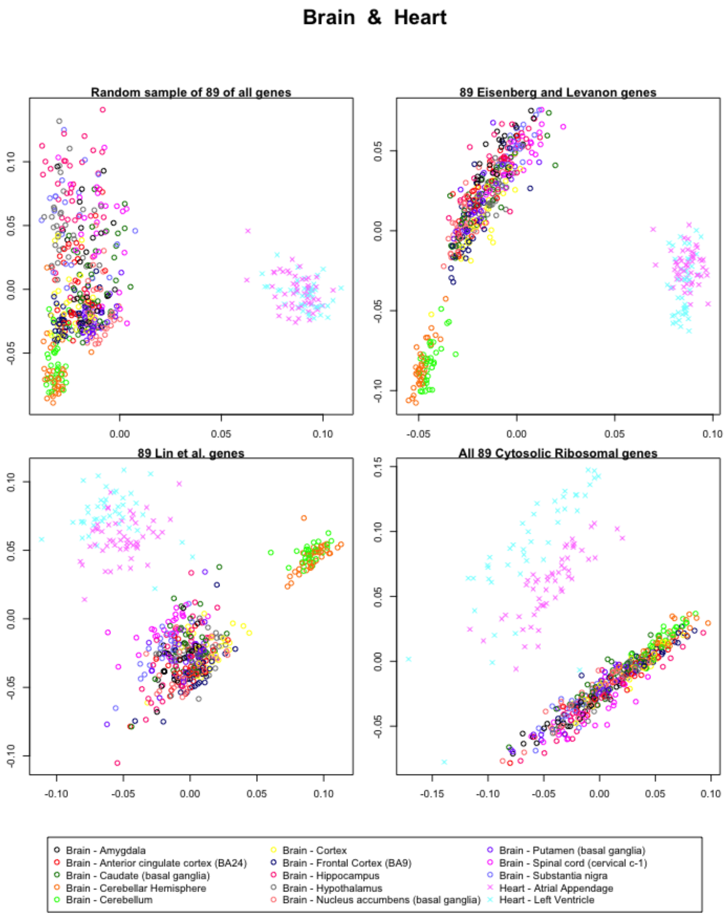


Figure 92: The first and second singular vectors of the GTEx data from the brain and the heart.

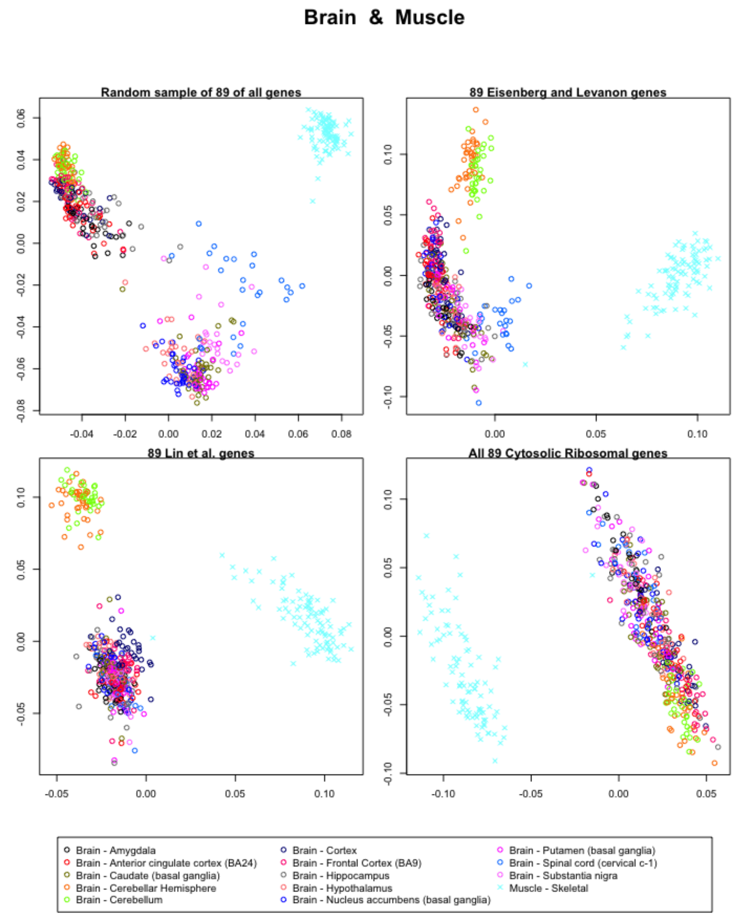


Figure 93: The first and second singular vectors of the GTEx data from the brain and the muscle.

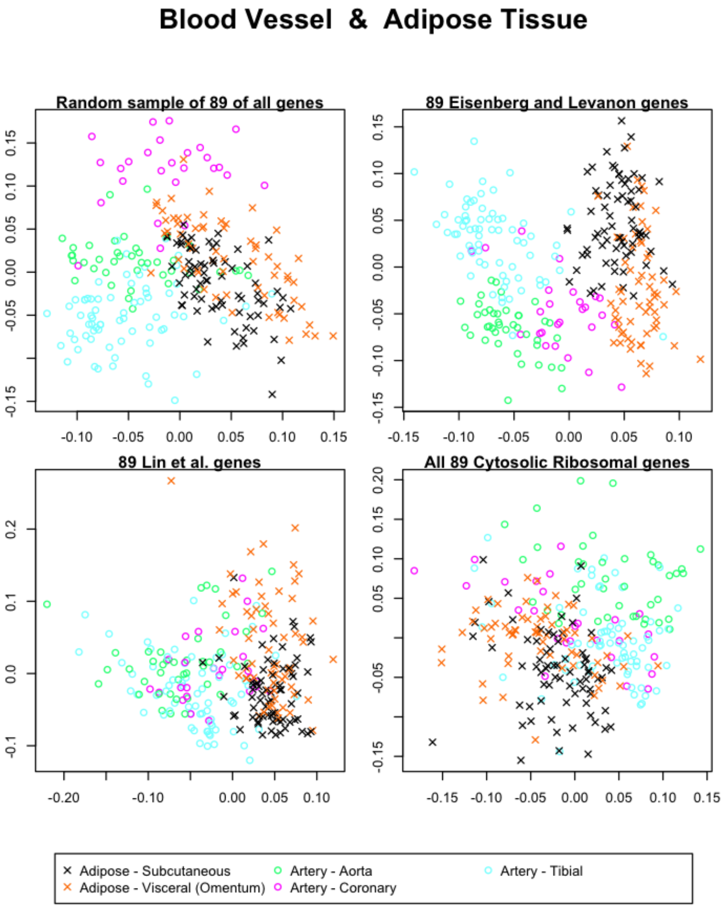


Figure 94: The first and second singular vectors of the GTEx data from the blood vessel and the adipose tissue.

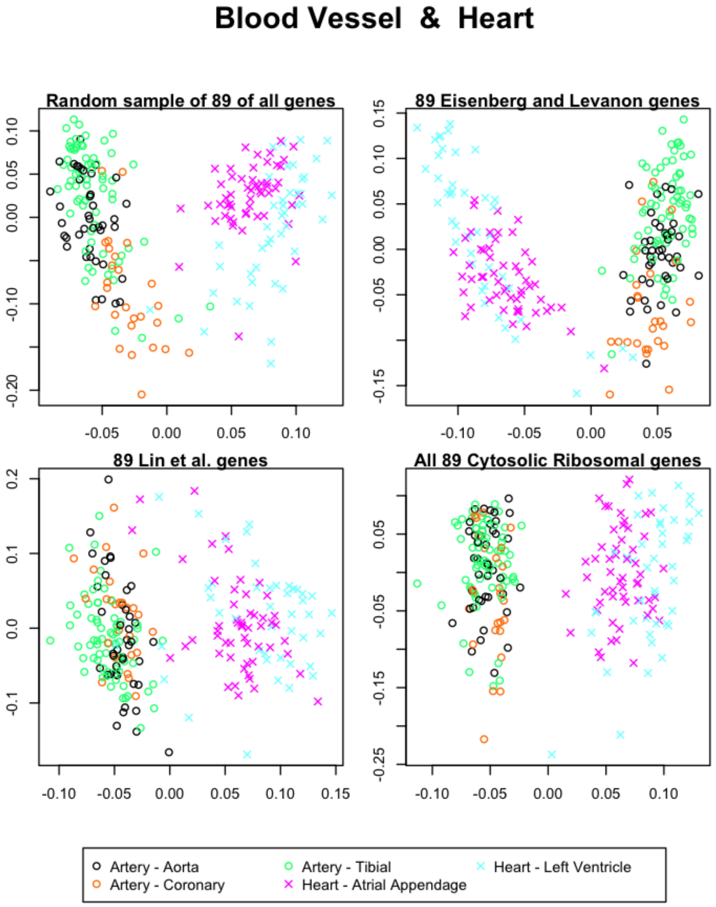


Figure 95: The first and second singular vectors of the GTEx data from the blood vessel and the heart.

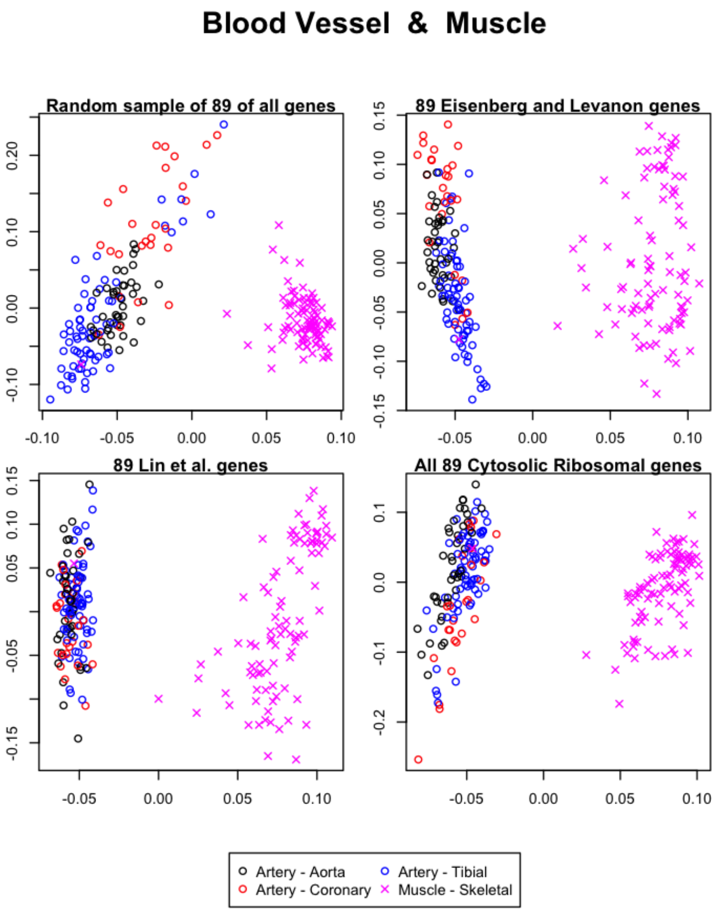


Figure 96: The first and second singular vectors of the GTEx data from the blood vessel and the muscle.

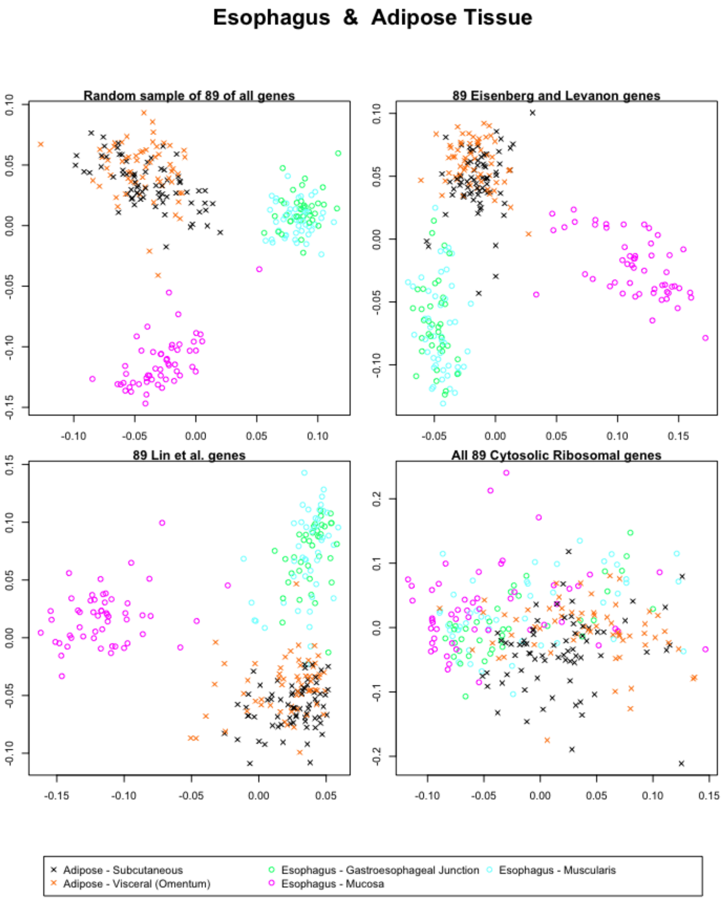


Figure 97: The first and second singular vectors of the GTEx data from the esophagus and the adipose tissue.

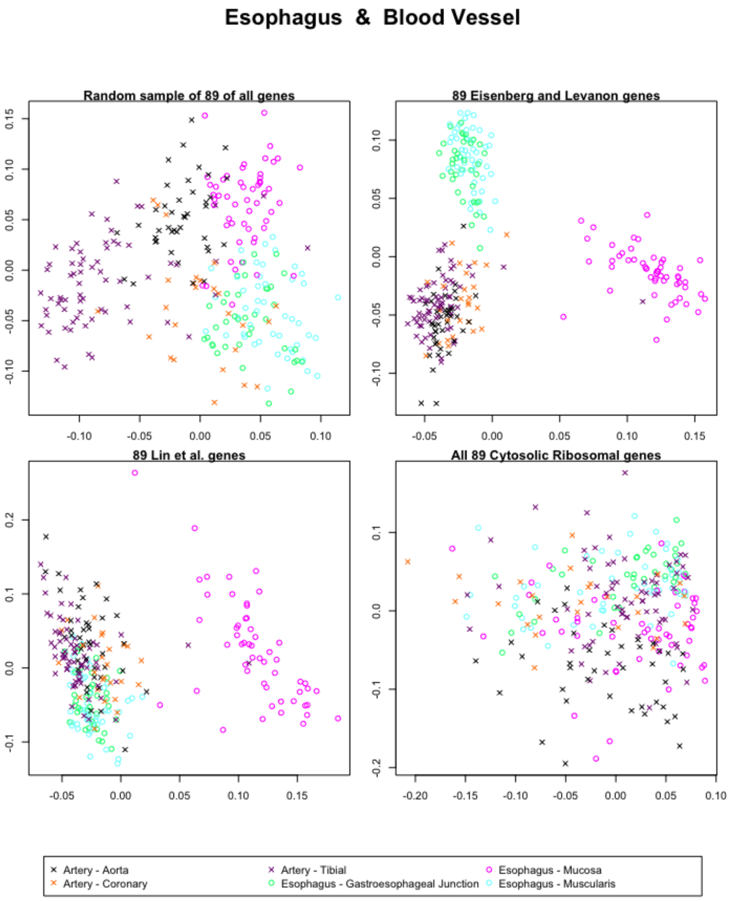


Figure 98: The first and second singular vectors of the GTEx data from the esophagus and the blood vessel.

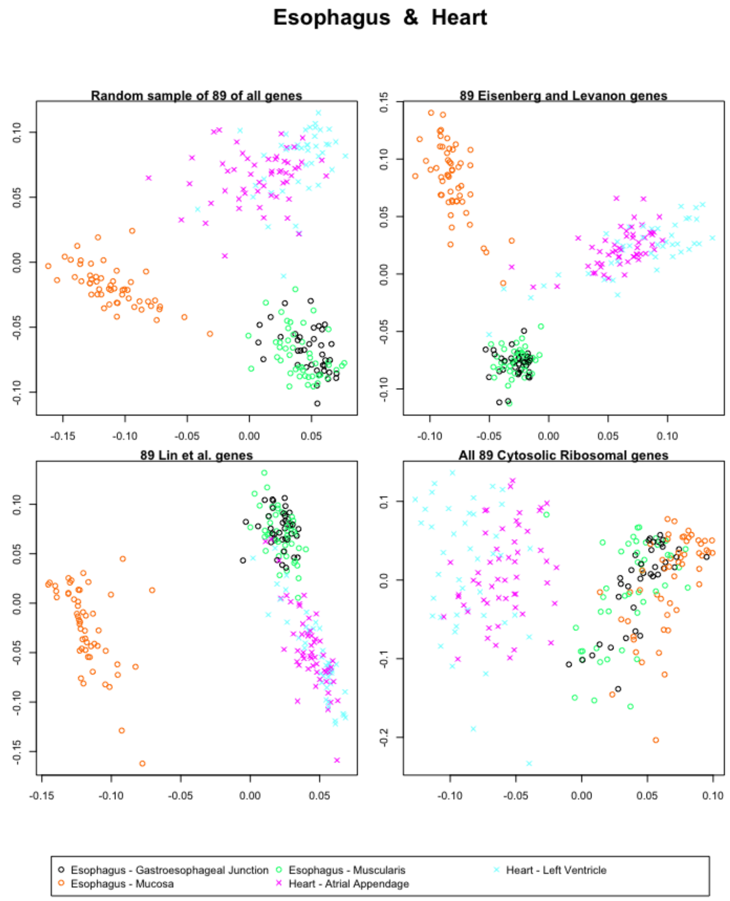


Figure 99: The first and second singular vectors of the GTEx data from the esophagus and the heart.

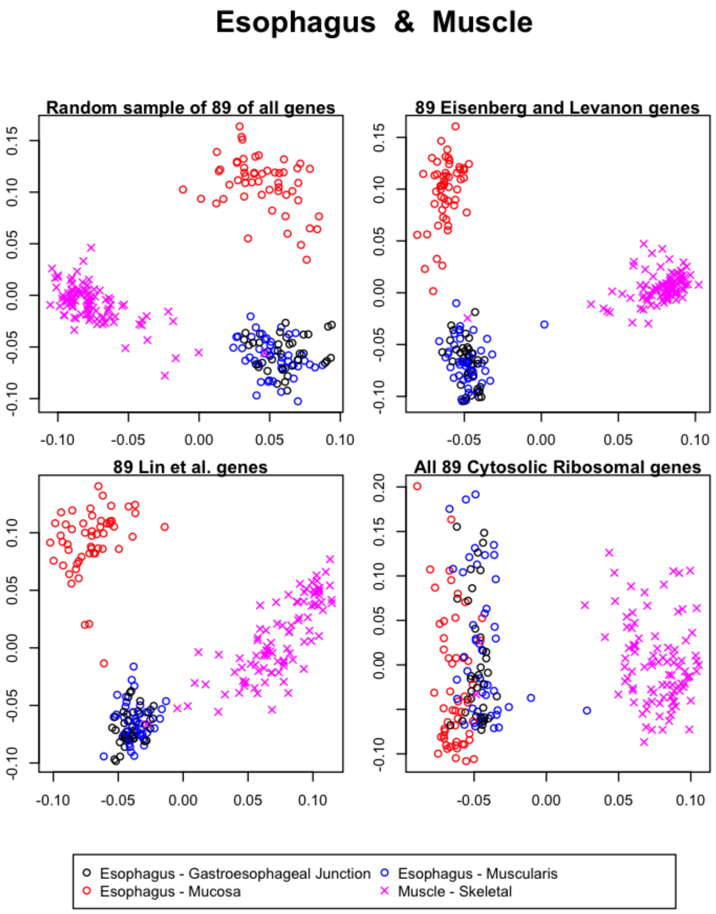


Figure 100: The first and second singular vectors of the GTEx data from the esophagus and the muscle.

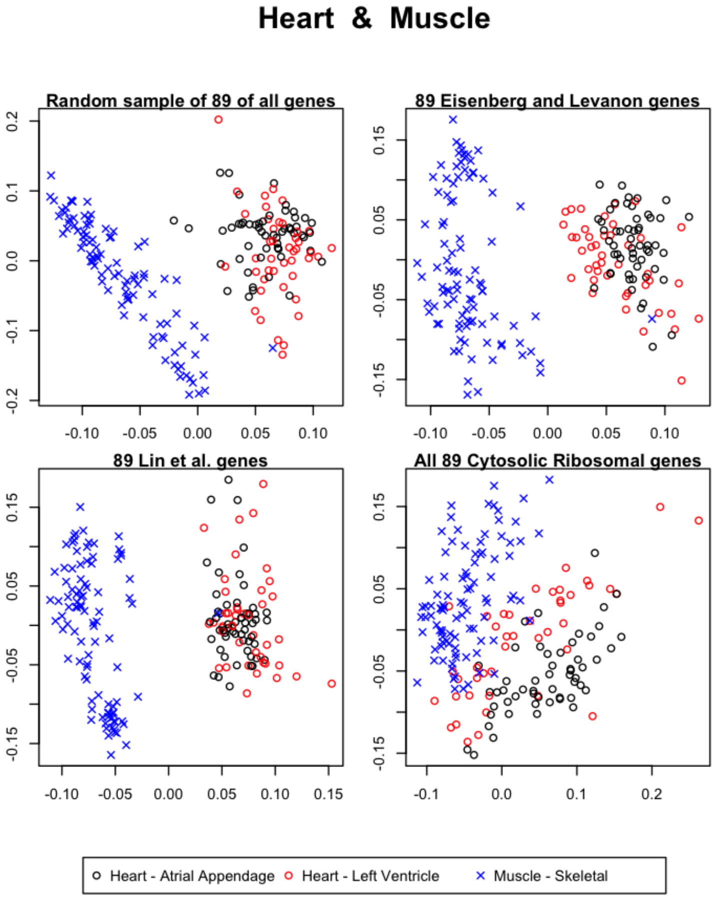


Figure 101: The first and second singular vectors of the GTEx data from the heart and the muscle.

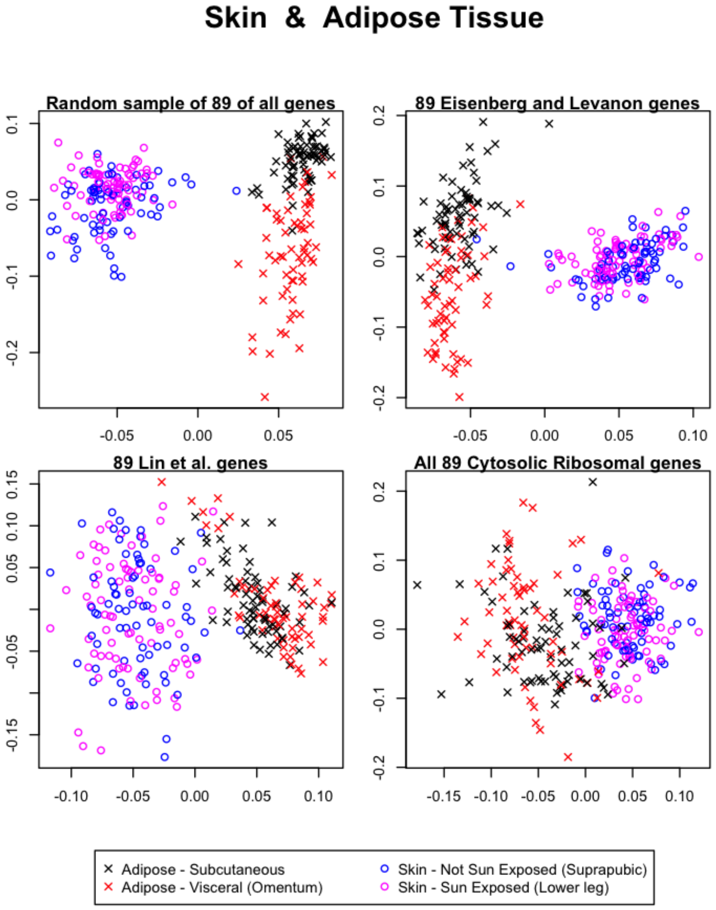


Figure 102: The first and second singular vectors of the GTEx data from the skin and the adipose tissue.

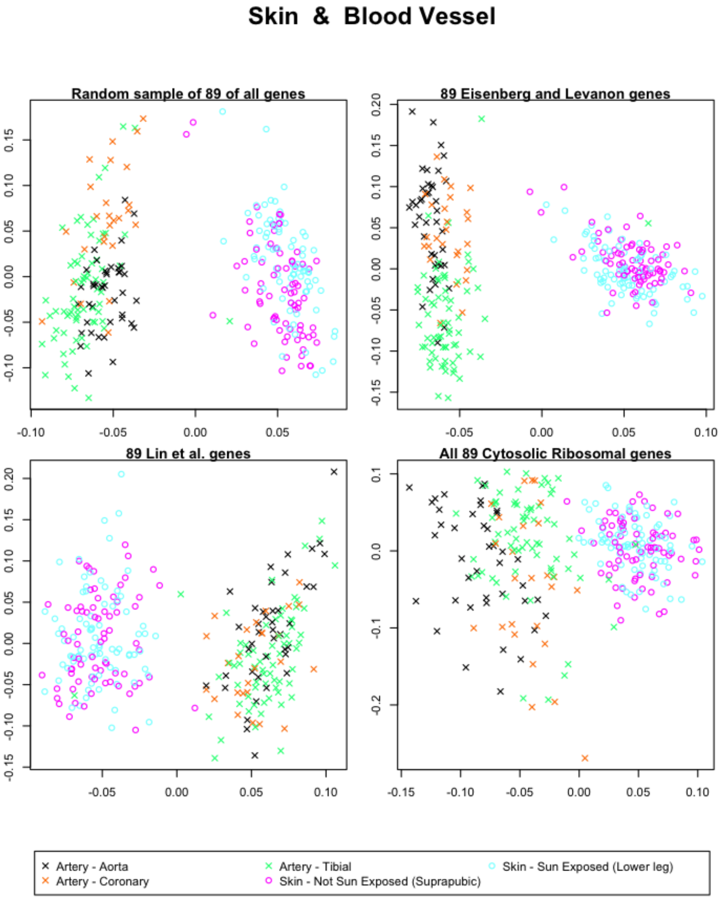


Figure 103: The first and second singular vectors of the GTEx data from the skin and the blood vessel.

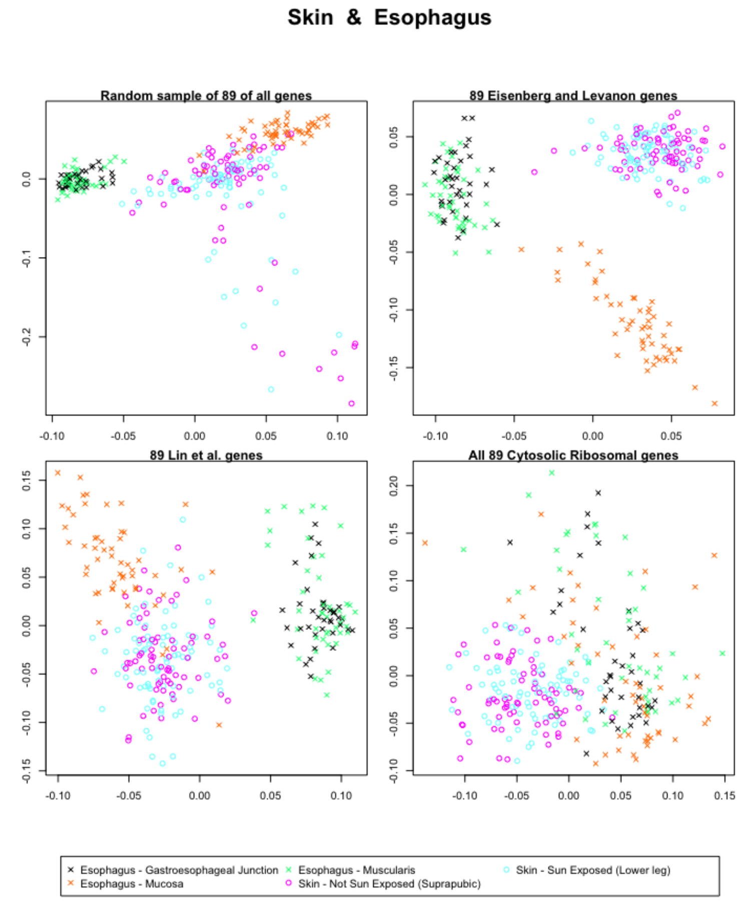


Figure 104: The first and second singular vectors of the GTEx data from the skin and the esophagus.

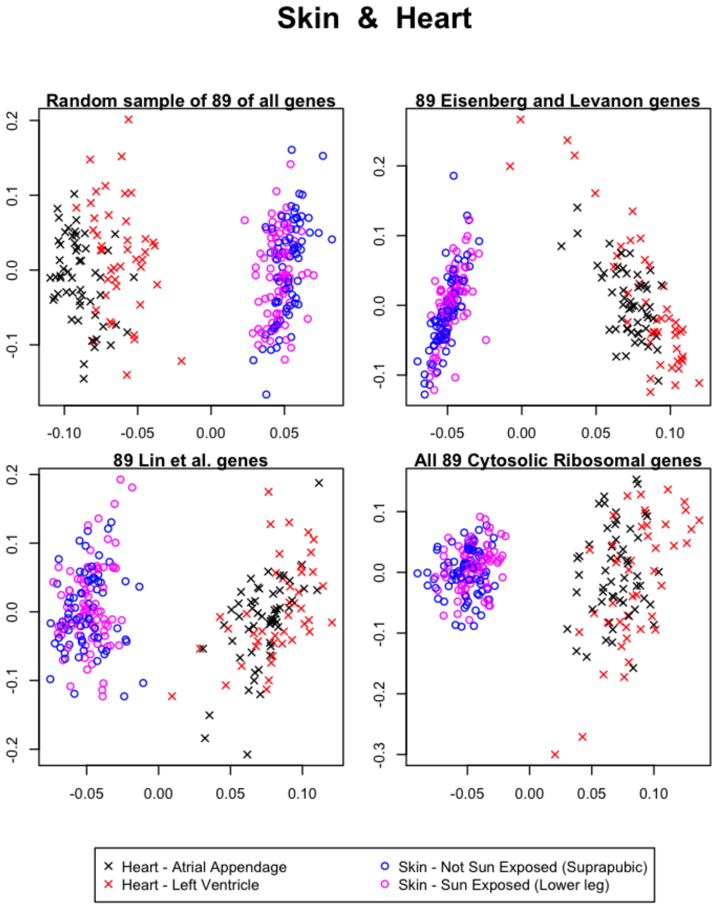


Figure 105: The first and second singular vectors of the GTEx data from the skin and the heart.

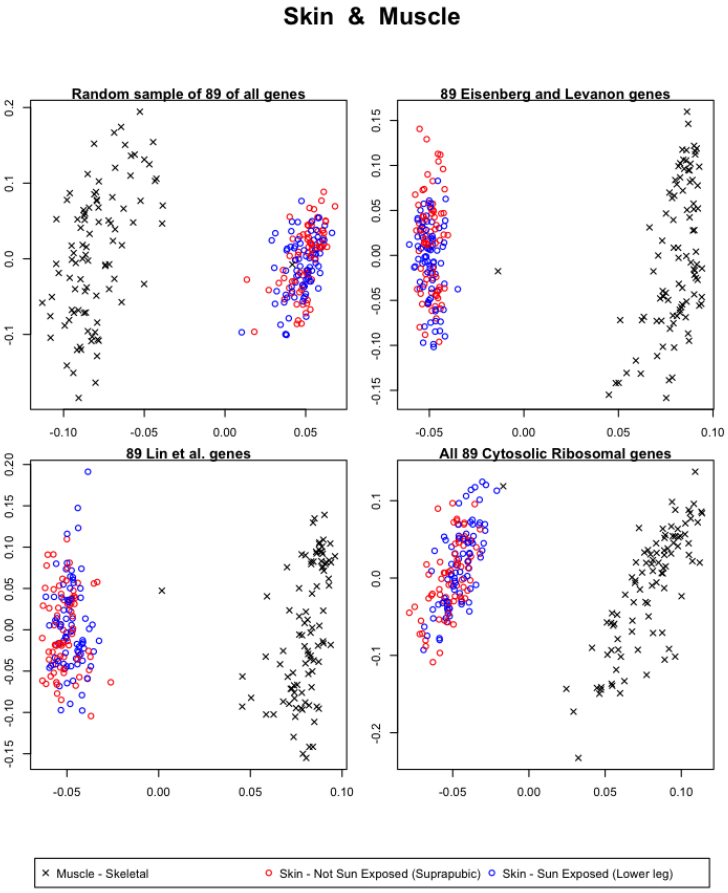


Figure 106: The first and second singular vectors of the GTEx data from the skin and the muscle.
